# Resolving evolutionary relationships in the groundsels: phylogenomics, divergence time estimates, and biogeography of *Packera* (Asteraceae: Senecioneae)

**DOI:** 10.1101/2023.07.18.549592

**Authors:** Erika R. Moore-Pollard, Jennifer R. Mandel

## Abstract

The genus *Packera* belongs to the sunflower family and contains an estimated 64 species and varieties endemic to North America. Some *Packera* are known to hybridize or exhibit polyploidy, making it difficult to reconstruct evolutionary relationships within the group. Previous molecular phylogenetic studies of *Packera* employing ITS data recovered low resolution trees, providing little information on the evolutionary relationships within this complex genus. Therefore, we used next-generation sequencing data to infer nuclear and plastid phylogenies of *Packera* and related Senecioneae taxa. The nuclear phylogeny was calibrated to produce a timetree, then used to reconstruct the macroevolutionary history of *Packera,* including its historical biogeography. We then compared the reconstructed evolutionary history to previously published scenarios based on phylogenetic and geohistorical data. We found that the nuclear and plastid phylogenies were highly incongruent, with the nuclear tree presenting higher resolution than the plastid tree which had an apparent lack of plastid diversity. The nuclear tree indicated that geography may have played a major role in the evolution and taxonomic diversification of *Packera*. The estimated origin of *Packera* at approximately 19.2MY – 25.9MY (late Oligocene to early Miocene) is older than in most other studies. Nonetheless, it aligns well with previous geohistorical predictions, which suggest that speciation and diversification events in *Packera* were driven by changes in geography and climate in North America. Moreover, *Packera* likely originated in the western United States or Mexico, and subsequently diversified north and east into the rest of North America and Russia, in agreement with other studies.

## Introduction

Resolving evolutionary relationships in plant genera that have experienced large amounts of hybridization, polyploidy, and/or diversified rapidly is especially challenging. However, Hyb-Seq approaches have been widely used to shed light on species-level relationships, even in particularly difficult plant groups. For example, *Antennaria* Gaertn. (Asteraceae: Gnaphalieae) is a complicated genus which is known to exhibit extensive hybridization and polyploidy, creating multiple large species complexes (e.g., the *Antennaria rosea* Greene polyploid complex; Bayer, 1989a, 1989b; Thapa et al., 2021). The first molecular phylogeny of *Antennaria* was reconstructed using Internal transcribed spacer (ITS) DNA data, but the trees had little resolution (Bayer et al., 1996). In contrast, implementing a phylogenomic approach in the group yielded a robust phylogeny for *Antennaria*, enabling the testing of biogeographic and character trait hypotheses (Thapa et al., 2020). Similarly, rampant hybridization, coupled with a relatively recent radiation, contributed to the difficulty of developing a robust phylogenetic hypothesis for species of *Heuchera* L. (Saxifragaceae) (Folk & Freudenstein, 2014). However, phylogenomic methods made use of loci spanning intron-exon boundaries to generate highly informative data, ultimately producing a well-resolved phylogeny for the genus (Folk et al., 2015).

The genus *Packera* Á. Löve & D. Löve (Asteraceae: Senecioneae) has a complicated history of hybridization and polyploidy. *Packera* comprises 64 species and varieties (Trock, 2006), though the number is constantly changing with the description of new species and varieties (e.g., Mohlenbrock, 2004; Mahoney & Kowal, 2008; Kowal et al., 2011; Yeatts et al., 2011; Boufford et al., 2014). *Packera* was originally included in the genus *Senecio* L. as an informal group known as the “aureoid *Senecios*” (or Aureoids), first recognized by Asa Gray (Gray 1886, Gray and Torrey 1843, Mahoney 2000). Gray provided the earliest treatments of the group by recognizing that distinct members share most of these following characters: perennial herbs arising from creeping rootstocks or a stout caudex; basal leaves well developed, cauline leaves progressively reduced upward, leaf margins without callose denticles; roots fibrous, thin and branching; and haploid chromosome numbers of 22 or 23 (Barkley, 1988). *Packera* was then formally recognized as a distinct genus in 1976 (Löve & Löve, 1976; Weber & Löve, 1981; Jeffrey, 1992). Except for one species known from Siberia (*P. heterophylla* (Fisch.) (E.Wiebe), *Packera* is endemic to North America, with most species occurring in temperate western North America (Barkley, 1988). *Packera* is adapted to a variety of habitats, with some species that are abundant and widely distributed, and others that are narrow endemics, restricted to specialized or geographically isolated habitats (Fig. 1). The taxonomy of *Packera* remains complex due to geohistorical events during its evolution (Barkley, 1988), rampant hybridization documented in the wild, and a high incidence of polyploidy; roughly 40% of taxa exhibit polyploidy, aneuploidy, or other cytological disturbances (Trock, 2006).

**Fig. 1.**
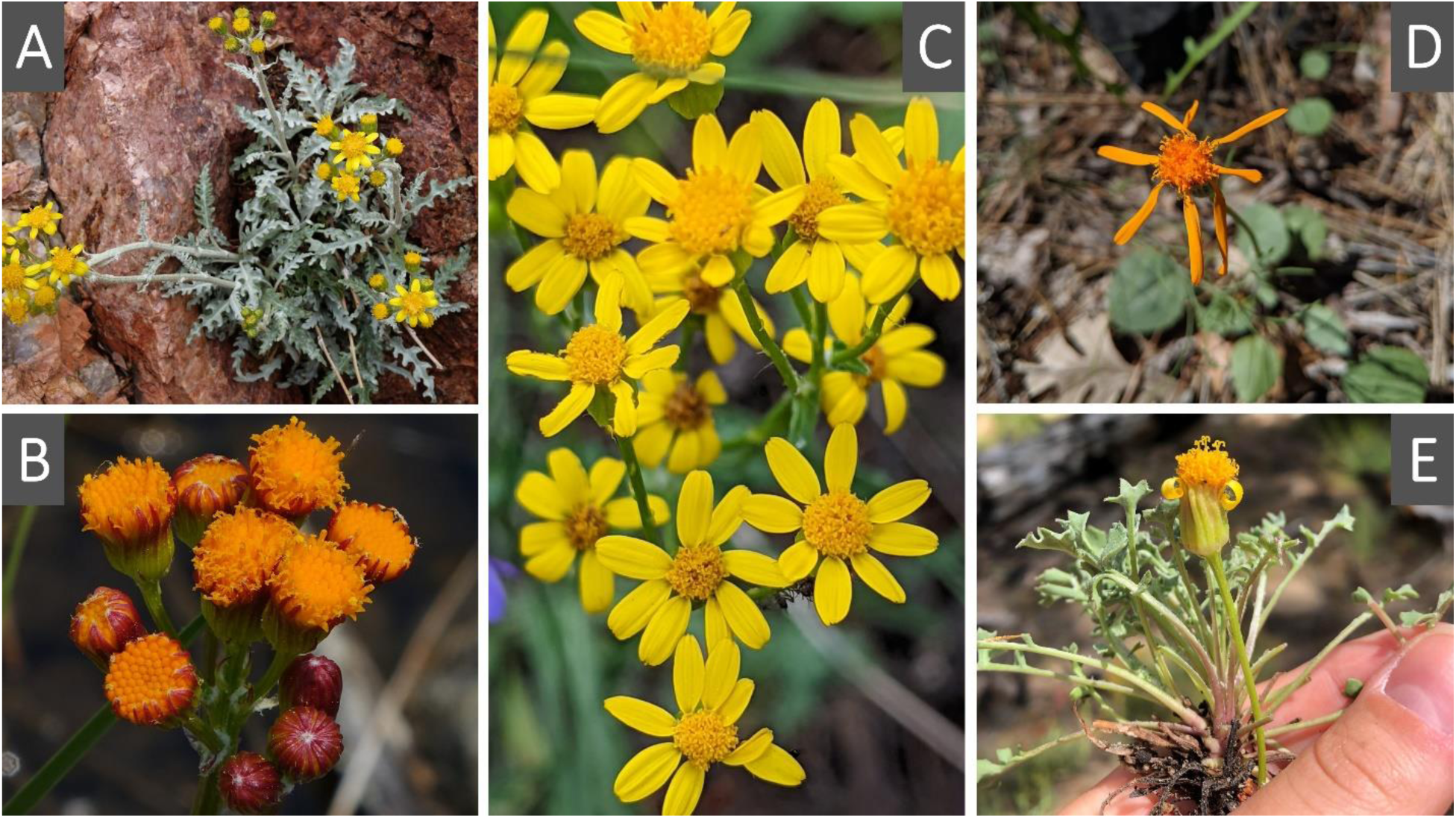
Images of *Packera* species highlighting some of the morphological diversity seen in the group. **A.** Fendler’s ragwort, *Packera fendleri* (A.Gray) W.A.Weber & Á.Löve; **B.** Weak groundsel, *P. debilis* (Nutt.) W.A.Weber & Á.Löve; **C**. Lobeleaf groundsel, *P. multilobata* (Torr. & A.Gray ex A.Gray) W.A.Weber & Á.Löve; **D**. Flame ragwort, *P. greenei* (A.Gray) W.A.Weber & Á.Löve; **E.** Tehachapi ragwort, *P. ionophylla* (Greene) W.A.Weber & Á.Löve. Image credits: **A**-**B**: Robert Lagier; **C**: Lucas C. Wheeler; **D**-**E**: Erika R. Moore-Pollard.

Numerous attempts have been made to better understand species-level relationships within *Packera* using molecular phylogenetic analyses of ITS DNA sequence data (Bain & Jansen, 1995; Bain & Golden, 2000; Schilling & Floden, 2015); however, these studies were unable to resolve species-level relationships in *Packera* due to a lack of phylogenetic signal in the ITS data, as ITS could only verify identifications of members of *Packera* to genus level (Schilling & Floden, 2015). The most recent phylogeny of *Packera* based on nuclear ITS sequences was published in 2000 and sampled 27 *Packera* species and 23 total outgroup taxa (Bain & Golden, 2000). The phylogeny had many unresolved areas (i.e., polytomies) possibly due to the existence of few phylogenetically informative sites in the DNA sequence alignment, sampling of only a few species, or reticulate evolution. Despite these challenges, the phylogeny indicated (1) that *Packera* is monophyletic, although weakly supported, (2) the Mexican, Gulf, and West Coast species are most likely the deepest branching lineages of the genus, (3) *Packera*’s closest relatives are “Old World” members of subtribe Senecioninae, and (4) there appears to be a relationship between geographic distribution and habitat (Bain & Golden, 2000).

To date, no phylogenetic studies have evaluated plastid variation across the genus *Packera*. Doing so would be beneficial since some studies involving *Packera* species have observed more plastid diversity than expected (Bain & Jansen, 1996; Golden & Bain, 2000; Bain & Golden, 2005). Additionally, the timing of diversifications in *Packera* has never been specifically investigated, although there is speculation that *Packera* evolved during the later stages of the mid Tertiary (25-40MY; Barkley, 1988; Bain & Golden, 2000). However, recent diversification rate analyses involving Asteraceae and Senecioneae recover an estimated age of 7-20MY for *Packera* (Pelser et al., 2010; Huang et al., 2016; Mandel et al., 2019; Zhang et al., 2021), considerably younger than predictions based on geohistorical interpretations.

To gain a better understanding of the evolutionary relationships and timing of origin of major clades within this complicated genus, we undertook phylogenetic analyses on target enrichment data obtained from 83 *Packera* and 23 outgroup taxa. Our study reconstructed phylogenomic relationships within *Packera* and estimated the timing of divergences within the genus using both target enrichment sequence data from conserved nuclear loci and by-catch plastid sequence data. Using our resulting phylogenetic hypothesis, we asked: (1) is *Packera*, as currently circumscribed, monophyletic, (2) are plastid and nuclear phylogenies congruent, (3) do the resulting species relationships agree with previous interpretations of *Packera*’s geohistory, (4) when did *Packera* originate, and (5) do biogeographic patterns agree with previous interpretations of *Packera*’s evolutionary history?

## 2 Materials and methods

### 2.1 Specimen collection

A nearly complete sampling of recognized *Packera* species, varieties, and accepted hybrids were used in this study. *Packera* species were selected given information published in International Plant Names Index (IPNI; available at http://www.ipni.org) and then edited from discussions with botanists that have previously specialized in *Packera* (D. Trock & J. Bain, pers. comm.). Outgroup taxa were selected by consulting recent studies of the group (Bain & Golden, 2000; Pelser et al., 2010; Mandel et al., 2019). A complete list of sampled species, herbarium vouchers, and publication status can be found in Supplemental Table 1.

### 2.2 DNA extraction and sequencing

Leaf tissue was obtained from herbarium sheets or live plants that were dried in silica beads. Dried leaf tissue (∼25mg) was ground using a Bead Mill 24 Homogenizer (ThermoFisher Scientific, Atlanta, Georgia, USA), and total DNA was extracted using the Omega Bio-Tek Kit (Atlanta, Georgia, USA) following manufacturer’s protocols with the addition of polyvinylpyrrolidone (PVP) and ascorbic acid to the first extraction buffer (10mL SQ1 buffer, 100mg PVP, 90mg ascorbic acid). Samples of *Packera* and outgroup species that were obtained from other researchers followed different DNA extraction protocols which are outlined in Supplemental Table 1. Using DNA from various DNA extraction methods should not influence the results (Healey et al., 2014). All DNA samples used in this study were quality checked using a Nanodrop 2000 (Thermo Fisher Scientific, Carlsbad, California, USA) and quantified using a Qubit High Sensitivity (HS) assay (ThermoFisher Scientific, Oregon, USA). If the DNA had low concentrations, or appeared to be of low-quality, the samples were cleaned following the E.Z.N.A. Cycle Pure Kit (Omega Biotek, Georgia, USA) and were re-quantified with the Qubit HS assay. Up to 1µg DNA per sample was sonicated with a QSonica machine (ThermoCube, New York, USA) to generate fragment sizes within the 400-500 base pair (bp) range. The amount of DNA used is dependent on initial total DNA concentration of the targeted sample. Time allotted for sonication of each sample varied depending on approximate length and degree of fragmentation of the DNA. Prior to sonication, total DNA was visualized on a 1% agarose gel in 1X TBE and GelRed 3x (Biotium) and the approximate length was estimated by comparing to a 100bp GeneRuler DNA ladder (ThermoFisher Scientific, Vilnius, Lithuania). If the sample of DNA appeared smaller than 400bp, the samples were not sonicated.

Libraries for each sample were prepared using the NEBNext Ultra II DNA Library Prep Kit and NEBNext Muliplex Oligos for Illumina Kits (New England Biolabs, Ipswich, Massachusetts, USA) following the NEBNext Ultra II Version 5 protocol with size selection on DNA fragments at 300-400bp range. To make library preparation more economic, the protocol was adjusted by halving all amounts of reagents and DNA used. The barcoded libraries (NEBNext multiplex oligos) were quality checked with an Agilent Bioanalyzer (Agilent Technologies, Santa Clara, California, USA) and quantified by Qubit HS assay.

Libraries were used for targeted sequence capture with the custom probe set MyBaits Compositae-1061 kit (Mandel et al., 2014) from Arbor Biosciences (Ann Arbor, Michigan, USA) following manufacturer’s protocols (version 2.3.1). About 500ng of total DNA library in 6µl of library, or equal amount of DNA per sample when pooling (maximum of 8 samples), was used in each MyBaits reaction for hybridization with the baits for 36 hours. Hybridized targets were recovered using Dynabeads M-280 Streptavidin (ThermoFisher Scientific, Vilnius, Lithuania) following the manufacturer’s protocol. Captured targets were amplified and quantified using KAPA library quantification kits (Kapa Biosystems, Wilmington, Massachusetts, USA). A 1:4 ratio of library was added to each of the captured target pools to obtain off-target plastid sequence data. The pooled libraries were sequenced on multiple instruments outlined in Supplemental Table 2.

### 2.3 Raw sequence processing

Raw FASTQ data were cleaned using the program Trimmomatic v. 0.36 (Bolger et al., 2014) which removes Illumina adapters from low-quality bases and adapter contamination. Trimmomatic offers two main quality trimming options: Sliding Window quality filter or Maximum Information quality filter. The Sliding Window quality filter scans from the 5’ end of the read and removes the 3’ end when the average quality within the window falls below a fixed threshold. The Maximum Information quality filter applies a trimming process that becomes more increasingly strict as it progresses through the read, rather than applying a fixed quality threshold. A recent study has shown that having less strict trimming methods is better for phylogenetic analyses (Portik & Wiens, 2020), so the Sliding Window quality filter was used in this analysis (illuminaclip 2:30:10, leading 20, trailing 20, sliding window 5:20). Cleaned reads were retained when they had a minimum length of 36 bp and corresponding forward and reverse pairs.

### 2.4 Nuclear ortholog assignment and assembly

The HybPiper v. 1.3.1 (Johnson et al., 2016) pipeline was used to match the reads to the target loci contained in the probe set (Mandel et al., 2014). HybPiper was chosen since it does not remove loci with paralogous sequences (hereafter referred to as ‘paralogs’), but instead chooses a single contig that has the greatest coverage depth or greatest percent identity to the reference sequence (Johnson et al., 2016). HybPiper is better suited to our data since higher ploidy levels, such as what is seen in *Packera*, are typically associated with greater numbers of paralogs. A combined reference/*de novo* assembly was performed using programs BWA v. 0.7.17 (Li & Durbin, 2009), which maps the pair-end reads to the targets, and SPAdes v. 3.5 (Bankevich et al., 2012), which assembles the pair-end reads into contigs with specified kmer lengths (21, 33, 55, 77, 99). A multifasta was generated for each locus containing the sequences of that specific locus for all taxa in the run. These multifastas were aligned into “supercontigs”, which contains both target and off-target sequence data, separately using MAFFT v. 7.407 (Katoh & Standley, 2013), and concatenated using FASconCAT-G v. 1.02 (Kück & Longo, 2014). Paralog detection was carried out on all exons using the “paralog_investigator” option in HybPiper (https://github.com/mossmatters/HybPiper/wiki/Paralogs) which flags loci that designate potential paralogs.

### 2.5 Plastid assembly

Plastid DNA was not targeted during sequence capture. Instead, reference mapping was performed to partially assemble plastid genes from off-target reads to a plastid genome using Bowtie2 v. 2.3.5.1 (Langmead & Salzberg, 2012; Langmead et al., 2019). Bowtie2 mapped reads of each sample against the published *Helianthus annuus* L. plastid genome (GenBank accession NC_007977; Timme et al., 2007). Then a consensus sequence for each assembly was obtained using SAMtools v. 1.9 and BCFtools v. 1.9 (Li et al., 2009). To obtain genes for a coalescent based species tree estimation analysis (hereafter referred to as “pseudo-coalescent”), consensus sequences were annotated using the online platform GeSeq (https://chlorobox.mpimp-golm.mpg.de/geseq.html; Tillich et al., 2017). The General Feature Format (gff3) output files were uploaded to Geneious v. 11.1.5 (https://www.geneious.com), along with a corresponding aligned consensus sequence. By importing both files, Geneious automatically annotates the genes to the aligned consensus sequence. Protein-coding gene sequences of individuals were then manually extracted for phylogenetic analyses and any genes that contained duplicates were removed.

### 2.6 Phylogenetic analyses

The final concatenated alignment of all gene trees from the HybPiper pipeline and plastid assembly were implemented in PartitionFinder v. 2.1.1 (Lanfear et al., 2012) to determine which nucleotide substitution model was most appropriate given the data provided. The overall best fitting model, GTR+I+Γ, which allows gamma-distributed substitution rates and a proportion of invariant sites, was used to build concatenated nuclear and plastid maximum likelihood trees in RAxML v. 8.1.3 (Stamatakis, 2014) with 1,000 bootstrap replicates. For the pseudo-coalescent analysis, nuclear and plastid genes were separated depending on which suitable nucleotide substitution model best fit each gene for the RAxML run (GTR, GTR+Γ, GTR+I+Γ). These gene matrices were implemented in ASTRAL-III v. 5.7.3 (Zhang et al., 2018), hereafter referred to as “ASTRAL”, to generate species trees with local posterior probability (LPP) values for branch support. LPP values indicate the probability that the branch is the true branch given the set of gene trees provided. LPP is considered the more reliable clade support measure when running ASTRAL since LPP is computed based on a quartet score (Sayyari & Mirarab, 2016) and assumes incomplete lineage sorting (ILS). All trees were visualized using the package phytools (Revell, 2012) in R v. 4.0.5 (R Core Team, 2016; RStudio, 2020).

### 2.7 Molecular clock dating

Estimates of divergence times were calculated on the best fitting nuclear tree using two relaxed-clock methods: treePL v. 1.0 (Smith & O’Meara, 2012) which utilizes the penalized likelihood (PL) method (Sanderson, 1997, 2002), and RelTime (Tamura et al., 2012, 2018) which utilizes the relative rate framework (RRF). Both methods are efficient for larger datasets, though recent studies have shown that RelTime produces more similar results to Bayesian approaches than treePL or other non-Bayesian dating methods (Tao et al., 2020; Barba-Montoya, 2021; Costa et al., 2022). Even so, both methods were tested to check for consistency in divergence time estimates.

The treePL analysis was implemented using command-line and followed the step-by-step protocol provided by Maurin (2020). Given that ASTRAL does not result in a topology with branch lengths in substitution rates, branch lengths were first re-estimated by fitting the concatenated matrix to the best-fitting ASTRAL species topology in RAxML using the -g command, which constrains to the best-fitting maximum likelihood (ML) tree topology, and -k command, which prints branch lengths on the bootstrap replicates. To estimate the 95% confidence intervals (CI) for divergence dates, we generated 100 bootstrap replications of our concatenated matrix and used the replicates to produce 100 trees with re-estimated branch lengths given the best-fitting ASTRAL topology.

The treePL analysis was primed five times to determine the best optimization parameters (lowest *opt* and *optad* values), then was time-calibrated with the thorough setting. Next, the replicate cross validation (CV) analysis was conducted using five different *cvstop* values (0.1, 0.001 10^-8^, 10^-20^, and 10^-30^) on the best ML tree to determine the best smoothing value, indicated by the smallest chi-square value. If multiple best smoothing values resulted from the five tests, then the overall lowest smoothing value was used. Both the best ML tree (noBS) and bootstrap replicates (BS) were dated, and results of the bootstrap replicates were summarized into a consensus tree to obtain a 95% CI using TreeAnnotator v. 2.5.2 (Bouckaert et al., 2019) with default settings.

RelTime was implemented using the command-line version of the MEGA software, MEGA-CC v. 11.0.11 (Kumar et al., 2012, 2016). MEGA-CC requires a special input file called a MEGA Analysis Options (.mao) file which specifies the analysis to run, as well as the analysis options to use. This .mao file was created by using the graphical user interface (GUI) software MEGA-X: Molecular Evolutionary Genetics Analysis across computing platforms (Kumar et al., 2018). Additionally, MEGA-CC requires a sequence alignment file in fasta or MEGA format, an aligned phylogenetic tree in Newick format, and a calibration and outgroup file indicating calibration points and identifying outgroups, respectively (Mello, 2018). Given that RelTime does not date specified outgroups, two additional taxa in tribe Gnaphalieae (*Disparago ericoides* Gaertn. and *Pseudognaphalium obtusifolium* (L.) Hilliard & B.L.Burtt) were included in the RelTime analyses but were later pruned to bring the total number of taxa back to 106. Voucher information for these two added species is included in Supplemental Table 1.

There are no fossils available within *Packera* or Senecioneae, so instead, multiple dating scenarios were tested utilizing both primary and secondary age constraints (Table 1). Briefly, a maximum (59.95MY) and minimum (31MY) age was tested for the Anthemideae/Senecioneae split given those dates are the highest and lowest values within the 95% CI from Mandel et al. (2019)’s diversification rate analysis of Asteraceae. Additionally, a minimum age of 13.8MY was tested at the *Packera* crown node from age estimates provided by Mandel et al. (2019). Next, a minimum age constraint of 31MY was tested within the outgroup-Anthemideae clade given the presence of an *Artemisia* L. fossil pollen (Hobbs & Baldwin, 2013; Wang, 2004), along with a maximum age of 59.95MY to accommodate calibration requirements in RelTime analyses. Finally, a maximum age constraint of 14.3MY was tested for *Pericallis* D.Don, a genus endemic to the Macaronesia islands (Jones et al., 2014), since that is the estimated age of Porto Santo, the oldest volcanic archipelago within the Madeira Islands (McDougall & Schmincke, 1976; Geldmacher et al., 2000).

**Table 1.** Dating scenarios with various age constraints (A-F) implemented in treePL and RelTime. **y** = age constraint was used in the scenario; **n** = age constraint was not used in the scenario.

|  | *Calibration type* | *Max or Min* | **Scenario 1** | **Scenario 2** | **Scenario 3** | **Scenario 4** | **Scenario 5** | **Scenario 6** | **Scenario 7** | **Scenario 8** |
| --- | --- | --- | --- | --- | --- | --- | --- | --- | --- | --- |
| **A:** Senecioneae/Anthemideae split - 59.95MY (max)<sup>†</sup> | *Secondary* | *Max* | y | y | y | y | n | n | n | n |
| **B:** Senecioneae/Anthemideae split - 31MY (min)<sup>†</sup> | *Secondary* | *Min* | y | y | n | n | y | y | n | n |
| **C:** *Artemisia* [Anthemideae] pollen fossil - 59.95MY (max)<sup>‡</sup> | *Secondary* | *Max* | y | y | y | y | y | y | y | y |
| **D:** *Artemisia* [Anthemideae] pollen fossil - 31MY (min)<sup>‡</sup> | *Primary* | *Min* | y | y | y | y | y | y | y | y |
| **E:** *Pericallis* - 14.3MY<sup>§</sup> | *Primary* | *Max* | y | y | y | y | y | y | y | y |
| **F:** *Packera* - 13.8MY<sup>†</sup> | *Secondary* | *Min* | y | n | y | n | y | n | y | n |

<sup>†</sup> Mandel JR, Dikow RB, Siniscalchi CM, Thapa R, Watson LE, Funk VA. 2019. A fully resolved backbone phylogeny reveals numerous dispersals and explosive diversifications throughout the history of Asteraceae. *Proceedings of the National Academy of Sciences of the United States of America* 116: 14083–14088.
<sup>‡</sup> Hobbs CR, Baldwin BG. 2013. Asian origin and upslope migration of Hawaiian *Artemisia* (Compositae-Anthemideae). *Journal of Biogeography* 40: 442–454. Wang WM. 2004. On the origin and development of *Artemisia* (Asteraceae) in the geological past. *The Botanical Journal of the Linnean Society* 145: 331–336.
<sup>§</sup> Geldmacher J, Van Den Bogaard P, Hoernle K, Schmincke HU. 2000. The <sup>40</sup>Ar/<sup>39</sup>Ar age dating of the Madeira Archipelago and hotspot track (eastern North Atlantic). *Geochemistry, Geophysics, Geosystems* 1.
McDougall I, Schmincke HU. 1976. Geochronology of Gran Canaria, Canary Islands: Age of shield building volcanism and other magmatic phases. *Bulletin Volcanologique* 40: 57–77.

All output files are provided in Supplemental Materials. Results of all tested scenarios were compared for biological relevance, though only one scenario of each dating method, treePL and RelTime, will be discussed.

### 2.8 Biogeography

Ancestral ranges were estimated on the dated phylogeny generated with treePL using the package BioGeoBEARS v. 1.1 (Matzke, 2013) in R. BioGeoBEARS estimates historical, biogeographic ranges by implementing popular models in a likelihood framework. Some of the most common models used are dispersal-extinction-cladogenesis (DEC; Ree & Smith, 2008), the likelihood version of DIVA (DIVALIKE; Ronquist, 1997), and the BayArea likelihood version of the range evolution model (BAYAREALIKE; Landis et al., 2013). In BioGeoBEARS, founder event speciation can be added to any of these models (*+J*) with flexibility in determining relative probability. Likelihood Ratio Test (LRT) and Akaike Information Criterion (AIC) was used to determine which model best fit the data.

All *Packera* taxa, along with their four most closely related outgroup genera (*Pericallis*, *Elekmania* B.Nord., *Werneria* Kunth, and *Xenophyllum* V.A.Funk), were coded as present or absent in ten geographic ranges outlined in Takhtajan (1986; Supplemental Table 4): Arctic Province, Canadian Province, Appalachian Province, Atlantic and Gulf Coast Plain Province, North American Prairies Province, Rocky Mountain and Vancouvarian Province, Madrean Region, Asia, Europe, and Central America. Distributions were determined from previously published literature (Freeman & Barkley, 1995; Trock, 1999, 2006; Gramling, 2006; Mahoney & Kowal, 2008; Elven et al., 2011; Yeatts, et al., 2011). Only four of the 23 outgroup taxa were chosen to limit the number of geographic ranges used in the analysis. The remaining outgroup taxa were pruned from the dated tree generated with treePL using the *pxrmt* function in phyx (Brown et al., 2017). Each taxon was limited to a max distribution of four ranges.

## 3 Results

### 2.1 Matrix results

Illumina sequencing resulted in approximately 723.5 million reads across 106 taxa. The minimum and maximum number of reads ranged from 535,934 in *Werneria aretioides* Wedd. and 20,685,576 in *Senecio vulgaris* L. (full list in Supplemental Table 2). The nuclear supermatrix comprised 531,668 characters spanning all 106 taxa and included 1,052 of the targeted 1,061 loci. The number of loci recovered for each taxon ranged from 613 in *Tephroseris newcombei* (Greene) B.Nord. & Pelser to 1,051 in *Roldana gilgii* (Greenm.) H.Rob. & Brettell. The supermatrix contained 19.6% missing data and 130,541 sites (24.55%) were parsimony informative. Out of the 1,052 loci included, 813 loci had paralog warnings (Supplemental Table 3). Three loci were removed when running ASTRAL since three or fewer taxa were present in those loci, totaling 1,049 loci in the pseudo-coalescent tree. ASTRAL reported that the nuclear tree had a normalized quartet score of 0.493, indicating that 49% of the gene trees match the final species tree. Off-target plastid sequencing resulted in a supermatrix composed of 80,160 characters spanning 106 taxa and includes 79 of the 86 recovered, protein-coding loci. This supermatrix contained 4.6% missing data and 3,099 sites (3.87%) were parsimony informative. Seven of the original annotated genes located in the repeat region were removed from the analysis (Supplemental Table 3).

Raw read and assembly information for the 106 taxa can be found in Supplemental Tables 2 & 3. General matrix information and summary statistics can be found in Supplemental Table 3. Raw data are deposited in NCBI (BioProject: PRJNA907383). Given the matrix statistics and biological relevance, the nuclear tree constructed and inferred using RAxML with a pseudo coalescent approach in ASTRAL, and the plastid tree constructed with a concatenation approach, were determined to be the best fitting species trees, and will be presented below. The remaining trees, the nuclear tree inferred using a concatenation approach and the ASTRAL plastid tree, are presented in Supplemental Figs. 1 & 2, respectively, but will not be discussed further.

**Fig. 2.**
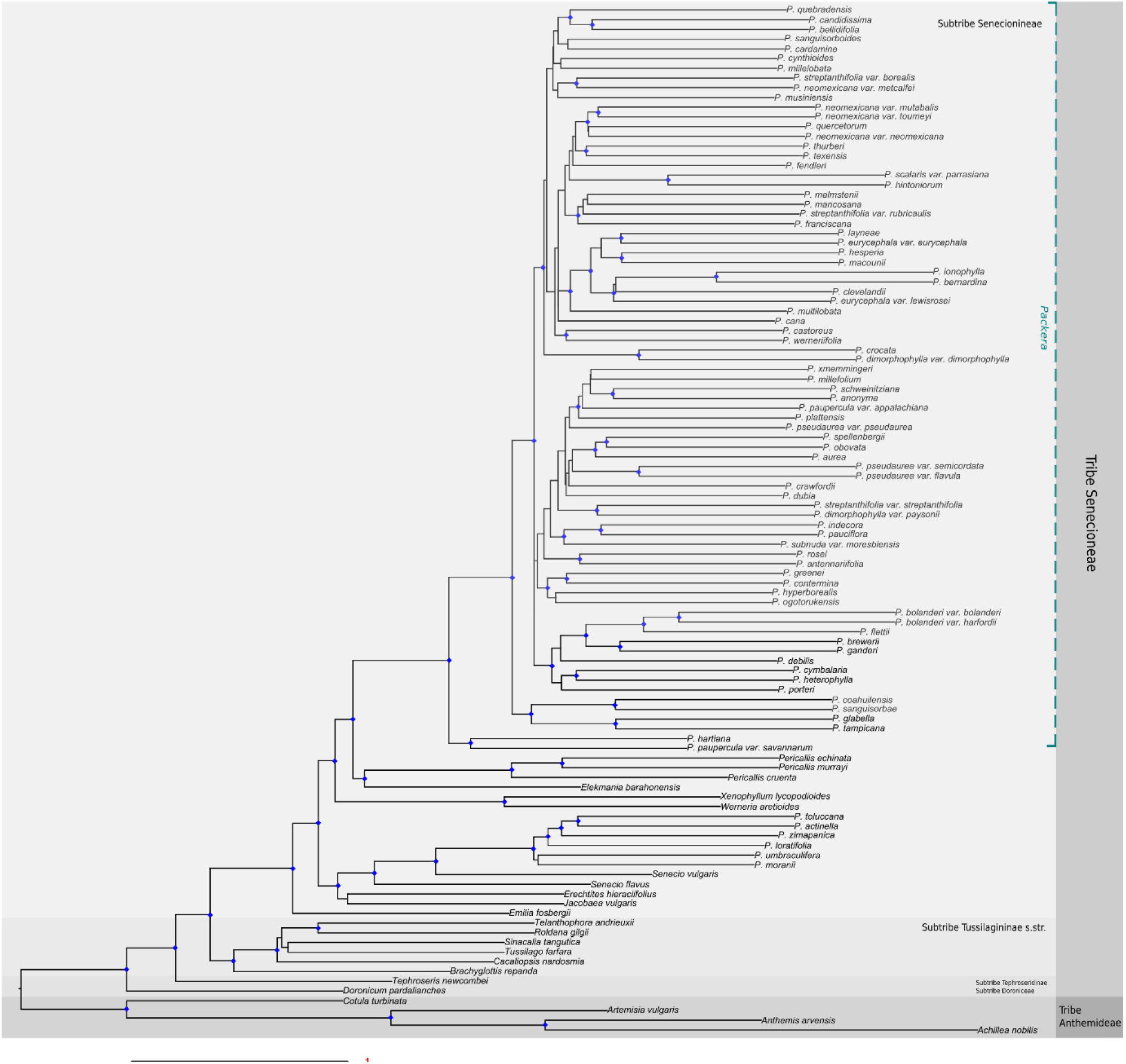
Nuclear phylogeny produced using a pseudocoalescent analysis (ASTRAL-III) and inferred with a maximum likelihood approach (RAxML). Scale bar to the bottom-left of the tree represents the mean number of nucleotide substitutions per site. Support values (LPP) of 0.9 or greater are marked with a blue diamond at the node. Taxa are highlighted in varying levels of gray according to tribal/subtribal status.

### 3.2 Nuclear vs. plastid trees

Phylogenetic topologies were highly incongruent with most *Packera* species being placed in different clades depending on sequence type (nuclear or plastid; Figs. 2 & 3, respectively) or phylogenetic method (i.e., concatenation instead or pseudo-coalescence; Supplemental Figs. 1 & 2, respectively). The nuclear tree showed moderate resolution with 51% of the nodes having <0.95LPP in the nuclear tree (Supplemental Fig. 3). In contrast, the resulting plastid tree showed much lower resolution with fewer nodes having higher support (89% of nodes have <0.95BS; Supplemental Fig. 4). Most of the topological conflict was present among *Packera* taxa, with minor differences among the remaining Senecioneae (Supplemental Fig. 5). Both the nuclear and plastid phylogenies show separation between subtribes Senecionineae and Tussilaginineae s.str. (Supplemental Fig. 5), though the species relationships within the subtribes varied.

**Fig. 3.**
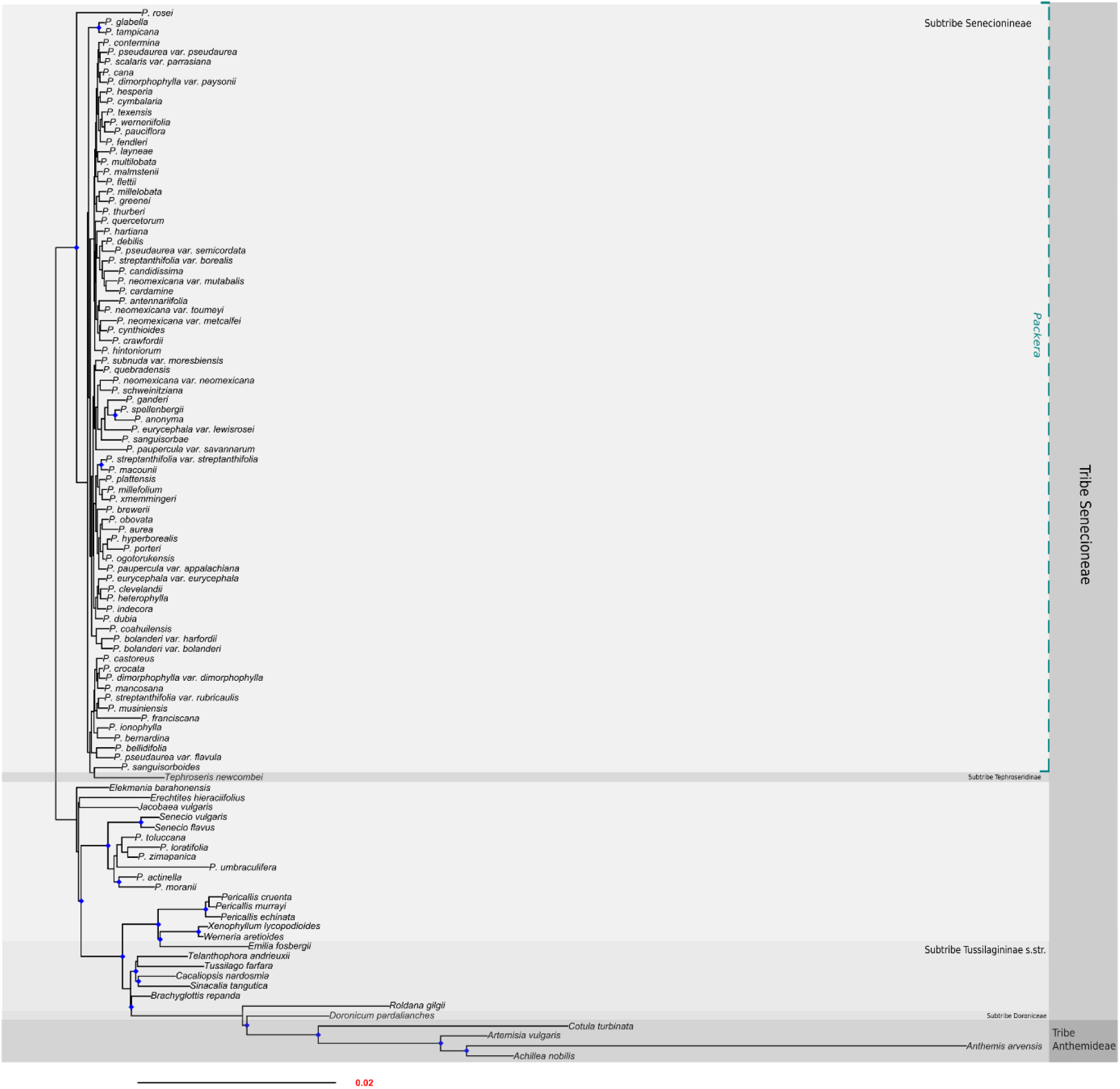
Plastome phylogeny produced using a concatenated approach, inferred with a maximum likelihood approach (RAxML). Scale bar to the bottom-left of the tree represents the mean number of nucleotide substitutions per site. Support values (bootstrap) of 90 or greater are marked with a blue diamond at the node. Taxa are highlighted in varying levels of gray according to tribal/subtribal status.

### 3.3 Molecular clock dating

Age estimates differed between dating methods. Using treePL, the resulting divergence rate estimates on all eight tested scenarios provided a range of ages for crown *Packera*, from 25.1MY in Scenario 1 to 31.1MY in Scenario 6. The Senecioneae/Anthemideae split ranged from 59MY in Scenario 1 to 78.1MY in Scenario 6. With RelTime, age estimates were generally smaller and ranged from 15.5MY in Scenarios 1 and 5 and 19.2MY for crown *Packera* in Scenarios 4 and 8, and 49.9MY in Scenarios 2 and 6 and 55.3MY for the Senecioneae/Anthemideae split in Scenarios 4 and 8.

Scenario 4 was selected as the overall best fitting dating scenario since it included all tested primary calibration points, and included the least amount of secondary calibration points, which provided a max age for the Senecioneae/Anthemideae split (Table 1). Although secondary calibration points are not recommended unless primary calibrations are not available (e.g., Müller & Reisz, 2005; Benton & Donoghue, 2007; Parham et al., 2012), the inclusion of these secondary calibrations were necessary to limit the age for the group. Without this maximum value, the ages drastically increased and were biologically unreasonable given what is known about the timing of divergences in the family (Scenarios 5-8, Supplemental Figs. 10-13 & 18-21). Scenario 4 estimated crown Senecioneae to be 50.4MY and crown *Packera* at 25.9MY with treePL (Fig. 4), and crown Senecioneae to be 55.3MY and crown *Packera* at 19.2MY with RelTime (Fig. 5). The dated trees from all eight scenarios can be found in Supplemental Materials.

**Fig. 4.**
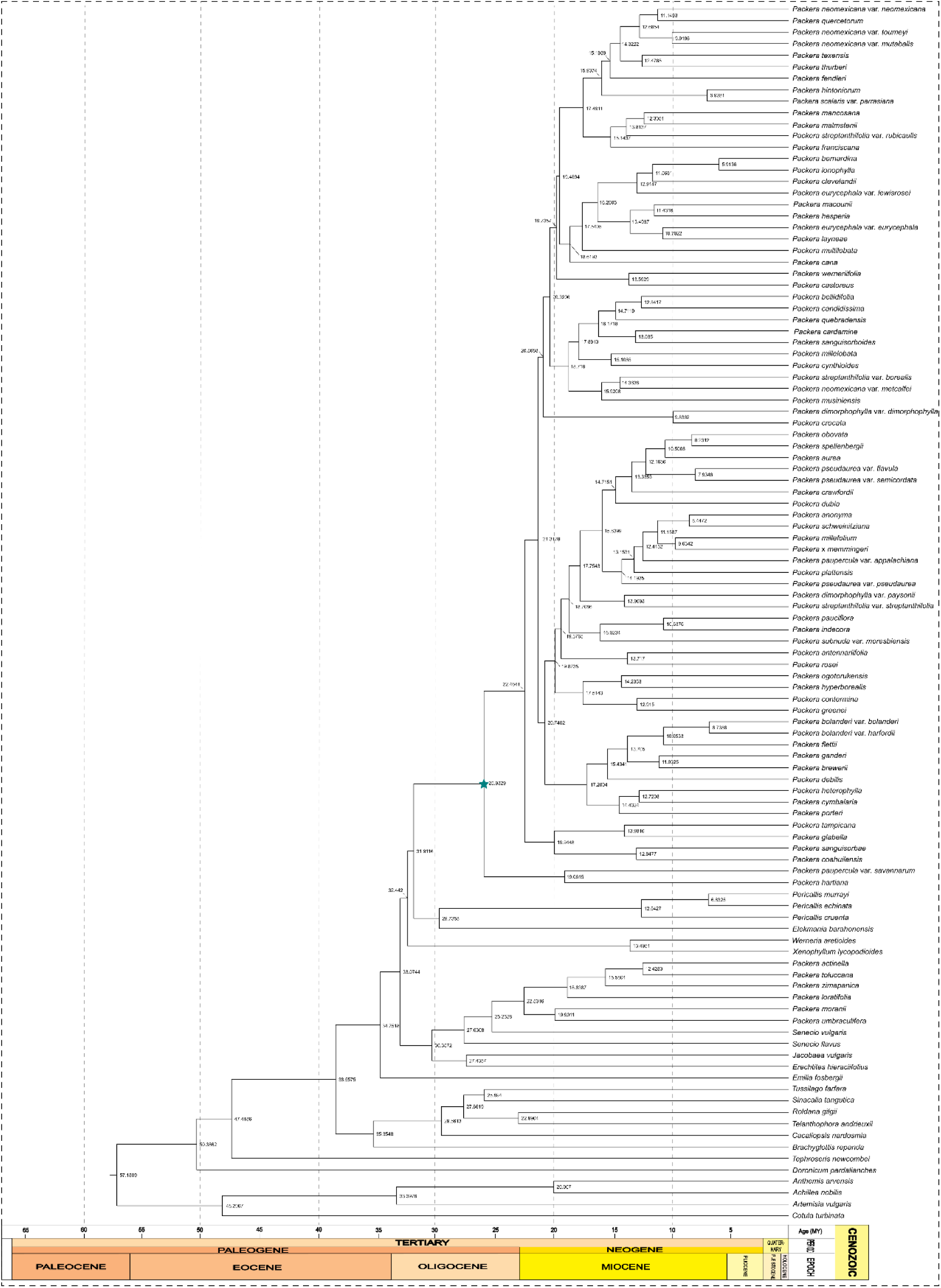
Time-calibrated nuclear phylogeny of *Packera* produced using treePL. Numbers next to each node correspond to the mean estimated age of that node. Crown *Packera* is indicated by a teal star.

**Fig. 5.**
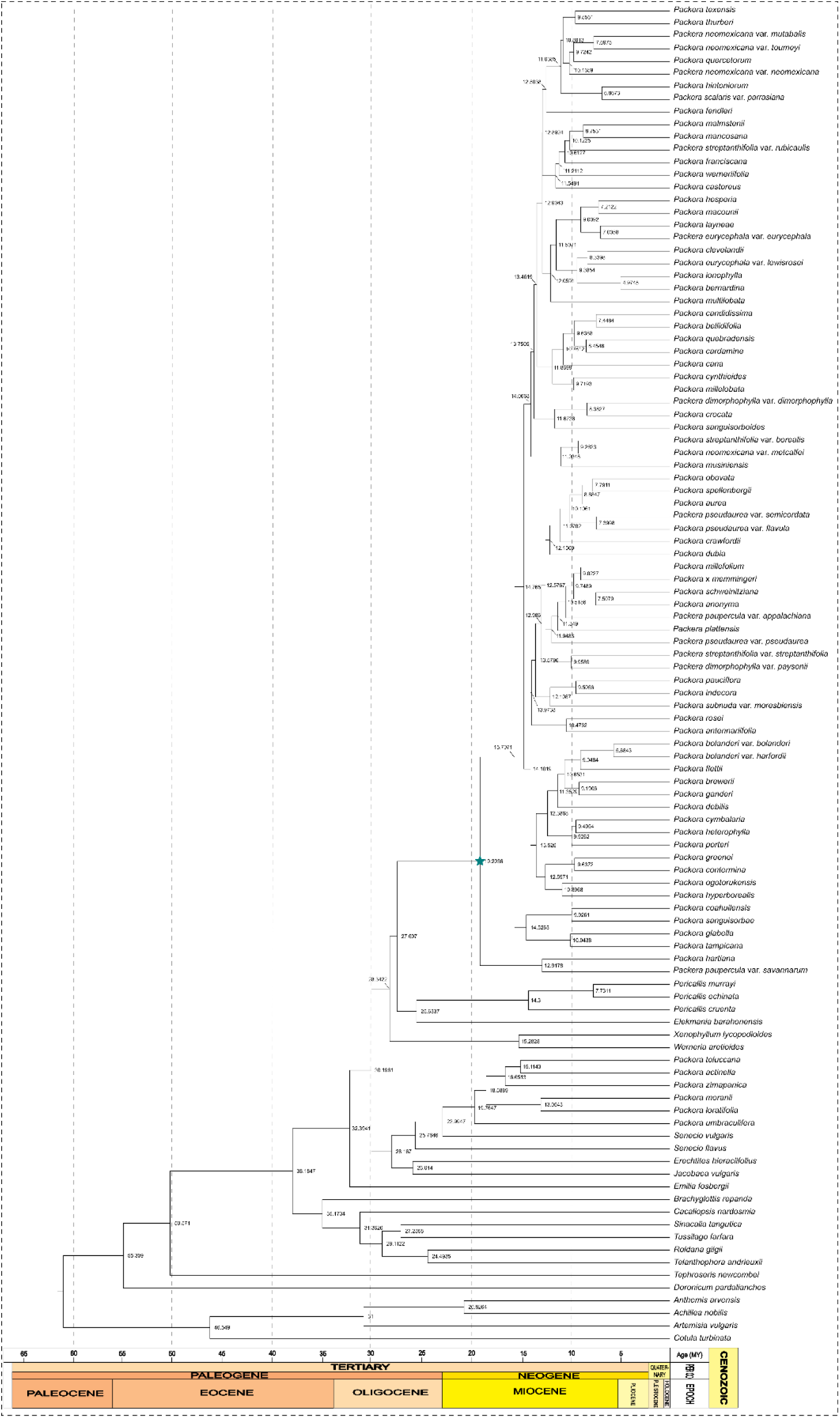
Time-calibrated nuclear phylogeny of *Packera* produced using RelTime. Numbers next to each node correspond to the mean estimated age of that node. Crown *Packera* is indicated by a teal star.

The dates within all tested scenarios were similar to what was expected given previous work in Asteraceae or Senecioneae, with RelTime dates being younger than treePL and aligning more with ages previously predicted in family-wide phylogenies (Pelser et al., 2010; Huang et al., 2016; Mandel et al., 2019; Zhang et al., 2021). Recent family-level phylogenies estimated the age of Senecioneae to be 36MY or 55MY, and *Packera* to be around 13.8MY or 20.2MY (Mandel et al., 2019; Zhang et al., 2021, respectively). Our results lie within the 95% estimated range from both analyses. Interestingly, the most recent tribe-level nuclear phylogeny yielded much younger results, with Senecioneae estimated to be 35MY and *Packera* to be 7.8MY (Pelser et al., 2010). This may be explained by the use of different calibration points, samples used, or dating methods.

### 3.4 Biogeography

Evaluation of the six different model choices showed that BAYAREALIKE was the best fit given our data considering that it has the lowest log likelihood (Ln *L*) and AIC scores, and second highest weighted AICc score (Table 2). The BAYAREALIKE model is a simplified likelihood interpretation of the BayArea program described by Landis et al. (2013), which tests the importance of cladogenesis on a particular dataset, but it does not depend on distance between taxa and allows for a larger number of areas. Generally, models including jump speciation (+*J*) provided a better fit than those without in the DEC and DIVALIKE models, and were the same or very similar in the BAYAREALIKE models (with and without *J*: Ln *L* = -259.09; Table 2), indicating a potential for jump dispersal or founder-event speciation in our data. Results of all model runs can be found in Supplemental Figs. 22-33.

**Table 2.** Biogeographical model selection for all *Packera* taxa and four outgroup genera (*Pericallis*, *Elekmania*, *Werneria*, *Xenophyllum*). Ln *L* = log likelihood values; Num params = number of free parameters; *d* = “dispersal” parameter; *e* = “extinction” parameter; *j* = “jump” parameter which specifies the weight of each jump dispersal event in the cladogenesis matrix; AIC = Akaike information criterion; AICc = Akaike information criterion with corrections for small sample sizes. “wt” represents the resulting model weights with AIC and AICc.

|  | Ln L | Num<br>params | d | e | j | AIC | AICc | AIC wt | AICc wt |
| --- | --- | --- | --- | --- | --- | --- | --- | --- | --- |
| DEC | -278.0081 | 2 | 0.00419 | 1.00e-12 | 0 | 560.0163 | 560.1591 | 4.43530e-09 | 4.52087e-09 |
| DEC+J | -273.1804 | 3 | 0.00388 | 1.00e-12 | 0.0078 | 552.3608 | 552.6500 | 2.03839e-07 | 1.93115e-07 |
| DIVALIKE | -291.6979 | 2 | 0.0051 | 1.00e-12 | 0 | 587.3957 | 587.5386 | 5.02983e-15 | 5.12687e-15 |
| DIVALIKE+J | -281.5982 | 3 | 0.00422 | 1.00e-12 | 0.0116 | 569.1963 | 569.4855 | 4.50301e-11 | 4.26612e-11 |
| BAYAREALIKE | -259.0869 | 2 | 0.00136 | 3.77e-02 | 0 | 522.1738 | 522.3166 | 7.31667e-01 | 7.45783e-01 |
| BAYAREALIKE+J | -259.0900 | 3 | 0.00136 | 3.77e-02 | 0.00001 | 524.1800 | 524.4691 | 2.68332e-01 | 2.54216e-01 |

## 4 Discussion

### 4.1 Nuclear vs. plastid trees

The results of our analyses showed that nuclear and plastid phylogenies are incongruent with each tree showing different topologies. Although no study has directly compared nuclear and plastid sequences among *Packera* species, earlier phylogenetic work in tribe Senecioneae recovered plastid vs. nuclear incongruences between the *Packera* and other Nearctic and Neotropical taxa (Pelser et al., 2010). Additionally, this work showed that *Packera*’s phylogenetic placement changed depending on which genera were included, indicating the potential for long branch attraction (Felsenstein, 1978), which was tested and ruled out by Pelser et al. (2010).

The most recent phylogenetic work on *Packera* demonstrated an immense lack of variation in ITS regions, with a range of only 0-5% sequence divergence and some species having identical sequences (0% sequence divergence; Bain & Golden, 2000). The resulting ITS tree had many unresolved areas, likely due to few phylogenetically informative sites, low species sampling, or reticulation events between species. Using target-enrichment data instead of ITS increased phylogenetic resolution at lower taxonomic levels; however, branch support remained low at many of the nodes with 51% of the nodes having <0.95LPP in the nuclear tree.

The differing phylogenetic topologies seen in our study can potentially be explained by the low levels of genetic variation found within *Packera*’s plastid sequences (Bain & Jansen, 1996). Plastid sequences within *Packera* have been shown to contain a high level of intra-populational DNA polymorphisms and a low level of interspecific genetic variation (Bain & Jansen, 1996; Golden & Bain, 2000; Panero & Funk, 2008; Funk et al., 2009). Additionally, plastid diversity in members of Compositae is generally low, making it difficult to distinguish evolutionary relationships within the family (Jansen et al., 1992; Bain & Jansen, 2006). Given this low resolution in the plastid tree, it is unclear whether the nuclear-plastid incongruence is due to underlying discordance or simply low phylogenetic signal in the plastid genome. There does not appear to be any morphological, ecological, or biogeographic trends supporting the topology of the clades within the plastid tree. For example, previous plastid haplotype work has shown that *P. sanguisorboides* (Rydb.) W.A.Weber & Á.Löve and *P. bernardina* (Greene) W.A.Weber & Á.Löve share a number of plastid DNA similarities (Bain & Jansen, 1996), though our plastid tree shows they are phylogenetically distant (Fig. 3). Additionally, the most recent *Packera* phylogeny has *P. bernardina* and *P. sanguisorboides* within the same smaller clade, though unresolved (Bain & Golden, 2000).

### 4.2 *Packera* relationship trends

The results of our study show that *Packera* as currently circumscribed is not monophyletic. Some *Packera* species, *P. actinella* (Greene) W.A.Weber & Á.Löve, *P. loratifolia* (Greenm.)W.A.Weber & Á.Löve, *P. toluccana* (DC.) W.A.Weber & Á.Löve, *P. umbraculifera* (S.Watson) W.A.Weber & Á.Löve, *P. zimapanica* (Hemsl.) C.C.Freeman & T.M.Barkley, and *P. moranii* (T.M.Barkley) C.Jeffrey do not fall within *Packera* but instead with the outgroups in a *Senecio* clade (Figs. 2 & 3). Most of these species, excluding *P. moranii*, have been previously proposed as potentially misclassified *Packera* species. For example, molecular phylogenetic work has shown that *P*. *actinella*, *P. loratifolia*, and *P*. *zimapanica* never grouped within *Packera* (Bain & Jansen, 1995; Bain & Golden, 2000; Pelser et al., 2007), but instead within *Senecio* in a group called the Lugentes (Barkley, 1985). The Lugentes have a base chromosome count of n = 20 (Ornduff et al., 1967), which differs from *Packera*’s base chromosome counts of n = 22, 23 (Trock, 2006). *Packera toluccana* is reported to have n = 20, and both *P. toluccana* and *P. zimapanica* have different pollen types (Senecioid) than what is characteristic of *Packera* (Helianthoid; Ornduff et al., 1967; Bain et al., 1997). In addition, studies in *P. umbraculifera* showed that it was closely related to *P. loratifolia* and may require placement back into the genus *Senecio* (Greenman, 1907). Therefore, our phylogeny provides further support for those species removal from *Packera* and back into *Senecio*.

Surprisingly, *P. moranii* did not place within *Packera* but instead within the outgroup *Senecio* clade. This was not expected since *P. moranii* has the same chromosome counts as most *Packera* taxa (n = 23; Barkley, 1978), and its taxonomic placement has never been questioned. A possible explanation for *P. moranii*’s placement is that the herbarium specimen used for this phylogeny was misidentified. However, we do not believe this is the case considering the specimen used is a paratype of *P. moranii* and was collected in the Sierra de San Pedro Mártir region of Baja California, Mexico, where it is the only known *Packera* species of that region (Freeman & Barkley, 1995; Rebman et al., 2016). There is a similar plant in Baja California that most likely represents an undescribed taxon; however, those plants are found in the Sierra de La Libertad and northern Baja California Sur regions, instead of the Sierra de San Pedro Mártir regions where *P. moranii* is located (Rebman et al., 2016). Therefore, the taxonomic assignment of *P. moranii* to *Packera* should be investigated further.

Most of the varieties included in this study are not grouping with their infraspecifcs (i.e., *Packera dimorphophylla* (Greene) W.A.Weber & Á.Löve varieties, *P. pseudaurea* (Rydb.) W.A.Weber & Á.Löve varieties, *P. streptanthifolia* (Greene) W.A.Weber & Á.Löve varieties, *P. eurycephala* (Torr. & A.Gray) W.A.Weber & Á.Löve varieties, *P. paupercula* (Michx.) Á.Löve & D.Löve varieties, etc.; Fig. 2). This was not expected since varieties are still typically seen as most closely associated to their infraspecifics, but usually differ from minute differences in morphology, ecology, habitat, etc. Therefore, it is still assumed the varieties would be sister to or within the same smaller clade with the infraspecific species. A potential explanation for this separation is that these varieties are actually distinct species, separate from their infraspecifics, and should be recognized as new species instead of varieties. Alternatively, these findings could be the result of historical hybridization or introgression between distantly related species. For example, it has previously been predicted that *P. streptanthifolia* var. *rubicaulis* (Greene) Dorn arose from a hybridization event between *P. streptanthifolia* var. *streptanthifolia* and *P. multilobata* (Torr. & A.Gray) W.A.Weber & Á.Löve (Bain, 1988; Barkley, 1988; Bain & Jansen, 1995). Given the results of our phylogeny, *P. multilobata*, *P. streptanthifolia* var. *streptanthifolia*, and *P. streptanthifolia* var. *rubicaulis* all appear distantly related (Fig. 2). If *P. multilobata* and *P. streptanthifolia* var. *streptanthifolia* did hybridize, you would expect separation between taxa, which is what is seen in our phylogeny since *P. streptanthifolia* var. *streptanthifolia* are separated from both varieties, *P. streptanthifolia* var. *borealis* (Torr. & A.Gray) Trock and *P. streptanthifolia* var. *rubicaulis*, and *P. multilobata*. Both predictions would require additional taxonomic work that goes beyond the scope of this study.

Many widespread species (i.e., *Packera cana* (Hook.) W.A.Weber & Á.Löve, *P. multilobata*, *P. paupercula*, *P. pseudaurea*, etc.) are recovered as unexpected relationships in the phylogeny. For example, Trock (1999) had predicted that *P. multilobata* would have affiliations with multiple taxa, such as *P. breweri* (Burtt Davy) W.A.Weber & Á.Löve, *P. quercetorum* (Greene) C.Jeffrey, *P. streptanthifolia*, *P. millelobata* (Rydb.) W.A.Weber & Á.Löve, and *P. neomexicana* var*. mutabilis* (Greene) W.A.Weber & Á.Löve. However, *P. multilobata* is not closely related to any of those taxa in our nuclear or plastid trees (Figs. 2 & 3, respectively). Interestingly, *P. multilobata* is showing a closer relationship to another widespread species, *P. cana*, which is morphologically and ecologically distinct from *P. multilobata*. Given the amount of introgression and hybridization present in the genus (see Supplemental Material), we hypothesize that these widespread species have a greater chance of encountering other taxa in different geographic regions, resulting in more genetic admixture, and ultimately influencing the phylogenetic relationships seen in our tree. Future studies including hybridization analyses would be interesting to test this hypothesis.

The results of our nuclear phylogeny indicate that geography appears to have a large influence on species relationships within *Packera*, supporting previous predictions made by other botanists (Bain & Jansen, 1995; Barkley, 1988; Trock, 1999; Bain & Golden, 2000). For example, Bain & Jansen (1995) showed that although *P. sanguisorboides*, *P. neomexicana* var. *toumeyi* (Greene) Trock & T.M.Barkley, *P. bellidifolia* (Kunth) W.A.Weber & Á.Löve, and *P. multilobata* are not morphologically similar, all four taxa are from the Great Basin and northern Mexico region and placed within the same smaller clade, perhaps indicating shared common ancestry. Although geography has shown to be important for species relationships within this group, it is important to note that geography does not explain all our findings, which can be seen with *P. hartiana* (A.Heller) W.A.Weber & Á.Löve and *P. paupercula* var. *savannarum* R.R.Kowal, *P. spellenbergii* (T.M.Barkley) C.Jeffrey and *P. obovata* (Willd.) W.A.Weber & Á.Löve, and *P. rosei* (Greenm.) W.A.Weber & Á.Löve and *P. antennariifolia* (Britton) W.A.Weber & Á.Löve within our phylogeny. Our results demonstrate a trend in clade composition that appears influenced by species distribution.

Interestingly, *Tephroseris newcombei* (Greene) B.Nord. & Pelser placed outside of the Tussilagininae s.str. subtribe clade and sister to tribe Senecioneae in the nuclear tree but placed as a member of *Packera* in the plastid tree (Supplemental Fig. 5). Although *T. newcombei* was once placed in *Packera* as *P. newcombei* (Greene) W.A.Weber & Á.Löve (Weber & Löve, 1981), there is an overwhelming consensus that it does not belong in *Packera*, or possibly even within Senecioneae (Janovec & Barkley, 1996; Bain & Golden, 2000; Golden et al., 2001; Nordenstam & Pelser, 2011). A similar trend has been found in other groups of Asteraceae where phylogenies place species as ingroups or outgroups dependent on sequence type. For example, when investigating the evolutionary relationships between *Helianthus* L. and related genera, Schilling (2001) found that the plastid tree placed *Phoebanthus* as an ingroup to *Helianthus*, while the nuclear (ITS) tree showed *Phoebanthus* as sister to *Helianthus*. More recent phylogenetic research using a target enrichment approach found that *Phoebanthus* is most closely related to and sister to *Helianthus* (Stephens et al., 2015), supporting the nuclear relationship.

### 4.3 Molecular clock dating

Overall, our dates are similar to and lie within the upper divergence rate estimates from previous studies (Pelser et al., 2010; Huang et al., 2016; Mandel et al., 2019; Zhang et al., 2021). It was expected that *Packera* would lie within the lower age estimates since younger taxa tend to have a higher chance of hybridization and introgression given a higher genetic similarity, allowing hybridization to result into valid new species (Rieseberg & Willis, 2007). However, our estimated ages are the oldest found in any analyses, not supporting that prediction. Alternatively, some research has shown that the combination of hybridization and polyploidy may cause low genetic diversity between species (i.e., Alix et al., 2017; Śarhanov et al., 2017; Uhrinová et al., 2017), potentially explaining our results.

Divergence time estimates between RelTime and treePL differed (i.e., ∼6MY difference for crown *Packera*), with treePL producing overall older estimates than RelTime. This is not surprising given that previous studies testing the efficacy of Bayesian vs. non-Bayesian dating methods have shown that treePL typically overestimates the ages and has narrower confidence intervals than RelTime (Costa et al., 2022). Even so, treePL is still being used with larger datasets (i.e., Li et al., 2019; Janssens et al., 2020; Maurin et al., 2022) given it is computationally less intense than RelTime and other Bayesian dating methods (Mello et al., 2017; Tao et al., 2020; Barba-Montoya, 2021; Costa et al., 2022).

Though our estimated dates of *Packera* members from treePL are older than RelTime, they align well with earlier geohistorical hypotheses for the group, particularly by T.M. Barkley. Barkley (1988) proposed that *Packera* members originated sometime during the Arcto-Tertiary Geoflora in the later stages of the mid-Tertiary (∼22-37MY), providing support for our date of speciation than what was previously predicted. Barkley further anticipated that *Packera* species diversified more during the late Tertiary (∼15-34MY) when new and drier habitats formed in the Rocky Mountains, and the Sierra Nevada and Cascade uplifts developed. Additionally, he posited that new species developed in the cool and dry inner plateaus of the upper Cordilleran region and the drier and water climates in the southern inner Cordilleran regions (Great Basin and southward). Some of Barkley’s predictions are not reinforced by either of our dating approaches, such as the formation of mountaintop endemic species occurring during the Pleistocene (12,000-2.6MY). For example, Barkley predicted that *Packera schweinitziana* (Nutt.) W.A.Weber & Á.Löve, a mountaintop rock outcrop species, migrated south and became restricted to the Appalachian Mountains during the coldest parts of the ice age. However, our results suggest *P. schweinitziana* originated 8.4MY (min: 8.1MY, max: 8.7MY; treePL) or 7.5MY (min: 3.8MY, max: 14.9MY; RelTime) (Supplemental Materials), almost three to four times as old as Barkley’s prediction. Even so, our estimated dates are consistent with research into other high-elevation rock outcrop species, similar to *P. schweinitziana*, that predict dates later in the Miocene (Meseguer et al., 2015; Quinlan et al., 2020; Stubbs et al., 2020; Schilling et al., 2022).

It is important to note that Barkley made these predictions while considering *Packera*’s relationships on his named subgroups (Aurei, Lobati, and Tomentosi; Barkley, 1988), which are not supported in this phylogeny or others (Bain & Jansen, 1995; Bain & Golden, 2000). Even so, Barkley’s underlying explanations give insight into *Packera* and how climate/environment may have influenced diversification in the group. For example, Barkley originally predicted that with a changing climate, species were able to migrate continuously and frequently encounter other species, ultimately developing various combinations and creating opportunities for hybridization (Barkley, 1988), which follows with our current understanding of the group.

### 4.4 Biogeography and species relationships in core *Packera*

Core *Packera* likely originated in Central America or somewhere in the western temperate regions of North America, particularly within the North American Prairies Province, Rocky Mountain and Vancouvarian Province, or Madrean Region around 26MY during the Oligocene (Figs. 4 & 6). From 20MY to present day, some *Packera* species did not appear to expand, but instead reduced their ranges within western North America, which can be seen by a loss of regions/provinces through time. Further, it appears that most *Packera* species left the North American Prairies Province and settled west in the Rocky Mountain and Vancouvarian Province, or south into the Madrean region. These trends are consistent with late-Tertiary events where the continuing uplifts of the Rocky Mountain system, plus the development of the Sierra Nevada and Cascade uplifts during the mid-Tertiary, gave rise to extensive new and drier habitats along the mountain ranges of western United States (Stanley, 1986), likely causing species to have restricted or narrower ranges. Around the same time, there appears to be a large expansion of *Packera* species into the Atlantic and Gulf Coast Plain and Appalachian Provinces during the early Miocene, around 21-20MY. This trend may relate to falling sea levels in the Atlantic Ocean, ultimately allowing plant species, particularly herbaceous plants, to expand their ranges more eastward than before (Stanley, 1986). Later, *Packera* members diversified multiple times north into the Arctic and Canadian Provinces or Asia during the mid-Miocene, around 16MY, aligning with the great Arcto-Tertiary Geoflora, when the north of North America had cooler, drier climates and was covered by a diverse mix of hardwood or hardwood-coniferous forests during the mid to late Miocene, allowing plants to extend their ranges northward. With a continuously changing climate, it is predicted that species migrated perpetually and frequently encountered other species providing opportunities for hybridization and introgression, developing the various combinations seen today.

**Fig. 6.**
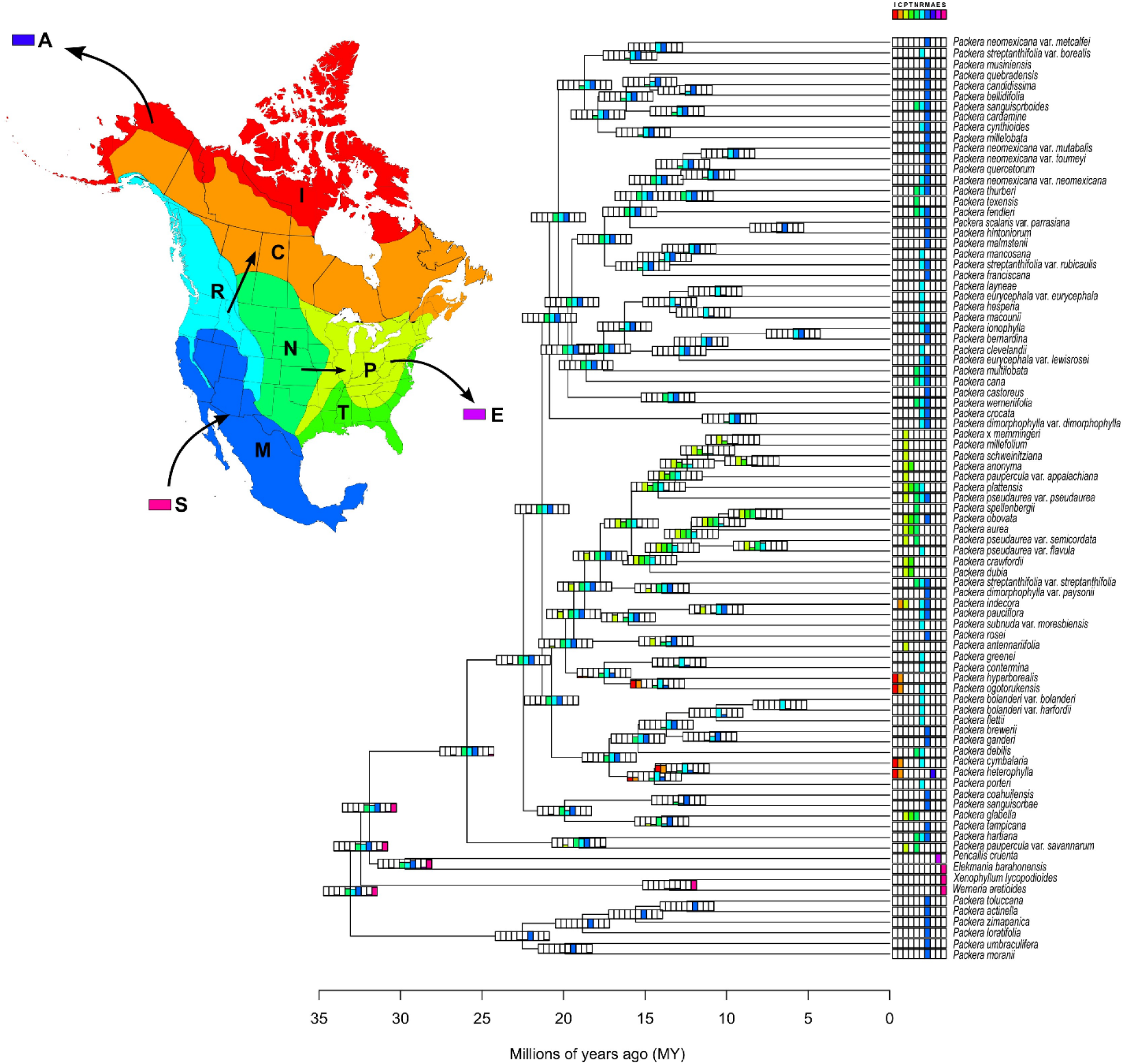
Predicted biogeographic regions generated from BioGeoBEARS on all *Packera* taxa and four outgroup genera (*Pericallis*, *Elekmania*, *Werneria*, and *Xenophyllum*) using the model BAYAREALIKE. The dated, nuclear tree was generated using treePL as input. Units at the bottom of the figure represent time in millions of years (MY). Colors on the tree and the map of North America correspond to the ten geographic regions outlined by Takhtajan (1986): Arctic Province (I), Canadian Province (C), Appalachian Province (P), Atlantic and Gulf Coast Plain Province (T), North American Prairies Province (N), Rocky Mountain and Vancouvarian Province (R), Madrean Region (M), Asia (A), Europe (E), and Central America (S). The bars to the left of the taxon tip labels represent current distributions of the species. Bars at the node of each tree represent the probability of that species being within that geographic region at that specific time.

The four outgroup taxa included in the biogeography study (*Pericallis*, *Elekmania*, *Werneria*, and *Xenophyllum*) are also believed to have originated in Central America or somewhere in the western temperate regions of North America, but with a higher probability of originating in Central America. This result is consistent with the current distributions of *Elekmania*, *Werneria*, and *Xenophyllum*, which are mainly within Central America (Funk, 1997; Nordenstam, 2007). Interestingly, our results show that *Pericallis* also likely originated in Central America or the Madrean region in North America around 30MY. Considering that all *Pericallis* species are in the Canary Islands of Spain (Jones et al., 2014), a relatively young chain of islands (∼14.3MY; McDougall & Schmincke, 1976; Geldmacher et al., 2000), these results imply that early *Pericallis* originated in the Americas then had an independent, long-distance dispersal and established in Europe once the Canary Islands emerged.

A detailed discussion on species relationships within *Packera* and remaining Senecioneae taxa can be found in Supplemental Materials.

## 5 Conclusion

Our study provides the largest and most completely sampled *Packera* nuclear and plastid/plastid phylogenies to date. Even so, the nuclear and plastid phylogenies produced are highly incongruent, with many species’ relationships differing dependent on sequence type. The plastid data had low sequence diversity and the resulting phylogeny had many areas of low support. The nuclear tree shows higher support overall, though some key areas of low resolution were recovered. For those species relationships that showed high support, many of them followed interesting correlations with geohistorical events of the mid-Tertiary resulting in predictable biogeographic patterns. Given the extensive hybridization and introgression reported in this genus, we hypothesize that areas of the phylogeny with low support may be due to various evolutionary processes including hybridization, introgression, and polyploidy. Our study recovers an older age for *Packera* than other studies have reported with *Packera* originating roughly 19.2MY – 25.9MY (RelTime vs. treePL, respectively), during the late Oligocene to early Miocene. Though these dates differ from prior estimates (∼7-20MY; Pelser et al., 2010; Huang et al., 2016; Mandel et al., 2019; Zhang et al., 2021), they align well with previous geohistorical predictions that *Packera* speciation and diversification events align with the changing geography and climate in North America, providing further evidence that these historical events have played a significant role in *Packera*’s diversification and biogeographical distributions. Finally, our study shows that *Packera* likely originated in western United States or Mexico, then diversified further north and east when climates changed, or sea levels lowered. These results provide a foundation for future studies wanting to taxonomically revise or develop more detailed studies on diversifications and evolutionary histories in this complex genus.

## Acknowledgments

We would like to thank Dr. Carol Siniscalchi and Matthew Pollard for their bioinformatic help, Drs. Randall Bayer, Duane McKenna, and Linda Watson for reading and editing prior to submission, and John Bain, Ed Schilling, and Brandon Fuller for donating genetic material for this study. We would also like to thank the following herbaria for allowing us to visit and destructively sample leaf tissue: ALA, ARIZ, ASU, CAS, MEM, NY, OSC, TEX, UNC, UCR, WTU. Permits to collect in the San Bernardino National Forest and Shasta-Trinity National Forest in California were obtained prior to collection of specimens by contacting the USDA Forest Service. This work was funded by the Center for Biodiversity (CBio) Research at the University of Memphis.

ERMP collected leaf tissue, performed wet-lab and bioinformatic work, wrote manuscript, and edited the manuscript. JRM helped with bioinformatics, provided suggestions, and edited the manuscript.

The authors declare no conflict of interest.

## Supplemental Materials

Discussion on species relationships within *Packera* and related Senecioneae taxa.

**nuc_alignments.phy** Nuclear supermatrix file.

**chl_alignments.phy** Plastid supermatrix file.

**Supplemental Table 1**. Voucher specimens of the 108 taxa used in this study. Publication status and authorities assigned by IPNI.

**Supplemental Table 2**. Raw sequence statistics using Phyx for each of the samples used in this study.

**Supplemental Table 3**. Matrix stats obtained by AMAS and stats about the number of loci and loci choice of the nuclear and plastid phylogenies.

**Supplemental Table 4**. Presence/Absence values of each species given the ten geographic regions outlined by Takhtajan (1986). A value of “0” indicates absense, “1” indicates presence. Letters to the right of each column heading relates to Figure 6. The four column headings above the geographic regions refer to the four groupings of *Packera* used in the discussion, no groupings were designated four Europe or Central America since they represent outgroup distributions.

**Supplemental Fig. 1**. Nuclear phylogeny produced using a concatenation approach and inferred with maximum likelihood (RAxML). Scale bar under the tree on the left represents the mean number of nucleotide substitutions per site. Support values (bootstraps) are indicated next to the node. Phylogeny generated using phytools in R.

**Supplemental Fig. 2**. Plastome phylogeny produced using a pseudocoalescent analysis (ASTRAL-III) and inferred with maximum likelihood (RAxML). ASTRAL-III reported that the plastid tree has a normalized quartet score of 0.526, meaning that 53% of the gene trees match the final species tree. Scale bar under the tree on the left represents the mean number of nucleotide substitutions per site. Support values (LPP) are next to the node. Phylogeny generated using *phytools* in R.

**Supplemental Fig. 3**. Figure 2 nuclear phylogeny produced using a pseudocoalescent analysis (ASTRAL-III) and inferred with maximum likelihood (RAxML), with all support values (LPP) indicated next to node. Scale bar on bottom left represents the mean number of nucleotide substitutions per site. Phylogeny generated using *phytools* in R.

**Supplemental Fig. 4**. Figure 3 plastid phylogeny produced using a concatenated approach, inferred with maximum likelihood (RAxML), with all support values (bootstrap) indicated next to node. Branch lengths were transformed so that node values can be more easily seen. Scale bar on bottom left represents the mean number of nucleotide substitutions per site. Phylogeny generated using *phytools* in R.

**Supplemental Fig. 5**. Tanglegram comparing taxon placement between nuclear (left) and plastid (right) genome sequences. The nuclear tree was produced using a pseudo-coalescent analysis (ASTRAL-III), while the plastid tree was produced using a concatenated matrix. Both nuclear and plastid trees were then visualized and inferred with maximum likelihood (RAxML) approaches and visualized in R. Support values (LPP [nuclear] and BS [plastid]) of 0.9LPP/90BS or greater are indicated with a blue diamond at the node. Dashed and solid lines represent correspondence between taxa in nuclear and plastid phylogenies: **solid line** – nuclear and plastome relationships agree, **dashed line** – nodes conflict, **red-colored dashed line** – major node conflict. Core *Packera* species’ tips are collapsed and indicated by a teal font.

**Supplemental Figs. 6 – 13**. Divergence rate estimates of Scenarios 1-8 using treePL in millions of years. Left tree contains the mean age of that designated node, the right tree shows the 95% age range for the designated node.

**Supplemental Figs. 14 – 21**. Divergence rate estimates of Scenarios 1-8 using RelTime in millions of years. Left tree contains the mean age of that designated node, the right tree shows the 95% age range for the designated node.

**Supplemental Figs. 22 & 23**. BioGeoBEARS results using DEC. Boxes at nodes represent the geographic regions, and pie charts show the probability of the geographic regions provided in Figure 6.

**Supplemental Figs. 24 & 25**. BioGeoBEARS results using DEC+*J*. Boxes at nodes represent the geographic regions, and pie charts show the probability of the geographic regions provided in Figure 6.

**Supplemental Figs. 26 & 27**. BioGeoBEARS results using DIVALIKE. Boxes at nodes represent the geographic regions, and pie charts show the probability of the geographic regions provided in Figure 6.

**Supplemental Figs. 28 & 29**. BioGeoBEARS results using DIVALIKE+*J*. Boxes at nodes represent the geographic regions, and pie charts show the probability of the geographic regions provided in Figure 6.

**Supplemental Figs. 30 & 31**. BioGeoBEARS results using BAYAREALIKE. Boxes at nodes represent the geographic regions, and pie charts show the probability of the geographic regions provided in Figure 6.

**Supplemental Figs. 32 & 33**. BioGeoBEARS results using BAYEAREALIKE+*J*. Boxes at nodes represent the geographic regions, and pie charts show the probability of the geographic regions provided in Figure 6.

## Notes

### Competing Interest Statement

The authors have declared no competing interest.

## References

1. Alix K, Gérard PR, Schwarzacher T, Heslop-Harrison JS. 2017. Polyploidy and interspecific hybridization: Partners for adaptation, speciation and evolution in plants. Annals of Botany 120(2): 183–194.

2. Bain JF, Jansen RK. 1995. A phylogenetic analysis of the aureoid *Senecio* (Asteraceae) complex based on ITS sequence data. Plant Systematics and Evolution 195(3–4): 209–219.

3. Bain JF, Jansen RK. 1996. Numerous chloroplast DNA polymorphisms are shared among different populations and species in the aureoid *Senecio* (*Packera*) complex. Canadian Journal of Botany 74(11): 1719–1728.

4. Bain JF, Tyson BS, Bray DF. 1997. Variation in pollen wall ultrastructure in New World Senecioneae (Asteraceae), with special reference to *Packera*. Canadian Journal of Botany 75(5): 730–735.

5. Bain JF, Golden JL. 2000. A phylogeny of *Packera* (Senecioneae; Asteraceae) based on internal transcribed spacer region sequence data and a broad sampling of outgroups. Molecular Phylogenetics and Evolution 16(3): 331–338.

6. Bain JF, Golden JL. 2005. Chloroplast haplotype diversity patterns in *Packera pauciflora* (Asteraceae) are affected by geographical isolation, hybridization, and breeding system. Canadian Journal of Botany 83(8): 1039–1045.

7. Bain JF, Jansen RK. 2006. A chloroplast DNA hairpin structure provides useful phylogenetic data within tribe Senecioneae (Asteraceae). Canadian Journal of Botany 84(5): 862–868.

8. Bankevich A, Nurk S, Antipov D, Gurevich AA, Dvorkin M, Kulikov AS, Lesin VM, Nikolenko SI, Pham S, Prjibelski AD, Pyshkin AV, Sirotkin AV, Vyahhi N, Tesler G, Alekseyev MA, Pevzner PA. 2012. SPAdes: A new genome assembly algorithm and its applications to single-cell sequencing. Journal of Computational Biology 19(5): 455–477.

9. Barba-Montoya J. 2021. GBE Assessing Rapid Relaxed-Clock Methods for Phylogenomic Dating. 13: 1–14.

10. Barkley TM. 1978. Three new species of *Senecio* (Asteraceae) from Mexico. Brittonia 30: 69– 75.

11. Barkley TM. 1985. Infrageneric groups in *Senecio*, S. L., and *Cacalia*, S. L. (Asteraceae: Senecioneae) in Mexico and Central America. Brittonia 37(2): 211–218.

12. Barkley TM. 1988. Variation among the Aureoid *Senecios* of North America: A geohistorical interpretation. The Botanical Review 54(1): 82–106.

13. Bayer RJ. 1989a. A taxonomic revision of the *Antennaria rosea* (Asteraceae: Inuleae: Gnaphaliinae) polyploid complex. Brittonia 41(1): 53–60.

14. Bayer RJ. 1989b. Patterns of isozyme variation in the *Antennaria rosea* (Asteraceae: Inuleae) polyploid agamic complex. Systematic Botany 14(3): 389–397.

15. Bayer RJ, Soltis DE, Soltis PS. 1996. Phylogenetic inferences in *Antennaria* (Asteraceae: Gnaphalieae: Cassiniinae) based on sequences from nuclear ribosomal DNA internal transcribed spacers (ITS). American Journal of Botany 83: 516–527.

16. Benton MJ, Donoghue PCJ. 2007. Paleontological evidence to date the tree of life. Molecular Biology and Evolution 24: 26–53.

17. Bolger AM, Lohse M, Usadel B. 2014. Trimmomatic: A flexible trimmer for Illumina sequence data. Bioinformatics 30: 2114–2120.

18. Bouckaert R, Vaughan TG, Barido-Sottani J, Duchêne S, Fourment M, Gavryushkina A, Heled J, Jones G, Kühnert D, De Maio N, Matschiner M, Mendes FK, Müller NF, Ogilvie HA, Du Plessis L, Popinga A, Rambaut A, Rasmussen D, Siveroni I, Suchard MA, Wu CH, Xie D, Zhang C, Stadler T, Drummond AJ. 2019. BEAST 2.5: An advanced software platform for Bayesian evolutionary analysis. PLoS Computational Biology 15 :1–28.

19. Boufford DE, Kartesz JT, Shi S, Zhou R. 2014. *Packera serpenticola* (Asteraceae; Senecioneae), a New Species from North Carolina, U. S. A. Systematic Botany 39: 1027–1030.

20. Brown JW, Walker JF, Smith SA. 2017. Phyx: Phylogenetic tools for unix. Bioinformatics 33: 1886–1888.

21. Costa FP, Schrago CG, Mello B. 2022. Accessing the relative performance of fast molecular dating methods for phylogenomic data. Research Square 1–14.

22. Elven R, Murray D, Razzhivin V, Yurtsev BA. 2011. Annotated Checklist of the Panarctic Flora (PAF) Vascular plants version 1.0. *Available at* http://panarcticflora.org/results?biogeographic=&bioclimatic=&region=&name=packera+heterophylla#paf-863101 (accessed May 17, 2022).

23. Felsenstein J. 1978. Cases in which parsimony or compatibility methods will be positively misleading. Systematic Zoology 27(4): 401–410.

24. Folk RA, Freudenstein JV. 2014. Phylogenetic relationships and character evolution in *Heuchera* (Saxifragaceae) on the basis of multiple nuclear loci. American Journal of Botany 101(9): 1532–1550.

25. Folk RA, Mandel JR, Freudenstein JV. 2015. A protocol for targeted enrichment of intron containing sequence markers for recent radiations: A phylogenomic example from *Heuchera* (Saxifragaceae). Applications in Plant Sciences 3: 1–10.

26. Freeman CC, Barkley TM. 1995. A synopsis of the genus *Packera* (Asteraceae: Senecioneae) in Mexico. *SIDA*, Contributions to Botany 16: 699–709.

27. Funk VA, Susanna A, Stuessy TF, Bayer RJ (eds.). 2009. Systematics, Evolution, and Biogeography of Compositae. Vienna: International Association for Plant Taxonomy.

28. Geldmacher J, Van Den Bogaard P, Hoernle K, Schmincke HU. 2000. The 40Ar/39Ar age dating of the Madeira Archipelago and hotspot track (eastern North Atlantic). *Geochemistry, Geophysics*, Geosystems 1(2).

29. Golden JL, Bain JF. 2000. Phylogeographic patterns and high levels of chloroplast DNA diversity in four *Packera* (Asteraceae) species in Southwestern Alberta. Evolution 54:1566– 1579.

30. Golden JL, Kim YD, Bain JF. 2001. A re-evaluation of North American *Tephroseris* and *Sinosenecio* (Asteraceae: Senecioneae) based on molecular and micromorphological data. Canadian Journal of Botany 79: 1195–1201.

31. Gramling A. 2006. A conservation assessment of Packera millefolium, a Southern Appalachian Endemic. University of North Caroline at Chapel Hill.

32. Gray A, Torrey J. 1843. 163. Senecio Linn. In: A flora of North America: containing abridged descriptions of all the known indigenous and naturalized plants growing north of Mexico, arranged according to the natural system. 436–446.

33. Gray A. 1886. Synoptical Flora of North America: The Gamopetalae, Vol I, Part II and Vol II, Part I. New York: Ivison, Blackeman, Taylor, and Co. 383–394.

34. Greenman JM. 1907. New species of *Senecio* and *Schoenocaulon* from Mexico. Contributions from the Gray Herbarium of Harvard University 34: 19–21.

35. Healey A, Furtado A, Cooper T, Henry RJ. 2014. Protocol: A simple method for extracting next generation sequencing quality genomic DNA from recalcitrant plant species. Plant Methods 10: 1–8.

36. Hobbs CR, Baldwin BG. 2013. Asian origin and upslope migration of Hawaiian *Artemisia* (Compositae-Anthemideae). Journal of Biogeography 40: 442–454.

37. Huang CH, Zhang C, Liu M, Hu Y, Gao T, Qi J, Ma H. 2016. Multiple polyploidization events across Asteraceae with two nested events in the early history revealed by nuclear phylogenomics. Molecular Biology and Evolution 33: 2820–2835.

38. Janovec JP, Barkley TM. 1996. *Sinosenecio newcombei* (Asteraceae : Senecioneae): A new combination for a North American plant in an Asiatic genus. Novon 6: 265–267.

39. Jansen RK, Michael HK, Wallace RS, Kim K-J, Keeley SC, Watson LE, Palmer JD. 1992. Chloroplast DNA variation in the Asteraceae: phylogenetic and evolutionary implications. In: Molecular systematics of plants. Springer, Boston, MA., 252–279.

40. Janssens SB, Couvreur TLP, Mertens A, Dauby G, Dagallier L-PMJ, Abeele S Vanden, Vandelook F, Mascarello M, Beeckman H, Sosef M, Droissart V, van der Bank M, Maurin O, Hawthorne W, Marshall C, Réjou-Méchain M, Beina D, Baya F, Merckx V, Verstraete B, Hardy O. 2020. A large-scale species level dated angiosperm phylogeny for evolutionary and ecological analyses. Biodiversity Data Journal 8.

41. Jeffrey C. (1992). The tribe Senecioneae (Compositae) in the Mascarene Islands with an annotated world check-list of the genera of the tribe: Notes on Compositae: VI. Kew Bulletin 47(1): 49.

42. Johnson MG, Gardner EM, Liu Y, Medina R, Goffinet B, Shaw AJ, Zerega NJC, Wickett NJ. 2016. HybPiper: Extracting coding sequence and introns for phylogenetics from high throughput sequencing reads using target enrichment. Applications in Plant Sciences 4: 1600016.

43. Jones KE, Alfredo Reyes-Betancort J, Hiscock SJ, Carine MA. 2014. Allopatric diversification, multiple habitat shifts, and hybridization in the evolution of *Pericallis* (Asteraceae), a Macaronesian endemic genus. American Journal of Botany 101: 637–651.

44. Jones KE, Fér T, Schmickl RE, Dikow RB, Funk VA, Herrando-Moraira S, Johnston PR, Kilian N, Siniscalchi CM, Susanna A, Slovák M, Thapa R, Watson LE, Mandel JR. 2019. An empirical assessment of a single family-wide hybrid capture locus set at multiple evolutionary timescales in Asteraceae. Applications in Plant Sciences 7: 1–27.

45. Katoh K, Standley DM. 2013. MAFFT multiple sequence alignment software version 7: Improvements in performance and usability. Molecular Biology and Evolution 30: 772–780.

46. Kowal RR, Judziewicz EJ, Edwards J. 2011. *Packera insulae-regalis* (Asteraceae, Senecioneae), a new species endemic to Isle Royale, Michigan, U.S.A. Brittonia 63: 343–354.

47. Kück P, Longo GC. 2014. FASconCAT-G: Extensive functions for multiple sequence alignment preparations concerning phylogenetic studies. Frontiers in Zoology 11: 1–8.

48. Kumar S, Stecher G, Li M, Knyaz C, Tamura K. 2018. MEGA X: Molecular evolutionary genetics analysis across computing platforms. Molecular Biology and Evolution 35: 1547– 1549.

49. Kumar S, Stecher G, Peterson D, Tamura K. 2012. MEGA-CC: Computing core of molecular evolutionary genetics analysis program for automated and iterative data analysis. Bioinformatics 28: 2685–2686.

50. Kumar S, Stecher G, Tamura K. 2016. MEGA7: Molecular Evolutionary Genetics Analysis Version 7.0 for Bigger Datasets. Molecular biology and evolution 33: 1870–1874.

51. Landis MJ, Matzke NJ, Moore BR, Huelsenbeck JP. 2013. Bayesian analysis of biogeography when the number of areas is large. Systematic Biology 62: 789–804.

52. Lanfear R, Calcott B, Ho SYW, Guindon S. 2012. PartitionFinder: Combined selection of partitioning schemes and substitution models for phylogenetic analyses. Molecular Biology and Evolution 29: 1695–1701.

53. Langmead B, Salzberg SL. 2012. Fast gapped-read alignment with Bowtie 2. Nature Methods 9: 357–359.

54. Langmead B, Wilks C, Antonescu V, Charles R. 2019. Scaling read aligners to hundreds of threads on general-purpose processors. Bioinformatics 35: 421–432.

55. Li H, Durbin R. 2009. Fast and accurate short read alignment with Burrows-Wheeler transform. Bioinformatics 25: 1754–1760.

56. Li H, Handsaker B, Wysoker A, Fennell T, Ruan J, Homer N, Marth G, Abecasis G, Durbin R. 2009. The Sequence Alignment/Map format and SAMtools. Bioinformatics 25: 2078–2079.

57. Li HT, Yi TS, Gao LM, Ma PF, Zhang T, Yang JB, Gitzendanner MA, Fritsch PW, Cai J, Luo Y, Wang H, van der Bank M, Zhang SD, Wang QF, Wang J, Zhang ZR, Fu CN, Yang J, Hollingsworth PM, Chase MW, Soltis DE, Soltis PS, Li DZ. 2019. Origin of angiosperms and the puzzle of the Jurassic gap. Nature Plants 5: 461–470.

58. Löve Á, Löve Á. 1976. Nomenclatural notes on arctic plants. Botaniska Notiser 128: 497–523.

59. Mahoney AM. 2000. Contributions to the systematics of the *Packera paupercula* complex (Asteraceae: Senecioneae). Ph.D. Dissertation. University of Wisconsin-Madison.

60. Mahoney AM, Kowal RR. 2008. Three new varieties of *Packera paupercula* (Asteraceae, Senecioneae) in midwestern and southeastern North America. Novon 18: 220–228.

61. Mandel JR, Dikow RB, Funk VA, Masalia RR, Staton SE, Kozik A, Michelmore RW, Rieseberg LH, Burke JM. 2014. A target enrichment method for gathering phylogenetic information from hundreds of loci: An example from the Compositae. Applications in Plant Sciences 2: 1300085.

62. Mandel JR, Dikow RB, Siniscalchi CM, Thapa R, Watson LE, Funk VA. 2019. A fully resolved backbone phylogeny reveals numerous dispersals and explosive diversifications throughout the history of Asteraceae. Proceedings of the National Academy of Sciences of the United States of America 116: 14083–14088.

63. Matzke NJ. 2013. BioGeoBEARS: Biogeography with Bayesian (and likelihood) evolutionary analysis in R scripts. R package, version 0.2. 1, 2013.

64. Maurin KJL. 2020. An empirical guide for producing a dated phylogeny with treePL in a maximum likelihood framework. arXiv: 2008.07054.

65. Maurin KJL, Smissen RD, Lusk CH. 2022. A dated phylogeny shows Plio-Pleistocene climates spurred evolution of antibrowsing defences in the New Zealand flora. New Phytologist 233:546–554.

66. McDougall I, Schmincke HU. 1976. Geochronology of Gran Canaria, Canary Islands: Age of shield building volcanism and other magmatic phases. Bulletin Volcanologique 40: 57–77.

67. Mello B. 2018. Estimating timetrees with MEGA and the timetree resource. Molecular Biology and Evolution 35: 2334–2342.

68. Mello B, Tao Q, Tamura K, Kumar S. 2017. Fast and accurate estimates of divergence times from big data. Molecular Biology and Evolution 34: 45–50.

69. Meseguer AS, Lobo JM, Ree R, Beerling DJ, Sanmartín I. 2015. Integrating fossils, phylogenies, and niche models into biogeography to reveal ancient evolutionary history: The case of *Hypericum* (Hypericaceae). Systematic Biology 64: 215–232..

70. Mohlenbrock RH. 2004. Validation of new combinations of Vascular Plants. Phytoneuron 67 :1– 3.

71. Müller J, Reisz RR. 2005. Four well-constrained calibration points from the vertebrate fossil record for molecular clock estimates. BioEssays 27: 1069–1075.

72. Nordenstam B, Pelser PB. 2011. Notes on the generic limits of *Sinosenecio* and *Tephroseris* (Compositae - Senecioneae). Comp. Newsl. 49: 1–7.

73. Ornduff R, Mosquin T, Kyhos DW, Raven PH. 1967. Chromosome numbers in Compositae. VI. Senecioneae. II. American Journal of Botany 54: 205–213.

74. Panero JL, Funk VA. 2008. The value of sampling anomalous taxa in phylogenetic studies: Major clades of the Asteraceae revealed. Molecular Phylogenetics and Evolution 47: 757– 782.

75. Parham JF, Donoghue PCJ, Bell CJ, Calway TD, Head JJ, Holroyd PA, Inoue JG, Irmis RB, Joyce WG, Ksepka DT, Patané JSL, Smith ND, Tarver JE, Van Tuinen M, Yang Z, Angielczyk KD, Greenwood JM, Hipsley CA, Jacobs L, Makovicky PJ, Müller J, Smith KT, Theodor JM, Warnock RCM, Benton MJ. 2012. Best practices for justifying fossil calibrations. Systematic Biology 61: 346–359.

76. Pease JB, Brown JW, Walker JF, Hinchliff CE, Smith SA. 2018. Quartet Sampling distinguishes lack of support from conflicting support in the green plant tree of life. American Journal of Botany 105: 385–403.

77. Pelser PB, Nordenstam B, Kadereit JW, Watson LE. 2007. An ITS phylogeny of tribe Senecioneae (Asteraceae) and a new delimitation of *Senecio* L. Taxon 56: 1077–1104.

78. Pelser PB, Kennedy AH, Tepe EJ, Shidler JB, Nordenstam B, Kadereit JW, Watson LE. 2010. Patterns and causes of incongruence between plastid and nuclear Senecioneae (Asteraceae) phylogenies. American Journal of Botany 97: 856–873.

79. Portik DM, Wiens JJ. 2020. Do alignment and trimming methods matter for phylogenomic (UCE) analyses? Systematic Biology.

80. Quinlan EJ, Mathews KG, Collins B, Young R. 2020. Phylogenetic divergence and ecophysiological variation in the disjunct *Kalmia buxifolia* (Sand-myrtle, Ericaceae). Systematic Botany 45: 900–912.

81. R Core Team. 2016. R: A language and environment for statistical computing. R Foundation for Statistical Computing, Vienna, Austria.

82. Rebman JP, Gibson J, Rich K. 2016. Annotated Checklist of the Vascular Plants of Baja California, Mexico.

83. Ree RH, Smith SA. 2008. Maximum likelihood inference of geographic range evolution by dispersal, local extinction, and cladogenesis. Systematic Biology 57:4–14. DOI: 10.1080/10635150701883881.

84. Revell LJ. 2012. phytools: An R package for phylogenetic comparative biology (and other things). Methods in Ecology and Evolution 3: 217–223.

85. Rieseberg LH, Willis JH. 2007. Plant speciation. Science 317: 910–914.

86. Ronquist F. 1997. Dispersal-vicariance analysis: A new approach to the quantification of historical biogeography. Systematic Biology 46:195–203.

87. RStudio. (2020). RStudio: Integrated Development for R. RStudio, PBC. http://www.rstudio.com/

88. Śarhanov P, Sharbel TF, Sochor M, Vaśut RJ, Danćák M, Trávníćek B. 2017. Hybridization drives evolution of apomicts in *Rubus* subgenus *Rubus*: Evidence from microsatellite markers. Annals of Botany 120: 317–328.

89. Sayyari E, Mirarab S. 2016. Fast coalescent-based computation of local branch support from quartet frequencies. Molecular Biology and Evolution 33: 1654–1668.

90. Schilling EE. 2001. Phylogeny of *Helianthus* and related genera. OLC 8: 22–25.

91. Schilling EE, Floden A. 2015. Barcoding the Asteraceae of Tennessee, tribe Cichorieae. Phytoneuron 19: 1–8.

92. Schilling EE, Floden AJ, Weakley AS, Winder C, Small RL. 2022. Molecular barcoding reveals unexpected diversity in eastern North American stitchworts (Caryophyllaceae). Botanical Journal of the Linnean Society: 1–10.

93. Stamatakis A. 2014. RAxML version 8: A tool for phylogenetic analysis and post-analysis of large phylogenies. Bioinformatics 30: 1312–1313.

94. Stanley SM. 1986. Earth and life through time. W. H. Freeman, New York City.

95. Stephens JD, Rogers WL, Mason CM, Donovan LA, Malmberg RL. 2015. Species tree estimation of diploid *Helianthus* (Asteraceae) using target enrichment. American Journal of Botany 102: 910–920.

96. Stubbs RL, Folk RA, Soltis DE, Cellinese N. 2020. Diversification in the Arctic: Biogeography and systematics of the North American *Micranthes* (Saxifragaceae). Systematic Botany 45: 802–811.

97. Takhtajan A. 1986. Floristic Regions of the World. Berkley and Los Angeles, California: University of California Press.

98. Tao Q, Tamura K, Kumar S. 2020. The Molecular Evolutionary Clock. The Molecular Evolutionary Clock.

99. Thapa R, Bayer RJ, Mandel JR. 2020. Phylogenomics resolves the relationships within *Antennaria* (Asteraceae, Gnaphalieae) and yields new insights into its morphological character evolution and biogeography. Systematic Botany 45: 387–402.

100. Thapa R, Bayer RJ, Mandel JR. 2021. Genetic diversity, population structure, and ancestry estimation in the *Antennaria rosea* (Asteraceae: Gnaphalieae) polyploid agamic complex. Taxon 70: 139–152.

101. Tillich M, Lehwark P, Pellizzer T, Ulbricht-Jones ES, Fischer A, Bock R, Greiner S. 2017. GeSeq - Versatile and accurate annotation of organelle genomes. Nucleic Acids Research 45: W6–W11.

102. Timme RE, Kuehl JV, Boore JL, Jansen RK. 2007. A comparative analysis of the *Lactuca* and *Helianthus* (Asteraceae) plastid genomes: Identification of divergent regions and categorization of shared repeats. American Journal of Botany 94: 302–312.

103. Trock DK. 1999. A revisionary synthesis of the genus *Packera* (Asteraceae: Senecioneae). Ph.D. Dissertation. Kansas State University.

104. Trock DK. 2006. *Packera* Á. Löve & D. Löve. In: Flora of North America Editorial Committee*, eds. 1993+*. New York and Oxford, 570–602.

105. Uhrinová V, Zozomová-Lihová J, Bernátová D, Paule J, Paule L, Gömöry D. 2017. Origin and genetic differentiation of pink-flowered *Sorbus* hybrids in the Western Carpathians. Annals of Botany 120: 271–284.

106. Walker JD, Geissman JW, Bowring SA, Babcock LE, compilers. (2018). Geologic Time Scale v. 5.0: Geological Society of America. ©2018 The Geological Society of America Wang WM. 2004. On the origin and development of *Artemisia* (Asteraceae) in the geological past. Botanical Journal of the Linnean Society 145: 331–336.

107. Weber WA, Löve Á. 1981. New combinations in the genus *Packera* (Asteraceae). Phytologia 49: 44–50.

108. Yeatts L, Schneider A, Schneider B. 2011. *Packera mancosana* (Asteraceae: Senecioneae), a new species and shale barren endemic of southwestern colorado. Phytoneuron: 1–8.

109. Zhang C, Huang CH, Liu M, Hu Y, Panero JL, Luebert F, Gao T, Ma H. 2021. Phylotranscriptomic insights into Asteraceae diversity, polyploidy, and morphological innovation. Journal of Integrative Plant Biology 63: 1273–1293.

110. Zhang C, Rabiee M, Sayyari E, Mirarab S. 2018. ASTRAL-III: Polynomial time species tree reconstruction from partially resolved gene trees. BMC Bioinformatics 19: 15–30.

