## Supplementary material for "Resolving evolutionary relationships in the groundsels: phylogenomics, divergence time estimates, and biogeography of *Packera* (Asteraceae: Senecioneae)": Moore-Pollard_Mandel_SupplementalMaterials.docx

**Supplemental Materials**

Further discussion on evolutionary relationships in *Packera* will follow four groupings of the ten floristic provinces/regions of North America outlined in Supplemental Table 4, with the following modifications: West Coast United States and Mexico (Rocky Mountain and Vancouvarian Province and the Madrean Region), Arctic/Alpine (Arctic Province, Canadian Province, and Asia), East Coast (Appalachian Province and Atlantic and Gulf Coast Plain Province), and Central North America (North American Prairies Province). Additionally, a detailed discussion of the remaining Senecioneae taxa can be found following the *Packera* discussion below.

**Species relationships in core *Packera***

West Coast United States and Mexico

Most species endemic to the state of California are found in a single clade, hereafter referred to as the “Californian endemic clade”, with the exception of *P. brewerii*, *P. ganderi*, and *P. greenei* which are found in more distant clades (Figs. 2 & 6). Both *P. ganderi* and *P. brewerii* are Californian endemics, with *P. ganderi* found only in the mountain ranges of San Diego County, and *P. brewerii* found along California’s coastal ranges from San Francisco Bay County to Los Angeles County. These species are the southernmost Californian endemics, apart from *P. bernardina* and *P. ionophylla* which are found almost exclusively on mountain tops in southern California. *Packera bernardina* is found only in the San Bernardino mountains, while *P. ionophylla* is found in San Bernardino, San Gabriel, and Tehachapi mountains, all of which are slightly more north of the San Diego County mountains, and more inland than the coast (Trock, 1999). Even though these species are close geographically, our phylogeny shows that they are distantly related, with *P. bernardina* and *P. ionophylla* placing in the Californian endemic clade and *P. ganderi* and *P. brewerii* showing closer relationships with Arctic/Alpine species in a separate clade (i.e., *P. bolanderi* var. *bolanderi*, *P. bolanderi* var. *harfordii*, and *P. flettii*; Fig. 6). *Packera greenei* is also an endemic Californian species and is only found on the coastal ranges in northern California from Napa to Trinity counties. It is placed separately from the Californian endemic clade, and along with *P. contermina*, *P. hyperborealis*, and *P. ogotorukensis*, which are distributed through central to northern North America.

Although they are geographically isolated from each other, other Californian endemic species in this clade form a smaller clade comprised of *Packera eurycephala* var. *lewisrosei*, *P. clevelandii*, *P. ionophylla*, *and P. bernardina* (Fig. 6)*.* *Packera eurycephala* var. *lewisrosei* and *P. clevelandii* which are found further north, closer to Sacramento, California. The other species found in this clade are not endemic to, but have a centralized distribution in California. Examples of this include *P. layneae*, *P. eurycephala* var. *eurycephala*, *P. hesperia*, and *P. macounii*, whose distributions stretch from Mexico to Washington state. More widespread species, such as *P. cana* and *P. multilobata*, are not centralized in California but are largely distributed throughout the state.

Mexico and Southwestern United States (US) species are polyphyletic with most taxa found in four clades, two of which are distantly separated from the other two, and only one Mexican endemic species, *Packera rosei*, was placed outside of those clades (Fig. 6). The first mainly Mexico and Southwestern US clade is almost exclusively Mexican and Southwestern US taxa, apart from *P. streptanthifolia* var. *borealis* and *P. musiniensis*, which are found solely in Central North America (NA) and do not appear to overlap with any other taxa in this clade. These two taxa place within the same smaller clade, though they are not sister to each other: *P. streptanthifolia* var. *borealis* is sister to *P. neomexicana* var. *metcalfei* (found exclusively in the mountains along Mogollon Rim in Arizona and New Mexico), which is sister to *P. musiniensis* (Fig. 6).

Previous phylogenetic work found that *Packera multilobata*, *P. sanguisorboides*, *P. neomexicana*, and *P. bellidifolia* are genetically similar and grouped together (Bain & Jansen, 1995). Bain & Golden (2000) found similar results to Bain & Jansen (1995); however, their phylogeny also included *P. hartiana* and *P. quercetorum* in the smaller clade. Our phylogeny follows these trends, apart from *P. multilobata* and *P. hartiana* which are placed in separate clades. Bain & Jansen (1995) also commented that they would have expected *P. millelobata* to group with these taxa, which is supported in our phylogeny (Fig. 2).

One of the strongest relationships in the Mexican and Southwestern US clades was between *Packera candidissima* and *P. bellidifolia* (1.0LPP; Supplemental Fig. 3), which was expected considering they are morphologically very similar, have similar distributions, and share many alkaloid properties (Bah et al., 1994; Trock, 1999; Vivar et al., 2007; Fragoso-Serrano et al., 2012). Distinctions between *P. candidissima* and *P. bellidifolia* are weak, causing some researchers to speculate that they may prove to be varieties of one species instead of independent species (Trock 1999). However, our phylogeny does not provide any evidence for or against that hypothesis.

The second Mexico and Southwestern US clade is a subclade of a larger clade comprising of Mexican and Southwestern US members in one, and majority Central NA members in the other (Fig. 6). In the Mexico and Southwestern US subclade, some non-Mexico and Southwestern US members include *P. thurberi* and *P. fendleri* which have more widespread distributions and overlap with other Mexican and Southwestern US taxa in New Mexico, though their main distributions are in the Rocky Mountains of Colorado and Wyoming. The second subclade comprises four species that are distributed through Central North America and will be discussed in a later section.

The third Mexican and Southwestern US clade contains four of the five members from the Sanguisorbae alliance that was originally proposed by Freeman (1985) (Fig. 6). Three of those four taxa (*Packera sanguisorbae*, *P. tampicana*, and *P. coahuliensis*) have distributions in Mexico and/or the Southwestern US. The fourth member, *P. glabella*, has distributions in as far south as Texas, though it is mainly distributed throughout the East Coast. Even so, its morphology is drastically different than other East Coast species, so much so that botanists have always associated it more with Mexican and Southwestern US taxa, particularly with the Sanguisorbae alliance members (Barkley 1988, Freeman 1985, Trock 1999). Placement of Sanguisorbae alliance taxa within our tree support previous findings by Bain & Golden (2000) who found that *P. sanguisorbae* and *P. glabella* are “basal” (i.e., the deepest branching lineages) in *Packera*, along with *P. montereyana* (not tested). Additionally, previous ITS sequencing found that *P. glabella* and *P. tampicana* are almost completely identical (Schilling & Floden, 2015), whereas we found that *P. glabella* and *P. tampicana* are sister with high support (1.0LPP). Additionally, *P. coahuliensis* and *P. sanguisorbae* are known to closely resemble each other (Trock, 1999), and our phylogeny placed them as sisters with high support (1.0LPP; Supplemental Fig. 3).

The fourth Mexico and Southwestern US clade at crown *Packera* is comprised of two unexpected species: *Packera hartiana* and *P. paupercula* var. *savannarum*. *Packera hartiana*, a species found in Southwestern US, closely resembles *P. paupercula* and *P. plattensis*, both widespread species, but has been separated based on geography and some morphological characters (Trock, 2006). Other than geography, *P. paupercula* var. *savannarum*, *P. hartiana*, and *P. plattensis* share only one characteristic, adventitious roots and shoots (Mahoney & Kowal, 2008). Even so, it is surprising that *P. hartiana* and *P. paupercula* var. *savannarum* are sister to each other and with very high support (1.0LPP; Supplemental Fig. 3). Though unresolved, previous phylogenetic work placed *P. hartiana* with species found mostly in Mexican and Southwestern US and Central US clades (Bain & Golden, 2000), which is understandable given *P. hartiana*’s distribution.

Arctic/Alpine

United States northwestern, Alaskan, Canadian, and Russian (Asia) species, hereafter referred to as Arctic/Alpine, are found in multiple subclades within a larger clade composed of the United States East Coast and some Central North American species. One smaller clade is found within the Arctic/Alpine clades containing species that are not characterized as Arctic/Alpine: *P. rosei* and *P. antennariifolia*. *Packera rosei* is a widespread species found only in the southern Sierra Madre Occidental in Durango and Jalisco, Mexico, while *P. antennariifolia* is a species restricted to sloping areas on shale barrens along the East Coast. There are no known connections between the species, so their placement as sister to each other with high support (1.0LPP) is surprising (Supplemental Fig. 3). Noticeably, *P. antennariifolia*’s placement changed drastically depending on taxa being included or removed from the phylogeny (not shown). Additionally, if multiples of *P. antennariifolia* were used, then they placed as sister and within the East Coast clade, which is what would be predicted given its geographic distribution. However, when only one *P. antennariifolia* individual was utilized, its placement changed in various clades with high support.

Previous work has shown that five Arctic/Alpine species, *P. contermina*, *P. ogotorukensis*, *P. hyperborealis*, *P. cymbalaria*, and *P. subnuda* var. *subnuda*, have a complex taxonomic history. In 1972, Packer investigated the taxonomy between *P. contermina*, *P. hyperborealis*, *P. cymbalaria*, and *P. subnuda* var. *subnuda* (not in study) and concluded that *P. ogotorukensis* was a new and distinct species, separated from *P. contermina*, based on gross morphology, cytological data, and pollen grain morphology (Packer, 1972; Trock, 1999). Our phylogeny supports their separation since both *P. ogotorukensis* and *P. contermina* place separately within the same smaller clade. Barkley (1981) and Packer (1972) considered *P. ogotorukensis* conspecific with *P. hyperborealis* given they are strictly Alaska-Yukon endemics and morphologically pretty similar, but later determined that they were distinct species (Whitton & Bain, 1992). From this, Trock (2006) predicted that *P. ogotorukensis* and *P. hyperborealis* are most closely related, which is backed by our findings since both taxa are sister to each other with high support (1.0LPP; Supplemental Fig. 3).

*Packera cymbalaria* is also taxonomically complex and morphologically similar to multiple species. First, *P. contermina* and *P. cymbalaria* were previously considered synonymous (=*Senecio cymbalaria* Pursh) with a widespread distribution (Barkley, 1978). Eventually they were recognized as two distinct species based on morphological, geographic, and cytological differences (Packer, 1972; Whitton & Bain, 1992). Next, *P. cymbalaria* and *P. subnuda* var. *subnuda* are morphologically very similar and only differ in their ecology (Cronquist, 1955; Barkley, 1962). Even so, there is a general consensus that they remain separate species given differing chromosome counts (Packer, 1972). Then, *P. porteri* most closely resembles *P. cymbalaria*, but differs in height, flower stalks, and floral heads (Jeffrey, 1999). Additionally, *P. porteri* is found in one site in both Oregon and Washington (Trock, 1999), but has many sites in Colorado (Jeffrey, 1999), while *P. cymbalaria* is only found in Alaska and Canada (Trock, 2006). Given their morphological similarities, it is understandable that *P. porteri* and *P. cymbalaria* are within the same smaller clade with high support. It is surprising that *P. cymbalaria* and *P. porteri* are not more closely related to the other arctic/alpine species and are instead more closely related to United States west coast species (Fig. 2 & Supplemental Fig. 3). Although *P. porteri* was not tested, the most recent *Packera* phylogeny found a connection between United States west coast species, where *P. bolanderi*, *P. flettii*, *P. breweri*, and *P. ganderi* all formed a paraphyletic assemblage near the base of the tree; however, *P. cymbalaria* placed in a separate and more distant paraphyletic clade (Bain & Golden, 2000). If geography does play as large of a role in *Packera* speciation as anticipated, then that may explain the relationships in our tree since almost all taxa in that clade have distributions in California, Oregon, or Washington.

There is some taxonomic uncertainty as to whether *P. cymbalaria* and *P. heterophylla* are the same species. Both species have similar morphological characteristics and overlapping distributions (Trock, 2006; Krasnoborov, 2007), with *P. cymbalaria* distributed throughout the northern US, Canada, and Alaska and *P. heterophylla* distributed through Alaska, Canada, and Russia. It is believed that *P. cymbalaria* was discovered in Russia and named *P. heterophylla*, though most North American botanists currently recognize the species to be *P. cymbalaria* (Debra Trock, pers. comm.). Though we did not fully investigate their taxonomy, it is noteworthy that our phylogeny shows *P. cymbalaria* and *P. heterophylla* sister to each other with high support (0.98LPP; Supplemental Fig. 3). If *P. cymbalaria* and *P. heterophylla* are determined to be the same species, the valid name would be *Packera heterophylla* (Fischer) E. Wiebe, based on *Cineraria heterophylla* Fischer (Trock, 2006; Elven et al., 2011). Further investigation is still needed between these two taxa to determine their taxonomic status.

Finally, our phylogeny placed *P. pauciflora* sister to *P. indecora*, which is understandable since they are morphologically very similar, have predominantly discoid heads, and are self-fertile (mixed mating), which is not a common trait within *Packera* (Bain & Golden, 2005; Trock, 2006). Biosystematic studies have shown that even though they share these similarities, *P. pauciflora* and *P. indecora* possess distinctly different physiologies and should be maintained as distinct taxa (Bain & Whitton, 1994). *Packera pauciflora* and *P. indecora* are in different clades from other arctic and alpine taxa (e.g., *P. ogotorukensis* and *P. hyperborealis)*, which is interesting since they are found in similar habitats (Trock, 1999).

East Coast

One of the clades was comprised of almost all United States East Coast species, hereafter referred to as the East Coast clade, apart from *P. spellenbergii* which is a species endemic to two counties (Harding and Union) in New Mexico and one county (Kane) in Utah (Trock, 2006), and *P. pseudaurea* and varieties which are distributed more central to western United States. Our phylogeny shows *P. spellenbergii* sister to *P. obovata*, a very widespread species, that is morphologically and ecologically distinct from *P. spellenbergii*. The relationship between *P. spellenbergii* and *P. obovata* was not expected given how little similarity there is between the two species. *Packera spellenbergii* has typically been associated with *P. werneriifolia*, a Rocky mountain species, given their drastic similarities (Barkley, 1989; Trock, 1999); however, our phylogeny indicates *P. spellenbergii* and *P. werneriifolia* are genetically distant (Supplemental Fig. 3). All *P. pseudaurea* species and varieties (*P. pseudaurea* var. *pseudaurea*, *P. pseudaurea* var. *flavula*, and *P. pseudaurea* var. *semicordata*) are found within the East Coast clade (Fig. 6). Though these species and varieties are mainly distributed in central to western United States, they have affiliations with most east coast species (discussed below), so this placement is not surprising.

Relationships among East Coast species are highly complicated from hybridization events. Many of the species in this clade are widespread and known to hybridize with each other, such as *P. dubia* (previously known as *P. tomentosa* C.Jeffrey) which is known to hybridize with *P. anonyma*, *P. aurea*, and *P. plattensis* (Fernald, 1943; Barkley, 1962; Chapman et al., 1971), or the species are formed by known hybridization events, such as *P.* × *memmingeri* which is an accepted hybrid species resulting from a cross between the rare and threatened *P. millefolium* and widespread *P. anonyma* (Uttal, 1984; Gramling, 2006; Weakley et al., 2011). One of the most aggressive hybridizers is *P. plattensis*, which is known to hybridize with *P. aurea*, *P. anonyma*, *P. obovata*, *P. paupercula*, and *P. dubia* (Uttal, 1982), with taxonomic complications between *P. paupercula* and its varieties (Mahoney & Kowal, 2008). Additional instances of putative hybridization events of United States east coast species have been reported with *P. obovata*, *P. paupercula*, *P. schweinitziana* and *P. pseudaurea* (Barkley, 1962; Trock, 1999), almost all of which are within the East Coast clade (Supplemental Fig. 3).

One notable placement within the East Coast clade is *P. crawfordii*, which was treated as an accepted species for the first half of the twentieth century (Fernald, 1950) but was reduced to a synonym of *P. paupercula* for the last few decades (Barkley, 1962; Radford et al., 1968; Gleason & Cronquist, 1991; Trock, 1999, 2006). More recently, *P. crawfordii* has been validated as a new species given morphological, ecological, reproductive, and cytological distinctness (Kowal et al., 2015; Kowal & Mahoney, 2016). Though *P. paupercula* var. *paupercula* was not included in this study, *P. crawfordii* appears to be distantly related from other *P. paupercula* varieties (*P. paupercula* var. *appalachiana* and *P. paupercula* var. *savannarum*; Fig. 2 & Supplemental Fig. 3), providing likely evidence for its distinctiveness from *P. paupercula*.

Central North America

Central North America (NA) species are polyphyletic and spread throughout the phylogeny, indicating that geography does not seem as important for these species’ relationships. Some clades are exclusively Central NA species, though most are found within other geographic clades (i.e., *P. pseudaurea* var. *pseudaurea* and *P. multilobata* in separate clades; Fig. 6). A potential explanation for this lack of geographic relevance could be that the species are centrally located and can more easily hybridize or introgress with the taxa in other geographic regions.

One subclade within this phylogeny is comprised of four species that are distributed through Central NA. Three of those taxa, *Packera malmstenii*, *P. mancosana*, and *P. streptanthifolia* var. *rubicaulis*, have distributions within Central NA, though their ranges do not overlap. The fourth species, *P. franciscana* is better classified as a Southwestern US species since it is only known from the San Francisco Peaks in Arizona (Fig. 6). Although *P. malmstenii* and *P. mancosana* have not been directly compared, both species have obvious affiliations with *P. werneriifolia* (Trock, 1999; Yeatts et al., 2011). *Packera malmstenii* and *P. werneriifolia* appear morphologically similar, and it is thought that further research into *P. malmstenii* may provide insight as to why *P. werneriifolia* is so variable within its range (Barkley, 1978). *Packera mancosana* is a newly described species that is similar to *P. werneriifolia* in appearance and ecology, but differs in general habitat and soil type (Yeatts et al., 2011). Some botanists that have seen *P. mancosana* in the field believe that it is simply a variable *P. werneriifolia* (Debra Trock, pers. comm.); however, our tree shows *P. mancosana* and *P. werneriifolia* as distantly related, with *P. werneriifolia* showing to be most closely related to *P. castoreus*, another Central NA subclade (Fig. 6).

The clade directly below the *Packera castoreus* and *P. werneriifolia* clade contains *Packera dimorphophylla* var. *dimorphophylla* and *P. crocata*, two widespread species found in Central NA. Both species are known to hybridize with each other forming a distinctive complex, and are “undoubtedly closely related” (Trock, 1999). Additionally, *P. crocata* and *P. dimorphophylla* have orange-yellow to almost brick colored ray florets compared to the typical bright yellow (Trock, 1999). Thus, it is understandable that these taxa would be closely related with high support in the nuclear tree (1.0LPP; Supplemental Fig. 3), and sister in the plastid tree (Fig. 3 & Supplemental Fig. 4).

The fourth and final Central NA clade contains *P. streptanthifolia* var. *streptanthifolia* and *P. dimorphophylla* var. *paysonii*, as sister taxa with high support (1.0LPP; Supplemental Fig. 3). Interestingly, there are not many similarities between the two taxa other than overlapping distributions (Trock, 1999), so their placement is somewhat surprising, especially because of known interactions that these species have with other taxa. For example, it has previously been hypothesized that there is a close relationship between *P. streptanthifolia* and *P. thurberi* (=*P. tridenticulata*), leading to potential hybridization events resulting in the variety *P. streptanthifolia* var. *borealis* (Barkley, 1962; Bain, 1988), though that is not seen within our phylogeny.

**Species relationships in remaining Senecioneae taxa**

Subtribe Senecioninae

Recent phylogenomic work in tribe Senecioneae has shown that *Bethencourtia* (not tested), *Emilia*, *Erechtites*, *Jacobaea*, and *Pericallis* are the most closely related genera to *Packera* (Funk et al., 2009; Pelser et al., 2007, 2010). Our nuclear phylogeny recovered a clade containing *Pericallis* and *Elekmania* as sister to *Packera* (Supplemental Fig. 5). The relationship between *Packera* and *Pericallis* is expected given that previous phylogenetic work showed strong support that *Packera* was most closely related to *Pericallis* (Panero et al., 1999; Bain & Golden, 2000), along with *Jacobaea vulgaris*, who appears more genetically distant in our nuclear and plastid phylogenies and is sister to *Erechtites* (Supplemental Fig. 5). Bain et al (1997) had originally predicted that *Packera* and *Pericallis* would be related given they both have the Helianthoid pollen type, though that does not hold true with other tested members that also have the same pollen type (i.e., *Doronicum* and *Telanthophora*), since they place distantly to *Packera* (Supplemental Fig. 5). Interestingly, when included, *Elekmania* has a more distant relationship to *Packera* in other phylogenies compared to ours (Pelser et al., 2007).

*Xenophyllum* and *Werneria* are next most closely related to *Packera*, which is unexpected since there is not a known connection between *Xenophyllum* and *Werneria* with either *Packera* or *Pericallis*. For example, their distributions do not appear to overlap: *Xenophyllum* and *Werneria* are found exclusively in South America, differing from *Packera* (North American endemic) and *Pericallis* (Canary Islands, Spain endemic), and they have varying morphologies (Funk et al., 2009). We predict these unusual results may be from low species sampling across Senecioneae in this analysis. However, the relationship seen between *Xenophyllum* and *Werneria* in the nuclear and plastid trees, sister with high support (1.0LPP/100BS, Supplemental Figs. 3 & 4), is expected given that the genus *Xenophyllum* was formed from members previously included in the genus *Werneria* (Funk, 1997).

*Emilia* is sister to the remaining Senecioninae taxa, and ultimately the most distantly related to *Packera*, which is surprising given that previous phylogenetic work has placed *Emilia* as closely related to *Packera* many times (Funk et al., 2009; Pelser et al., 2007, 2010). The plastid phylogeny agrees with the nuclear phylogeny by placing *Emilia* as more distantly related to *Packera*, but alternatively has *Pericallis*, *Xenophyllum*, and *Werneria* within the same smaller clade as *Emilia* instead of being sister to *Packera* (Supplemental Fig. 5). Thus, the plastid relationships are almost inverse to the nuclear relationships.

Subtribe Tussilagininae s.str.

Members of the currently recognized Tussilagininae s.str. subtribe are paraphyletic with *Tephroseris newcombei* placing outside of the main Tussilagininae s.str. subtribe clade and sister to tribe Senecioneae in the nuclear tree (Supplemental Fig. 3). This differs from the plastid tree which placed *T. newcombei* not within subtribe Tussilagininae s.str. (current placement), but instead as a member of *Packera*. Although *T. newcombei* was once placed into *Packera* as *P. newcombei* (Weber & Löve, 1981), there is an overwhelming consensus that it does not belong in *Packera*, or possibly even within Senecioneae (Janovec & Barkley, 1996; Bain & Golden, 2000; Golden et al., 2001; Nordenstam & Pelser, 2011). The genus *Tephroseris* has a complicated history, along with two other genera *Sinosenecio* B. Nord. and *Nemosenecio* (Kitam.) B. Nord., where their taxonomic placement within Senecioneae has changed continuously (Nordenstam & Pelser, 2011). These three genera are sometimes even recognized as a separate subtribe Tephroseridinae (Jeffrey & Chen, 1984). Even though *Sinosenecio* and *Nemosenecio* species were not tested in this study, *Tephroseris* placing outside of Tussilagininae s.str. provides additional evidence of the new subtribe’s validity; however, more research is needed to formally recognize this new subtribal status.

Earlier work in Tussilagininae s.str. has shown a close relationship between *Telanthophora* and *Roldana*, with *Telanthophora* sometimes nesting within *Roldana* (Quedensley et al., 2018), though not always (Pelser et al., 2010). *Telanthophora* and *Roldana* are placing as sister with high support in the nuclear tree (Supplemental Fig. 3), following the previous findings of Pelser et al. (2010), while our plastid tree is showing their separation with *Roldana* sister to *Doronicum* (Supplemental Fig. 4), which is likely not the true relationship.

Previous phylogenetic work in Senecioneae separated Tussilagininae s.str. into multiple subclades, one of which is the *Senecio medley-woodii*–*Brachyglottis* clade comprised of *Brachyglottis*, eight allied genera, and some African succulent species formerly classified as *Senecio*. This clade ultimately places sister to the remaining Tussilaginineae s.str. taxa and Othonninae subtribe (Pelser et al., 2007; Funk et al., 2009). Though no Othonninae or *S. medley-woodii*–*Brachyglottis* clade *Senecio* taxa were included in our phylogeny, our nuclear and plastid phylogenies place *Brachyglottis* within Tussilagininae s.str. but sister to the remaining Tussilagininae s.str. taxa, providing support for previous findings’ placement of *Brachyglottis* within Senecioneae.

Subtribe Doroniceae

Similarly to *Tephroseris*, *Doronicum* also placed outside of the currently recognized Senecioneae subtribes in both our nuclear and plastid phylogenies (Supplemental Fig. 5). *Doronicum* is a genus in Asteraceae that has traditionally been included in the tribe Senecioneae based on gross morphology. However, recent phylogenetic studies have found that there is little support for the inclusion of *Doronicum* in Senecioneae (Goertzen et al., 2003; Panero, 2005; Funk et al., 2009; Pelser et al., 2007, 2010; Ren et al., 2021, Zhang et al., 2021). In fact, ITS data has demonstrated that *Doronicum* is distantly related to Senecioneae and appears closer to the tribes Astereae and Gnaphalieae (Goertzen et al., 2003). Plastid and transcriptome data agrees with ITS that *Doronicum* is not closely related to Senecioneae, but instead shows *Doronicum* closest to tribe Calenduleae (Panero, 2005; Ren et al., 2021). Panero (2005) formally placed *Doronicum* into a monogeneric subtribe, Doroniceae, which is continually gaining support (Fu et al., 2016; Ren et al., 2021; Zhang et al., 2021); however, it’s tribal level affiliation is still uncertain and not fully accepted (Funk et al., 2009).

**References**

Bah M, Bye R, Pereda-Miranda R. 1994. Hepatotoxic pyrrolizidine alkaloids in the Mexican medicinal plant *Packera candidissima* (Asteraceae: Senecioneae). *Journal of Ethnopharmacology* 43: 19–30.

Bain JF. 1988. Taxonomy of *Senecio streptanthifolius* Greene. *Rhodora* 90: 277–312.

Bain JF, Whitton J. 1994. Taxonomìc analysis of *Senecio pauciflorus* and *S. indecorus* (Asteraceae). *Nordic Journal of Botany* 14: 193–199.

Bain JF, Jansen RK. 1995. A phylogenetic analysis of the aureoid *Senecio* (Asteraceae) complex based on ITS sequence data. *Plant Systematics and Evolution* 195(3–4): 209–219.

Bain JF, Tyson BS, Bray DF. 1997. Variation in pollen wall ultrastructure in New World Senecioneae (Asteraceae), with special reference to *Packera*. *Canadian Journal of Botany* 75: 730–735.

Bain JF, Golden JL. 2000. A phylogeny of *Packera* (Senecioneae; Asteraceae) based on internal transcribed spacer region sequence data and a broad sampling of outgroups. *Molecular Phylogenetics and Evolution* 16(3): 331–338.

Bain JF, Golden JL. 2005. Chloroplast haplotype diversity patterns in *Packera pauciflora* (Asteraceae) are affected by geographical isolation, hybridization, and breeding system. *Canadian Journal of Botany* 83(8): 1039–1045.

Barkley TM. 1962. A Revision of *Senecio aureus*. *Transactions of the Kansas Academy of Science (1903-)* 65: 318–364.

Barkley TM. 1978. Three new species of *Senecio* (Asteraceae) from Mexico. *Brittonia* 30: 69–75.

Barkley TM. 1981. *Senecio* and *Erechtites* (Compositae) in the North American Flora: Supplementary Notes. *Brittonia* 33:523–527.

Barkley TM. 1988. Variation among the Aureoid *Senecios* of North America: A geohistorical interpretation. *The Botanical Review* 54(1): 82–106.

Barkley TM. 1989. New taxa and nomenclatural combinations in *Senecio* in Mexico and the United States. *Phytologia.* 67: 237–253.

Chapman GC, Jones SB, Jones Jr. SB. 1971. Hybridization between *Senecio smallii* and *S. tomentosus* (Compositae) on the granitic flatrocks of the Southeastern United States. *Brittonia* 23: 209–216.

Cronquist A. 1955. *Vascular plants of the Pacific Northwest. Part 5; Compositae.* University of Washington Press, Seattle.

Elven R, Murray D, Razzhivin V, Yurtsev BA. 2011.Annotated Checklist of the Panarctic Flora (PAF) Vascular plants version 1.0. *Available at* *http://panarcticflora.org/results?biogeographic=&bioclimatic=&region=&name=packera+heterophylla#paf-863101* (accessed May 17, 2022).

Fernald ML. 1943. Virginia botanizing under restrictions. *Rhodora* 45:485–511.

Fernald ML. 1950. *Gray’s Manual of Botany*. New York: American Book Co.

Fragoso-Serrano M, Figueroa-González G, Castro-Carranza E, Hernández-Solis F, Linares E, Bye R, Pereda-Miranda R. 2012. Profiling of alkaloids and eremophilanes in miracle tea (*Packera candidissima* and *P. bellidifolia*) products. *Journal of Natural Products* 75: 890–895.

Freeman CC. 1985. A revision of the aureoid species of *Senecio* (Asteraceae: Senecioneae) in Mexico, with a cytogeographic and phylogenetic interpretation of the aureoid complex. Ph.D. Dissertation. Kansas State University.

Freeman CC, Barkley TM. (1995). A synopsis of the genus Packera (Asteraceae: Senecioneae) in Mexico. *SIDA, Contributions to Botany*, *16*(4), 699–709.

Fu ZX, Jiao BH, Nie B, Zhang GJ, Gao TG, Chen ZD, Lu AM, Kong HZ, Wang XQ, Wang YZ, Zhou SL, Zhang SZ, Wang XM, Liu ZJ, Wang QF, Li JH, Li DZ, Yi TS, Hong MA, Soltis DE, Soltis PS, Li JH, Fu CX, Liu QX. 2016. A comprehensive generic-level phylogeny of the sunflower family: Implications for the systematics of Chinese Asteraceae. *Journal of Systematics and Evolution* 54: 416–437.

Funk VA. 1997. *Xenophyllum*, a New Andean Genus Extracted from *Werneria*. *Novon* 7: 235–241.

Funk VA, Susanna A, Stuessy TF, Bayer RJ (eds.). 2009. *Systematics, Evolution, and Biogeography of Compositae*. Vienna: International Association for Plant Taxonomy.

Gleason HA, Cronquist A. 1991. *Manual of Vascular Plants of Northeastern United States and Adjacent Canada*. Bronx: The New York Botanic Garden.

Goertzen LR, Cannone JJ, Gutell RR, Jansen RK. 2003. ITS secondary structure derived from comparative analysis: Implications for sequence alignment and phylogeny of the Asteraceae. *Molecular Phylogenetics and Evolution* 29: 216–234.

Golden JL, Kim YD, Bain JF. 2001. A re-evaluation of North American *Tephroseris* and *Sinosenecio* (Asteraceae: Senecioneae) based on molecular and micromorphological data. *Canadian Journal of Botany* 79: 1195–1201.

Gramling A. 2006. A conservation assessment of *Packera millefolium*, a Southern Appalachian Endemic. Ph.D. Dissertation. University of North Caroline at Chapel Hill.

Janovec JP, Barkley TM. 1996. *Sinosenecio newcombei* (Asteraceae : Senecioneae): A New Combination for a North American Plant in an Asiatic Genus. *Novon* 6: 265–267.

Jeffrey C, Chen Y-L. 1984. Taxonomic Studies on the Tribe Senecioneae (Compositae) of Eastern Asia. *Kew Bulletin* 39: 205–446.

Jeffrey C. 1992. The Tribe Senecioneae (Compositae) in the Mascarene Islands with an Annotated World Check-List of the Genera of the Tribe: Notes on Compositae: VI. *Kew Bulletin* 47: 49.

Kowal RR, Mahoney AM, Weakley AS, Estes D. 2015. Validation and lectotypification of *Packera crawfordii* (Asteraceae). *Phytoneuron* 2015: 1–2.

Kowal RR, Mahoney AM. 2016. Comments on the status of *Packera crawfordii* (Asteraceae, Senecioneae), a neglected species of the southeastern United States. *Brittonia* 68: 74–82.

Krasnoborov IM (ed.). 2007. *Flora of Sibera*. New Hampshire, US: An imprint of Edenbridge Ltd., British Isles.

Mahoney AM, Kowal RR. 2008. Three new varieties of *Packera paupercula* (Asteraceae, Senecioneae) in midwestern and southeastern North America. *Novon* 18(2): 220–228.

Nordenstam B, Pelser PB. 2011. Notes on the generic limits of *Sinosenecio* and *Tephroseris* (Compositae - Senecioneae). *Comp. Newsl.* 49: 1–7.

Packer JG. 1972. A taxonomic and phytogeographical review of some arctic and alpine *Senecio* species. *Canadian Journal of Botany* 50: 507–518.

Panero JL, Francisco-Ortega J, Jansen RK, Santos-Guerra A. 1999. Molecular evidence for multiple origins of woodiness and a New World biogeographic connection of the Macaronesian island endemic *Pericallis* (Asteraceae: Senecioneae). *Proceedings of the National Academy of Sciences of the United States of America* 96: 13886–13891.

Panero JL. 2005. New combinations and infrafamilial taxa in the Asteraceae. *Phytologia* 87: 1–14.

Pelser PB, Nordenstam B, Kadereit JW, Watson LE. 2007. An ITS Phylogeny of Tribe Senecioneae (Asteraceae) and a New Delimitation of *Senecio* L. *Taxon* 56: 1077–1104.

Pelser PB, Kennedy AH, Tepe EJ, Shidler JB, Nordenstam B, Kadereit JW, Watson LE. 2010. Patterns and causes of incongruence between plastid and nuclear Senecioneae (Asteraceae) phylogenies. *American Journal of Botany* 97: 856–873.

Quedensley TS, Gruenstaeudl M, Jansen RK. 2018. Phylogenetic relationships of the Mexican tussilaginoid genera (Asteraceae: Senecioneae). *Journal of the Botanical Research Institute of Texas* 12: 481–498.

Radford AE, Ahles HE, Bell CR. 1968. *Manual of the Vascular Flora of the Carolinas*. Chapel Hill: University of North Carolina Press.

Ren C, Wang L, Nie ZL, Johnson G, Yang QE, Wen J. 2021. Development and phylogenetic utilities of a new set of single-/low-copy nuclear genes in Senecioneae (Asteraceae), with new insights into the tribal position and the relationships within subtribe Tussilagininae. *Molecular Phylogenetics and Evolution* 162.

Schilling EE, Floden A. 2015. Barcoding the Asteraceae of Tennessee, tribe Cichorieae. *Phytoneuron* 19:1–8.

Trock DK. 1999. *A revisionary synthesis of the genus* Packera *(Asteraceae: Senecioneae)*. Ph.D. Dissertation. Kansas State University.

Trock DK. 2006. *Packera* Á. Löve & D. Löve. In *Flora of North America Editorial Committee, eds. 1993+* (Volume 20, pp. 570–602).

Uttal LJ. 1982. Promiscuity of *Senecio plattensis* Nutt. in a Virginia County. *Castanea* 47: 344–346.

Uttal LJ. 1984. *Senecio millefolium* T. & G. (Asteraceae) and its introgressants. *SIDA, Contributions to Botany* 10: 216–222.

Vivar AR De, Arciniegas A, Villaseñor JL. 2007. Secondary Metabolites from Mexican Species of the Tribe Senecioneae (Asteraceae). *Journal of the Mexican Chemical Society* 51: 160–172.

Weakley AS, LeBlond RJ, Sorrie BA, Witsell CT, Estes LD, Gandhi K, Mathews KG, Ebihara A. 2011. New combinations, rank changes, and nomenclatural and taxonomic comments in the vascular flora of The Southeastern United States. *Journal of the Botanical Research Institute of Texas* 5: 437–455.

Weber WA, Löve Á. 1981. New combinations in the genus *Packera* (Asteraceae). *Phytologia* 49: 44–50.

Whitton J, Bain JF. 1992. An analysis of morphological variation in *Senecio cymbalaria* (Asteraceae). *Canadian Journal of Botany* 70: 285–290.

Yeatts L, Schneider A, Schneider B. 2011. *Packera mancosana* (Asteraceae: Senecioneae), a new species and shale barren endemic of southwestern Colorado. *Phytoneuron* 1–8.

Zhang C, Huang CH, Liu M, Hu Y, Panero JL, Luebert F, Gao T, Ma H. 2021. Phylotranscriptomic insights into Asteraceae diversity, polyploidy, and morphological innovation. *Journal of Integrative Plant Biology* 63: 1273–1293.
