## Supplementary material for "Resolving evolutionary relationships in the groundsels: phylogenomics, divergence time estimates, and biogeography of *Packera* (Asteraceae: Senecioneae)": Moore-Pollard_Mandel_SupplementalTables-Figures.pdf

Supplemental Table of Contents


| **Name** | **Brief description** | **Page #(s)** |
|---|---|---|
| Supplemental Materials | Species relationships within *Packera* and related Senecioneae taxa | Attached (word document) |
| nuc_alignments.phy | Nuclear supermatrix file | Attached |
| chl_alignments.phy | Plastid supermatrix file | Attached |
| Supplemental Table 1 | Voucher specimens | Attached (excel sheet) |
| Supplemental Table 2 | Raw sequence statistics using *phyx* for each of the samples used in this study | Attached (excel sheet) |
| Supplemental Table 3 | Matrix stats obtained by *AMAS* and stats about the number of loci and loci choice of the nuclear and plastid phylogenies | Attached (excel sheet) |
| Supplemental Table 4 | Presence/Absence values of each species given the ten geographic regions outlined by Takhtajan (1986) | Attached (excel sheet) |
| Supplemental Fig. 1 | Nuclear phylogeny produced using a concatenation approach | 2 |
| Supplemental Fig. 2 | Plastid phylogeny produced using a pseudocoalescent (ASTRAL) approach | 3 |
| Supplemental Fig. 3 | Nuclear phylogeny produced using ASTRAL with all LPP values | 4 |
| Supplemental Fig. 4 | Plastid phylogeny produced using concatenation with all BS values | 5 |
| Supplemental Fig. 5 | Tanglegram comparing taxon placement between nuclear and plastid genome sequences | 6 |
| Supplemental Figs. 6 - 13 | Divergence rate estimates of Scenarios 1-8 using treePL | 7 - 14 |
| Supplemental Figs. 14 - 21 | Divergence rate estimates of Scenarios 1-8 using RelTime | 15 - 22 |
| Supplemental Figs. 22 & 23 | BioGeoBEARS results using DEC | 23 - 24 |
| Supplemental Figs. 24 & 25 | BioGeoBEARS results using DEC+*J* | 25 - 26 |
| Supplemental Figs. 26 & 27 | BioGeoBEARS results using DIVALIKE | 27 - 28 |
| Supplemental Figs. 28 & 29 | BioGeoBEARS results using DIVALIKE+*J* | 29 - 30 |
| Supplemental Figs. 30 & 31 | BioGeoBEARS results using BAYAREALIKE | 31 - 32 |
| Supplemental Figs. 32 & 33 | BioGeoBEARS results using BAYAREALIKE+*J* | 33 - 34 |

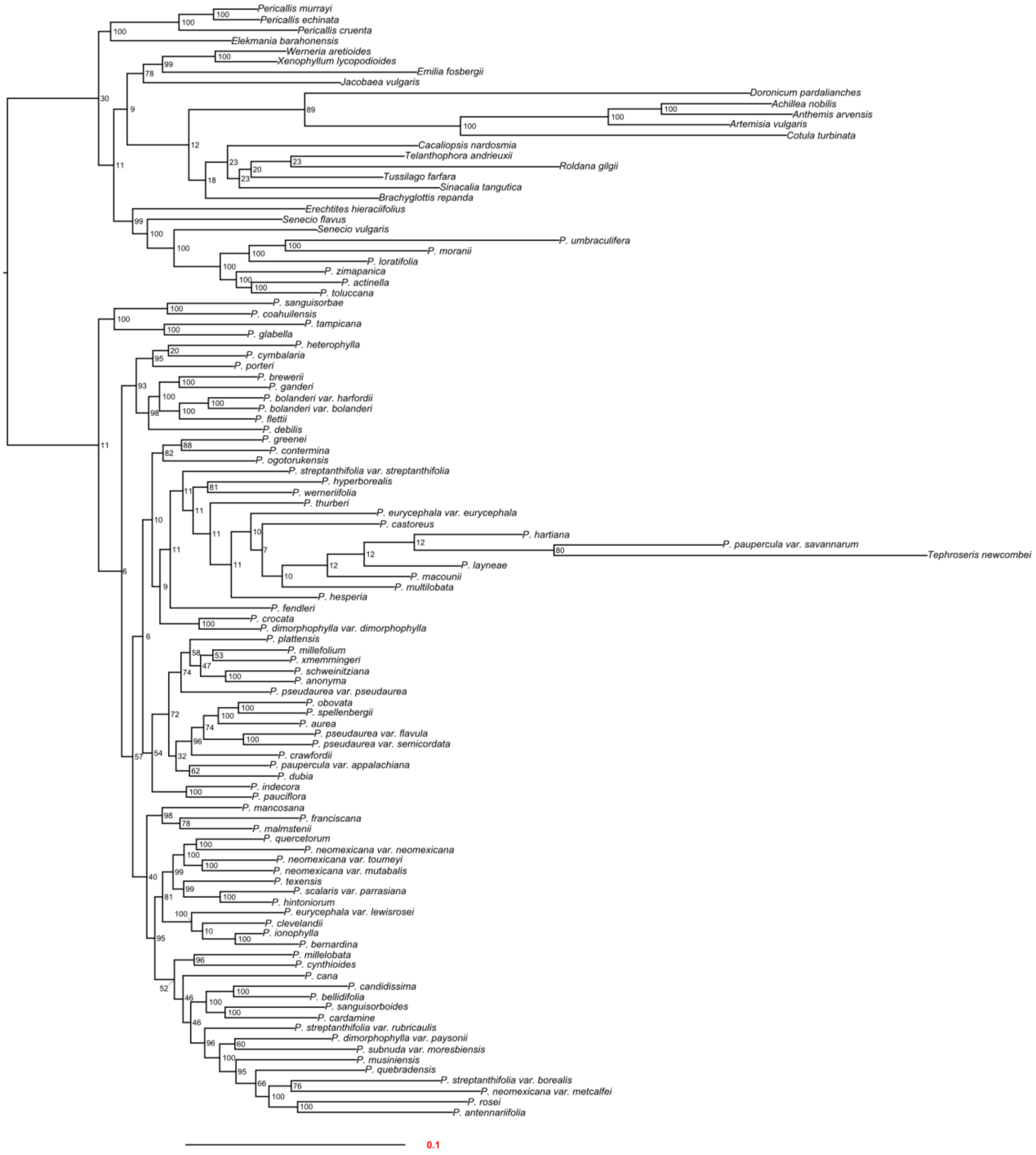


**Supplemental Fig. 1.** Nuclear phylogeny produced using a concatenation approach and inferred with maximum likelihood (RAxML). Scale bar under the tree on the left represents the mean number of nucleotide substitutions per site. Support values (bootstraps) are indicated next to the node. Phylogeny generated using *phytools* in R.

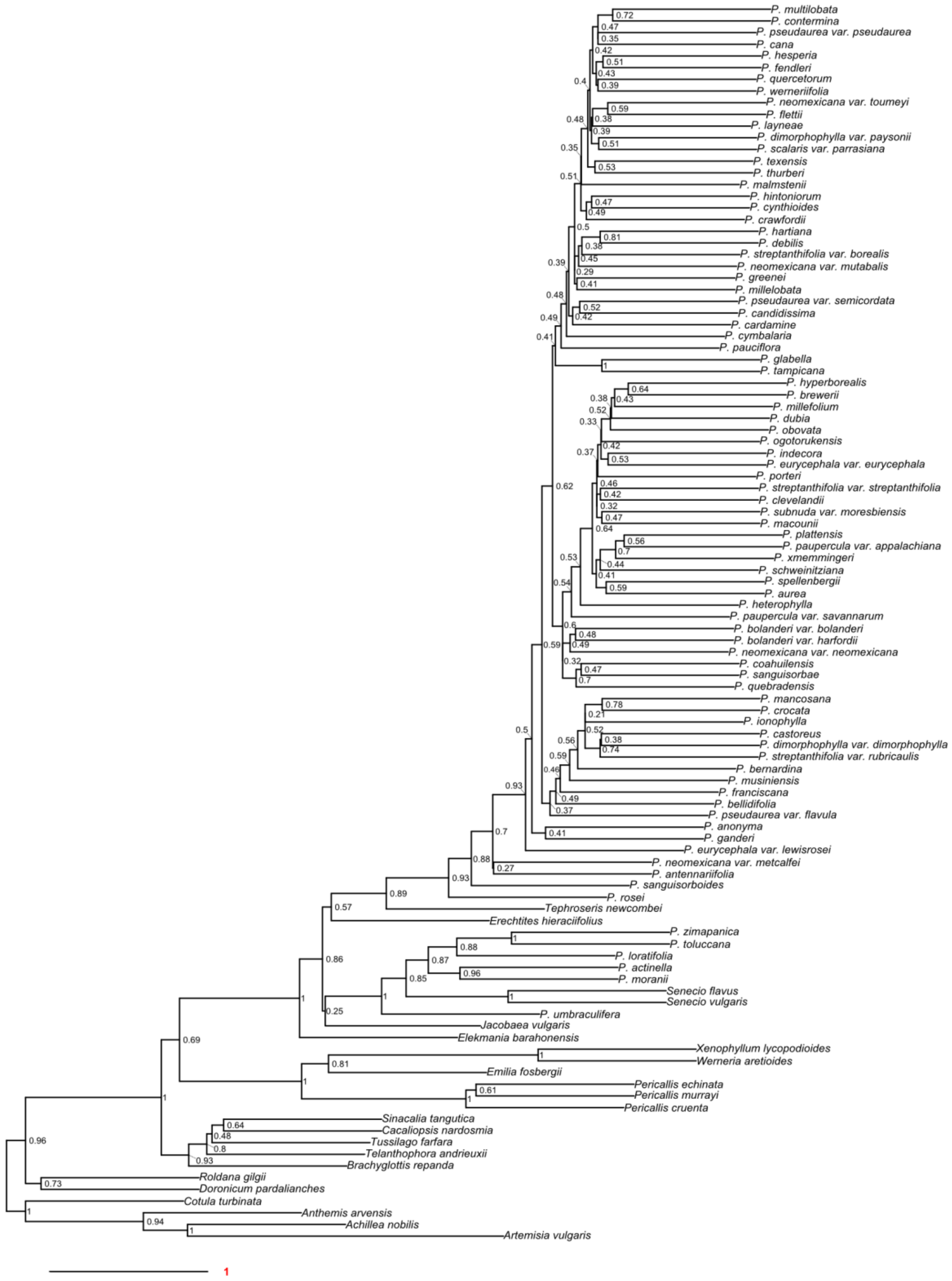


**Supplemental Fig. 2.** Plastome phylogeny produced using a pseudocoalescent analysis (ASTRAL-III) and inferred with maximum likelihood (RAxML). ASTRAL-III reported that the plastid tree has a normalized quartet score of 0.526, meaning that 53% of the gene trees match the final species tree. Scale bar under the tree on the left represents the mean number of nucleotide substitutions per site. Support values (LPP) are next to the node. Phylogeny generated using *phytools* in R.

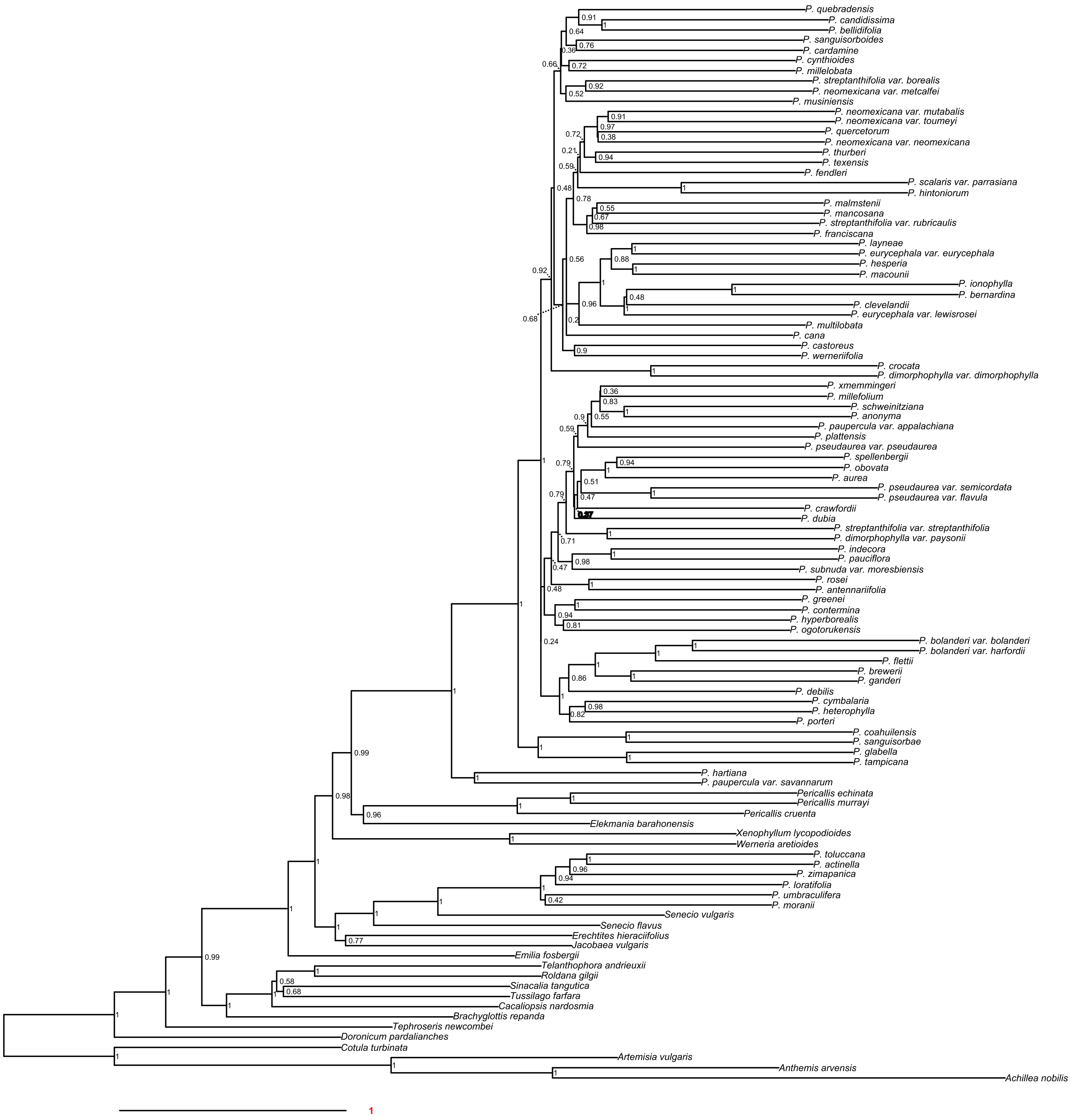


**Supplemental Fig. 3.** Figure 2 nuclear phylogeny produced using a pseudocoalescent analysis (ASTRAL-III) and inferred with maximum likelihood (RAxML), with all support values (LPP) indicated next to node. Scale bar on bottom left represents the mean number of nucleotide substitutions per site. Phylogeny generated using *phytools* in R.

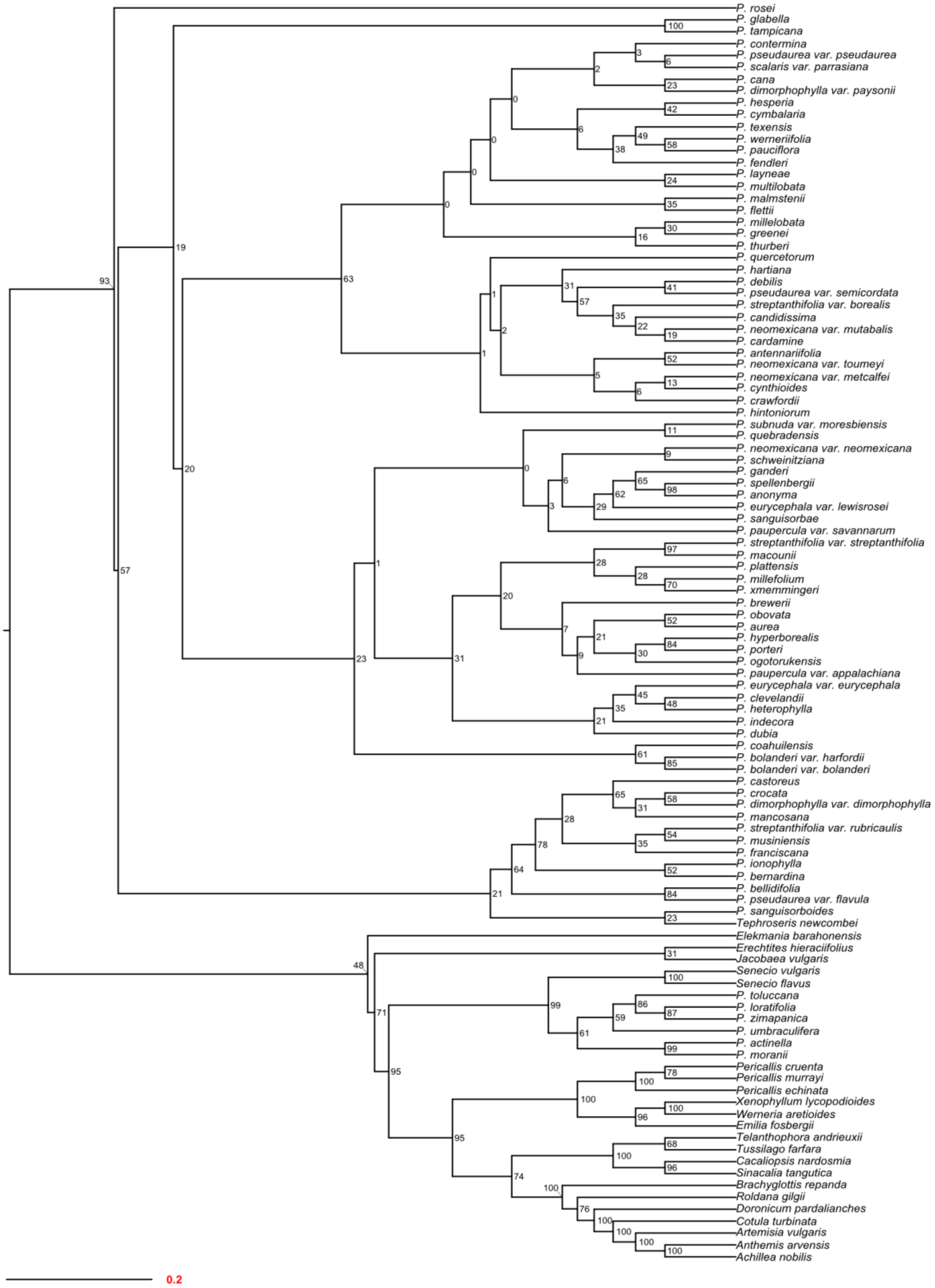


**Supplemental Fig. 4.** Figure 3 plastid phylogeny produced using a concatenated approach, inferred with maximum likelihood (RAxML), with all support values (bootstrap) indicated next to node. Branch lengths were transformed so that node values can be more easily seen. Scale bar on bottom left represents the mean number of nucleotide substitutions per site. Phylogeny generated using *phytools* in R.

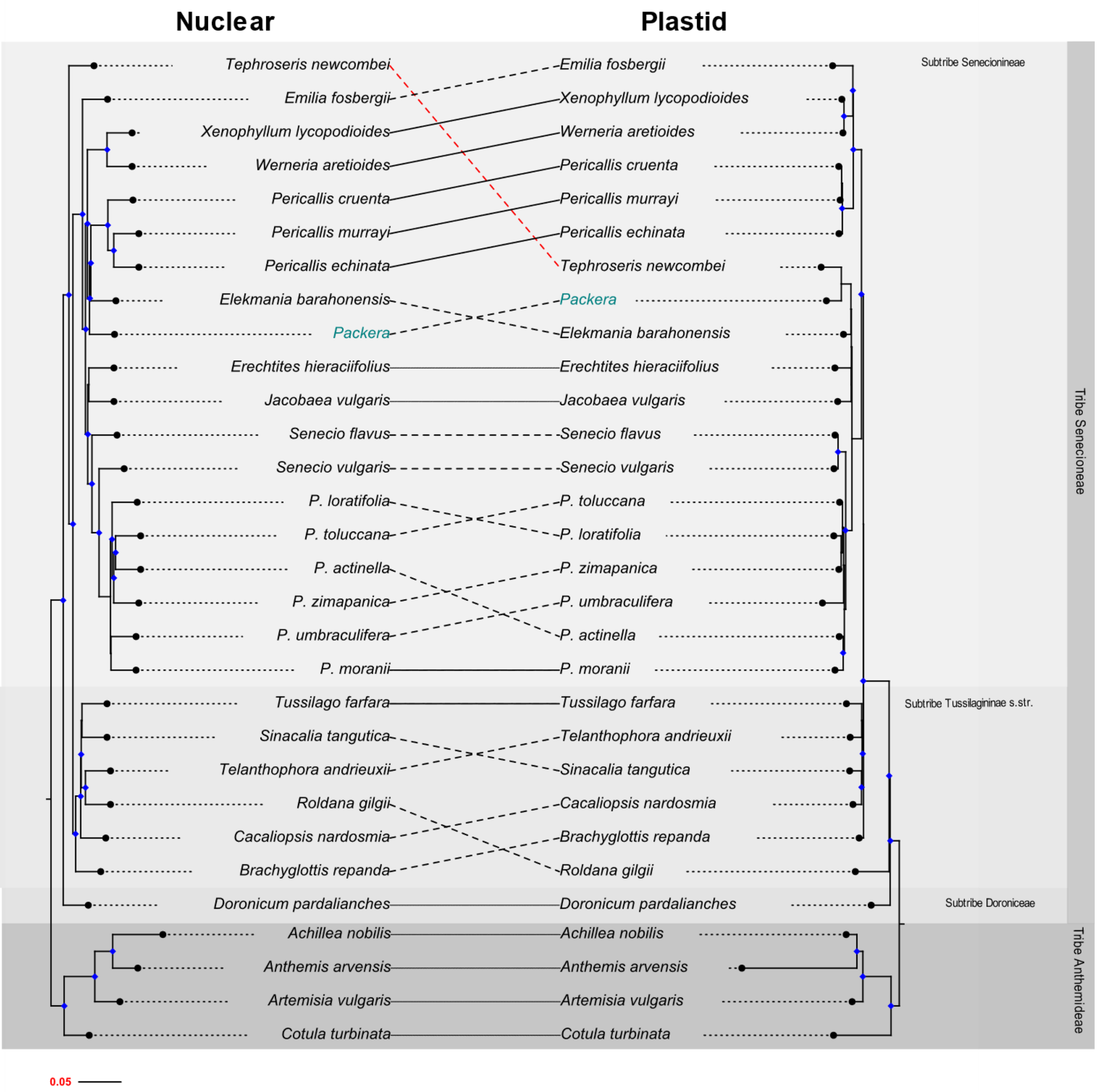


**Supplemental Fig. 5.** Tanglegram comparing taxon placement between nuclear (left) and plastid (right) genome sequences. The nuclear tree was produced using a pseudo-coalescent analysis (ASTRAL-III), while the plastid tree was produced using a concatenated matrix. Both nuclear and plastid trees were then visualized and inferred with maximum likelihood (RAxML) approaches and visualized in R. Support values (LPP [nuclear] and BS [plastid]) of 0.9LPP/90BS or greater are indicated with a blue diamond at the node. Dashed and solid lines represent correspondence between taxa in nuclear and plastid phylogenies: **solid line** – nuclear and plastome relationships agree, **dashed line** – nodes conflict, **red-colored dashed line** – major node conflict. Core *Packera* species' tips are collapsed and indicated by a teal font.

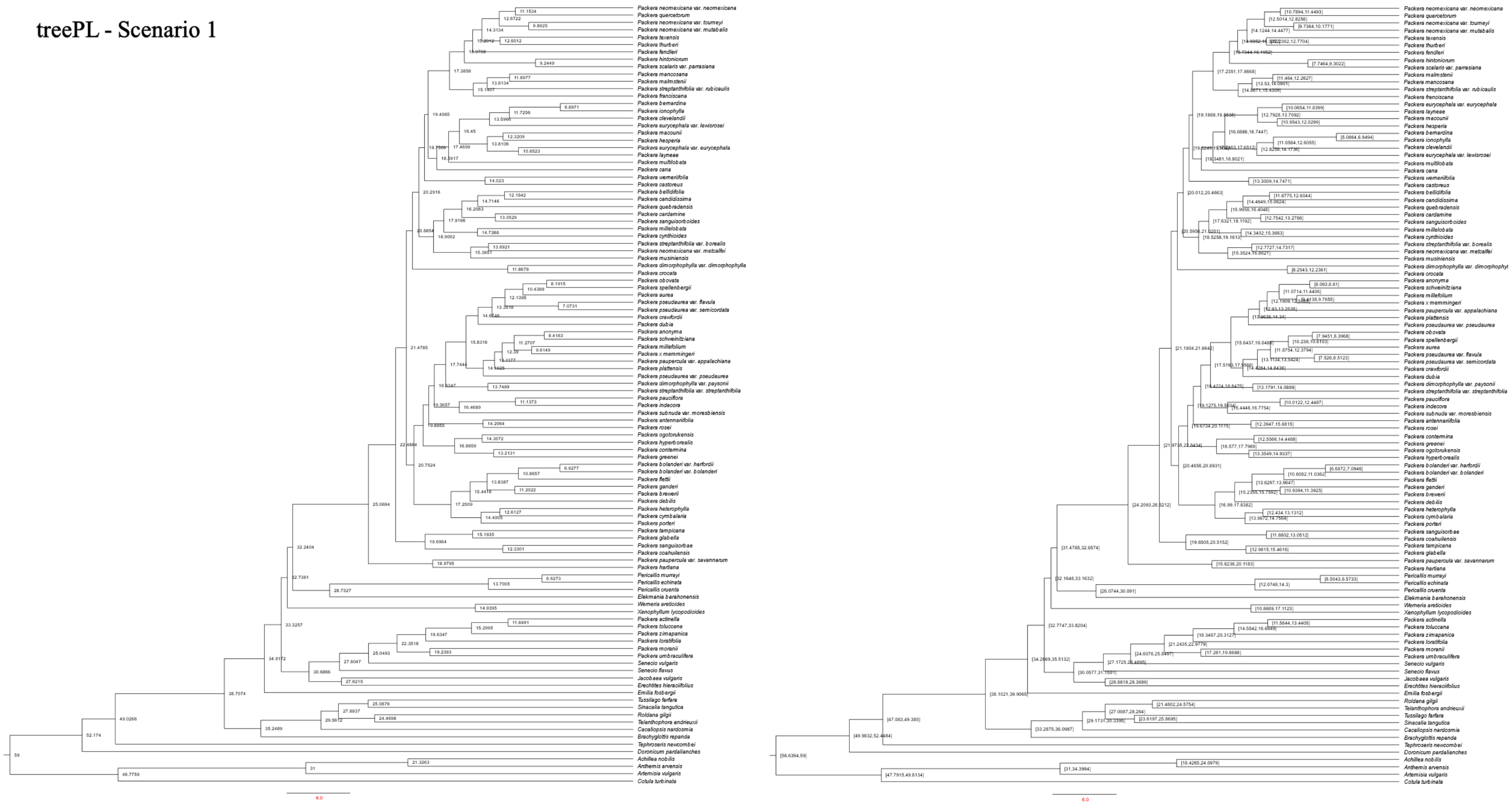


**Supplemental Fig. 6.** Divergence rate estimates of Scenario 1 using treePL in millions of years. Left tree contains the mean age of that designated node, the right tree shows the 95% age range for the designated node.

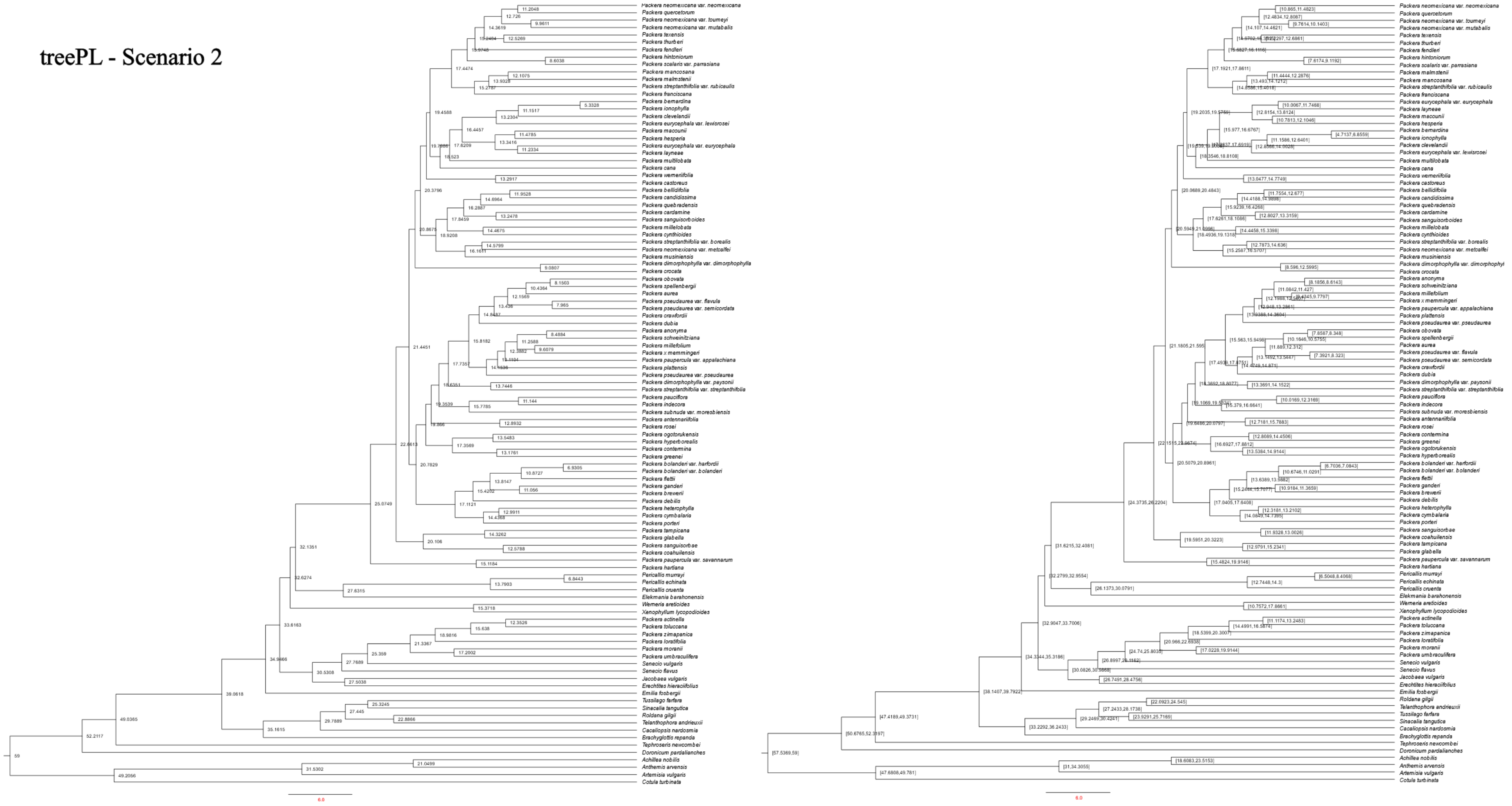


**Supplemental Fig. 7.** Divergence rate estimates of Scenario 2 using treePL in millions of years. Left tree contains the mean age of that designated node, the right tree shows the 95% age range for the designated node.

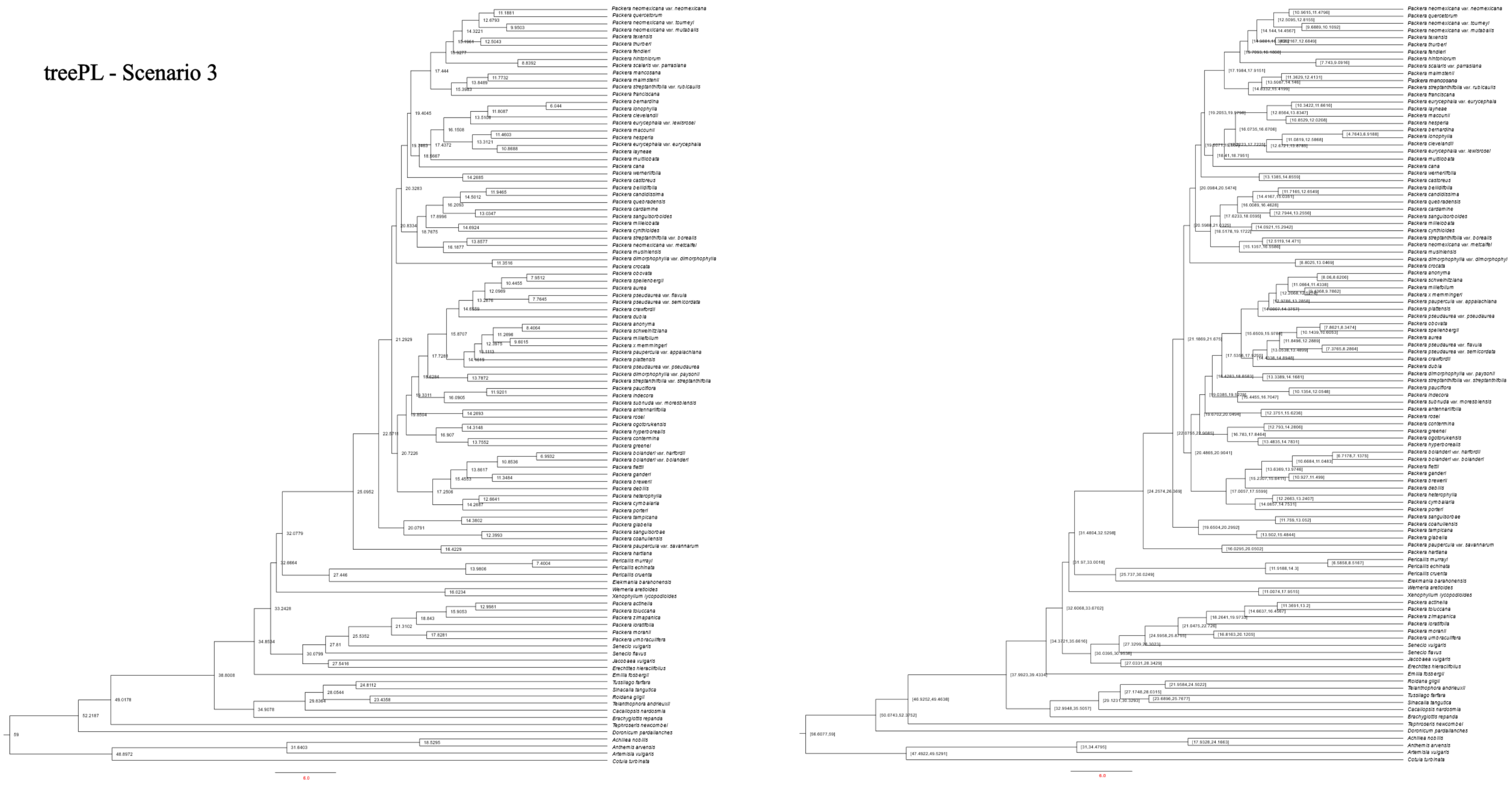


**Supplemental Fig. 8.** Divergence rate estimates of Scenario 3 using treePL in millions of years. Left tree contains the mean age of that designated node, the right tree shows the 95% age range for the designated node.

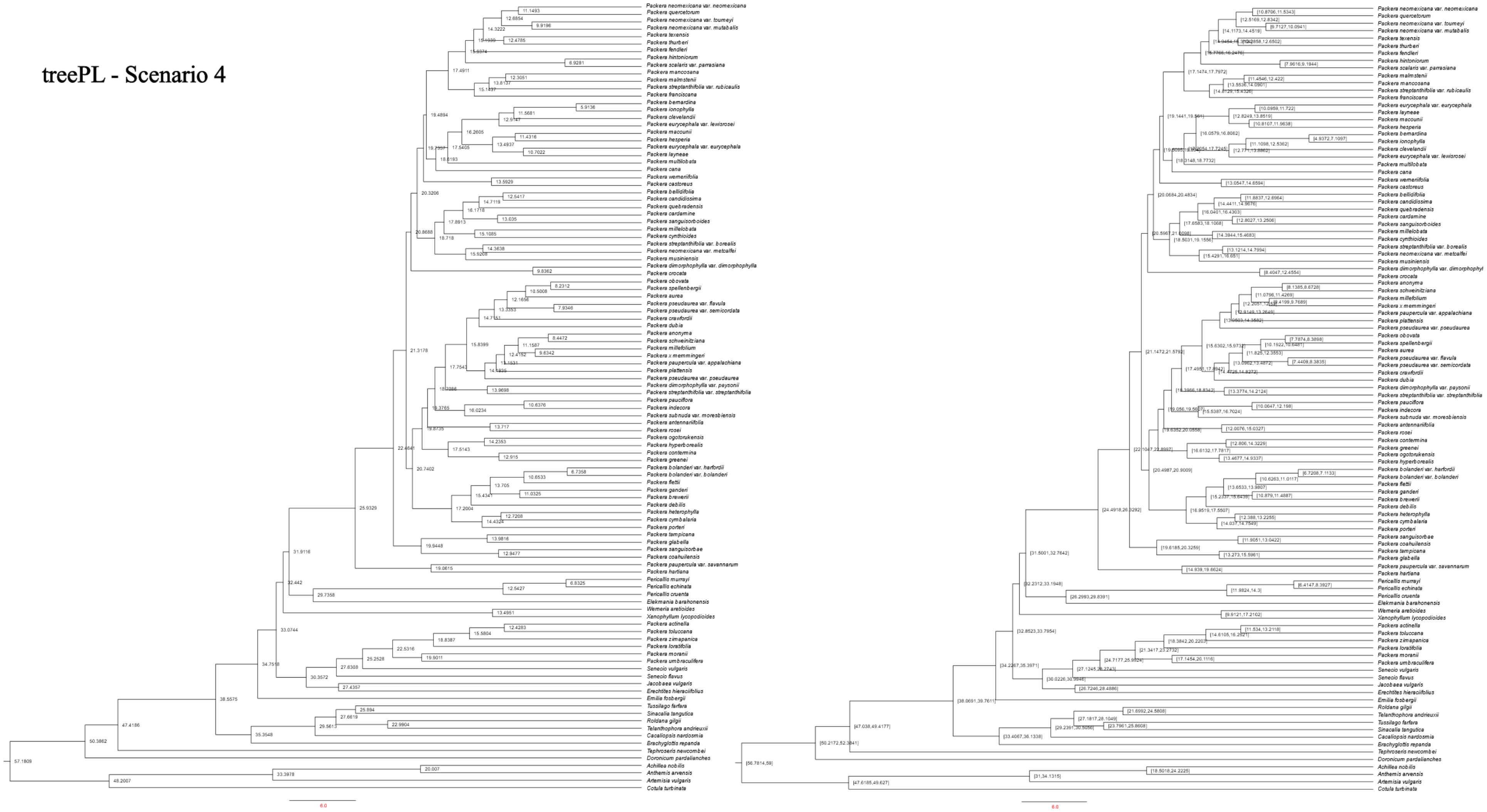


**Supplemental Fig. 9.** Divergence rate estimates of Scenario 4 using treePL in millions of years. Left tree contains the mean age of that designated node, the right tree shows the 95% age range for the designated node.

treePL - Scenario 5

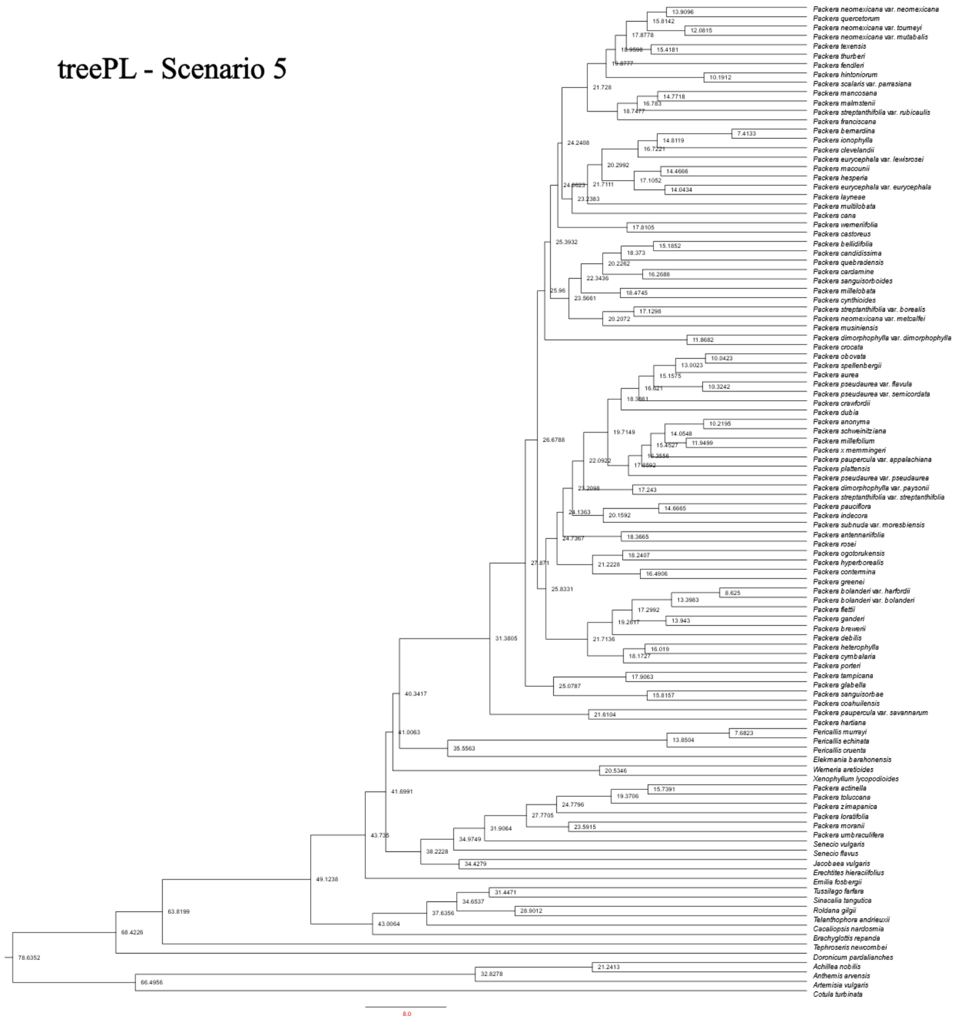

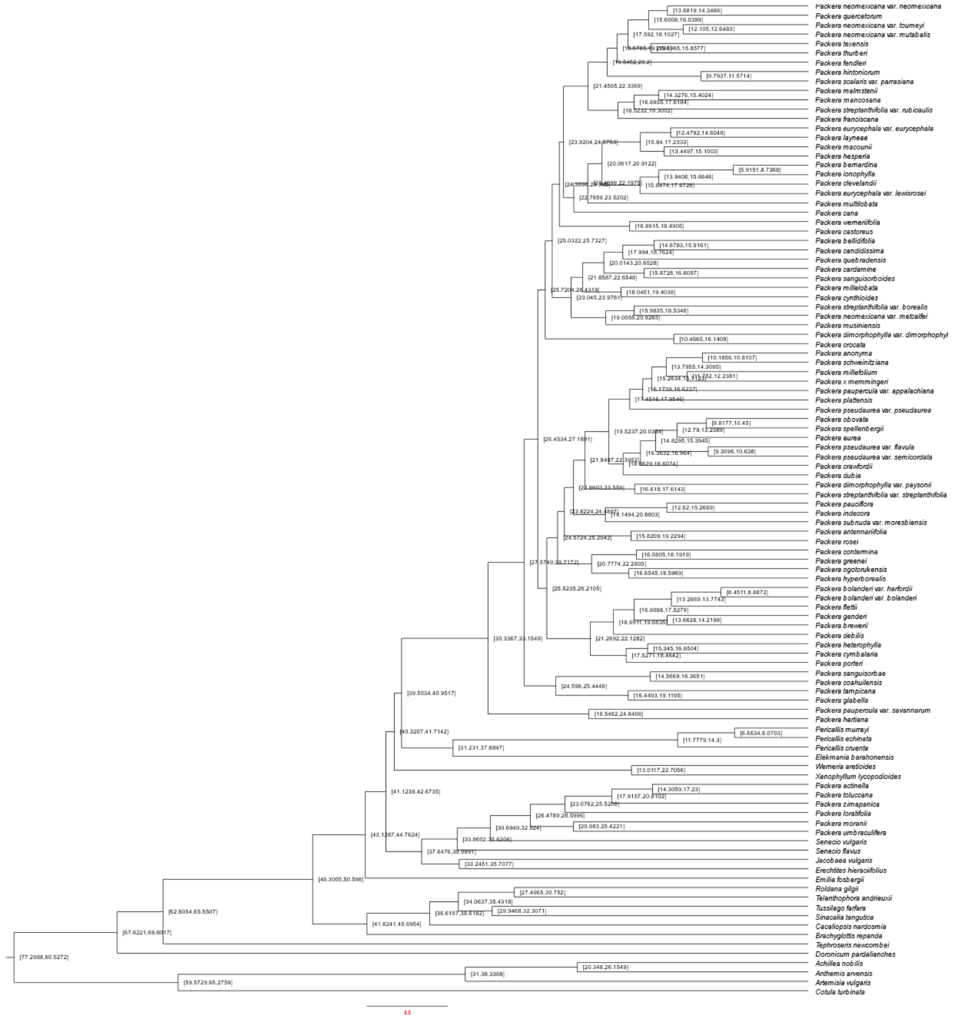

**Supplemental Fig. 10.** Divergence rate estimates of Scenario 5 using treePL in millions of years. Left tree contains the mean age of that designated node, the right tree shows the 95% age range for the designated node.

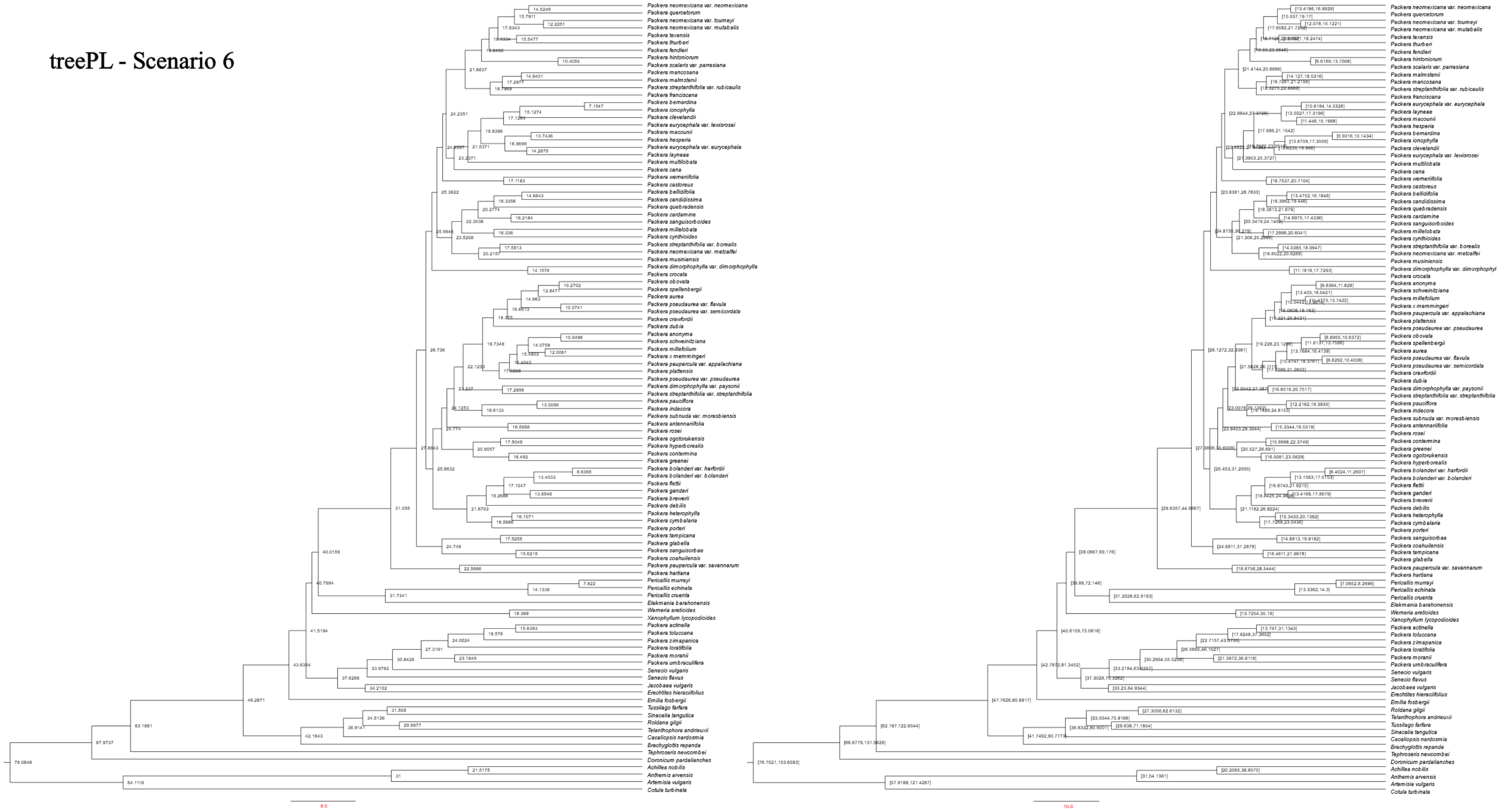


**Supplemental Fig. 11.** Divergence rate estimates of Scenario 6 using treePL in millions of years. Left tree contains the mean age of that designated node, the right tree shows the 95% age range for the designated node.

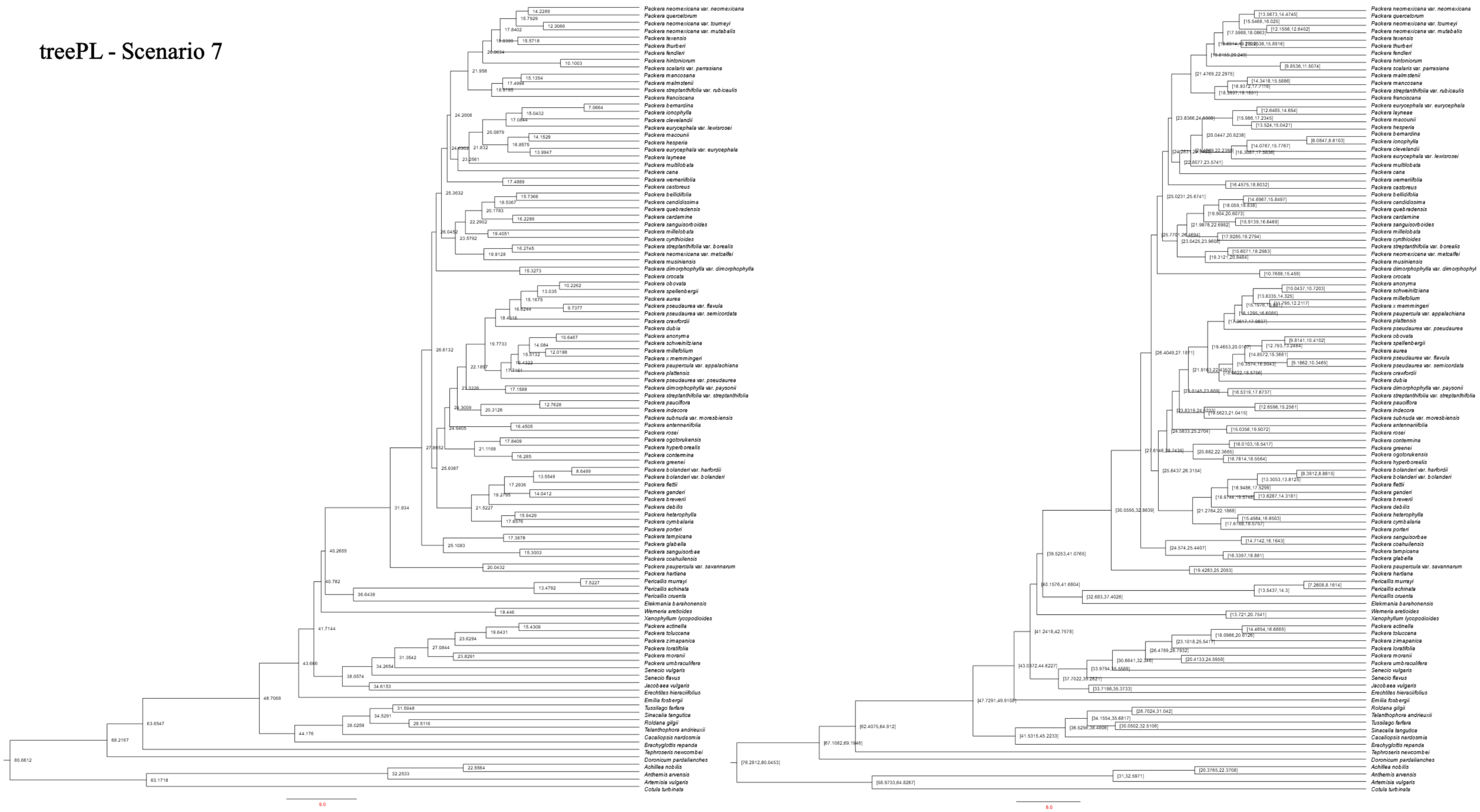


**Supplemental Fig. 12.** Divergence rate estimates of Scenario 7 using treePL in millions of years. Left tree contains the mean age of that designated node, the right tree shows the 95% age range for the designated node.

#### treePL - Scenario 8

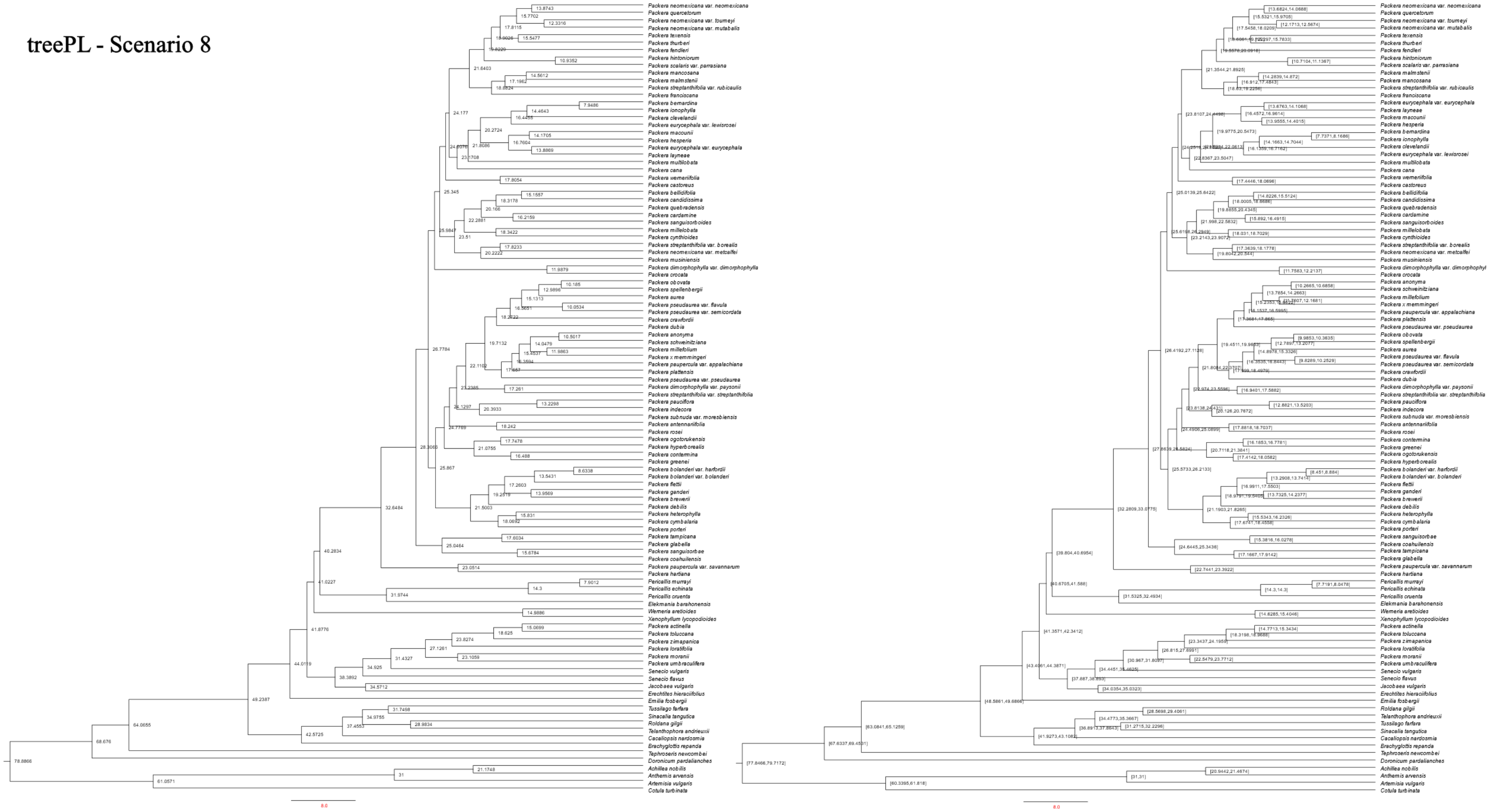

**Supplemental Fig. 13.** Divergence rate estimates of Scenario 8 using treePL in millions of years. Left tree contains the mean age of that designated node, the right tree shows the 95% age range for the designated node.

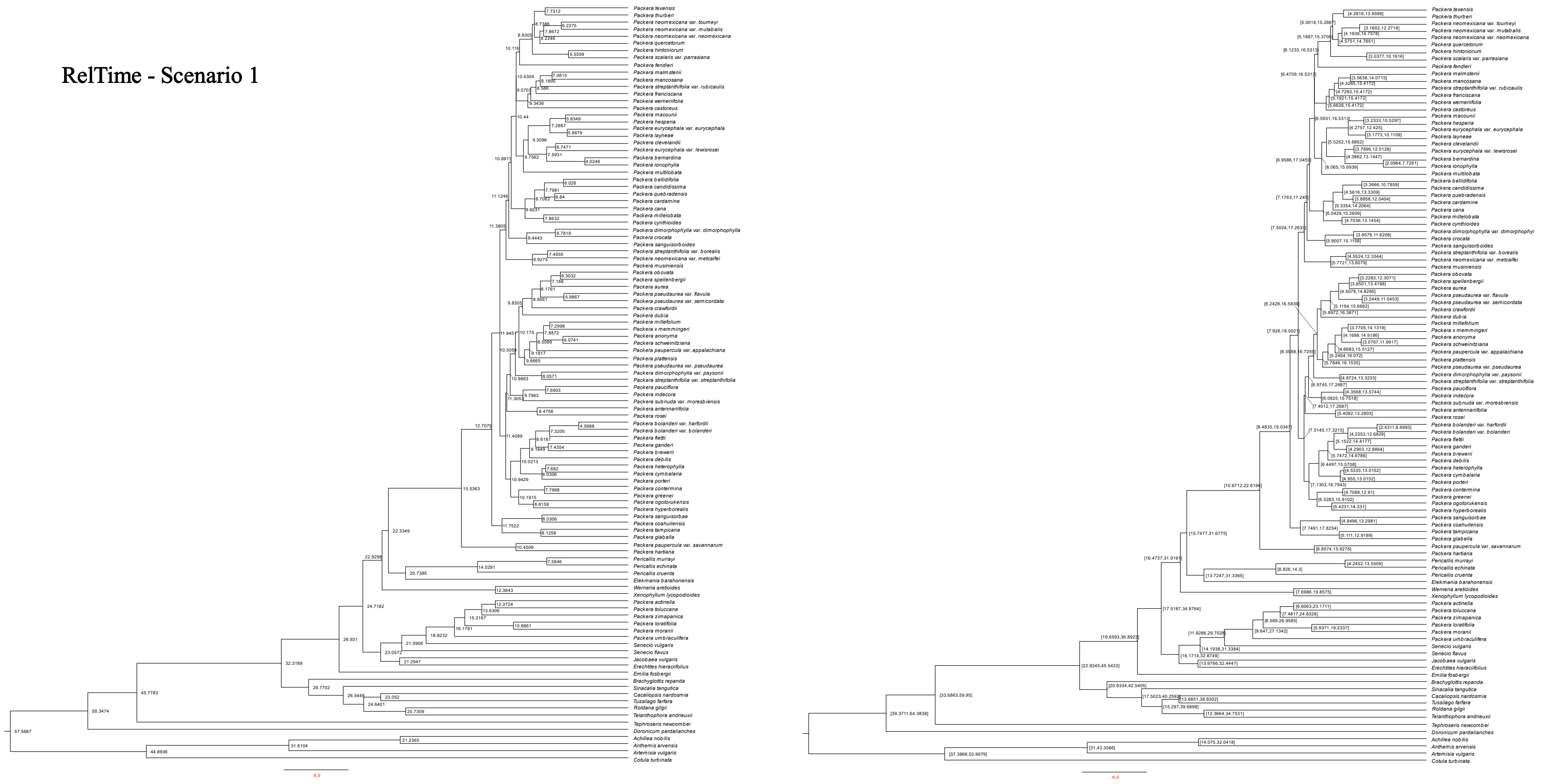


**Supplemental Fig. 14.** Divergence rate estimates of Scenario 1 using RelTime in millions of years. Left tree contains the mean age of that designated node, right tree shows the 95% age range for the designated node.

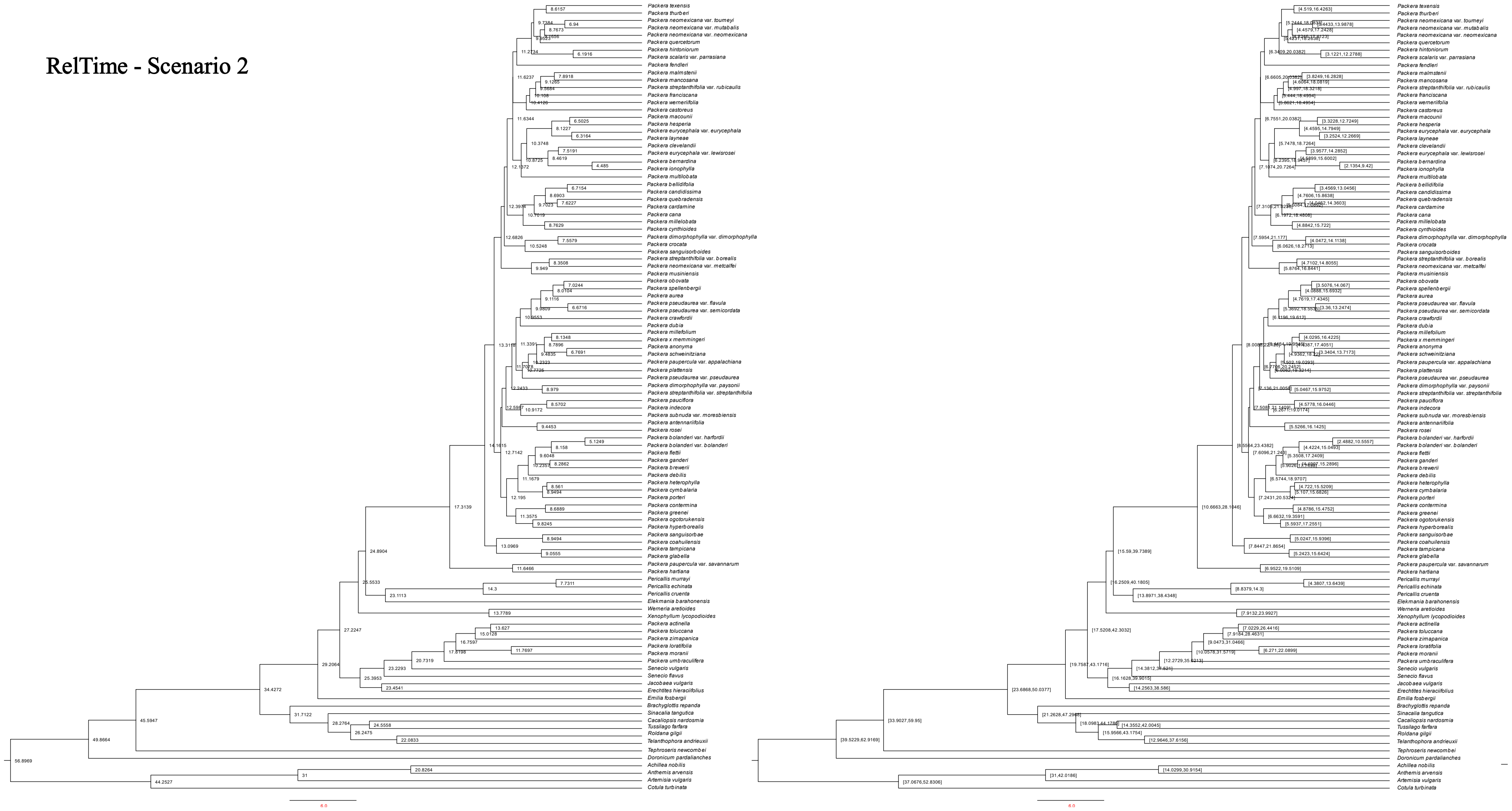


**Supplemental Fig. 15.** Divergence rate estimates of Scenario 2 using RelTime in millions of years. Left tree contains the mean age of that designated node, right tree shows the 95% age range for the designated node.

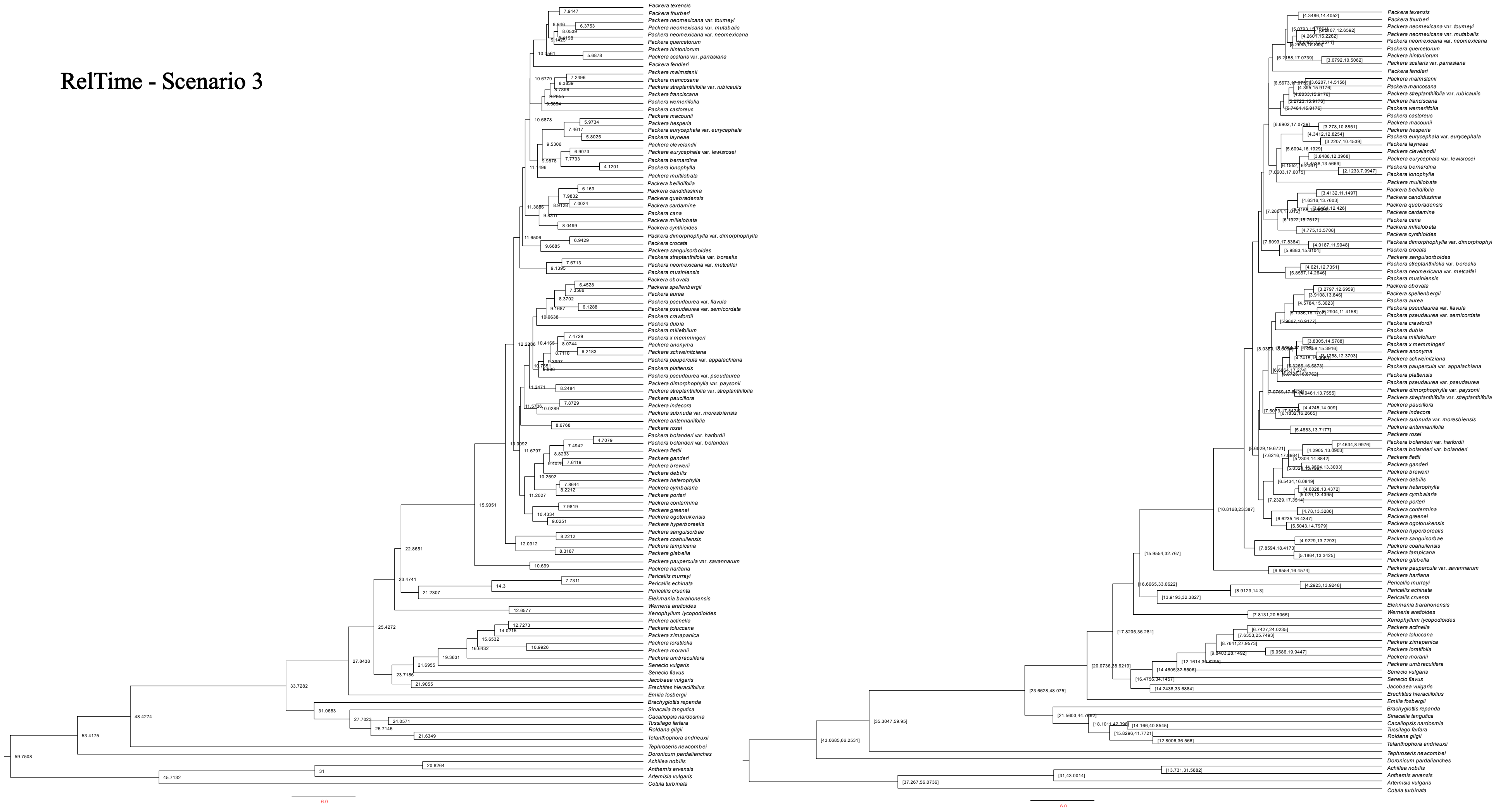


**Supplemental Fig. 16.** Divergence rate estimates of Scenario 3 using RelTime in millions of years. Left tree contains the mean age of that designated node, right tree shows the 95% age range for the designated node.

RelTime - Scenario 4

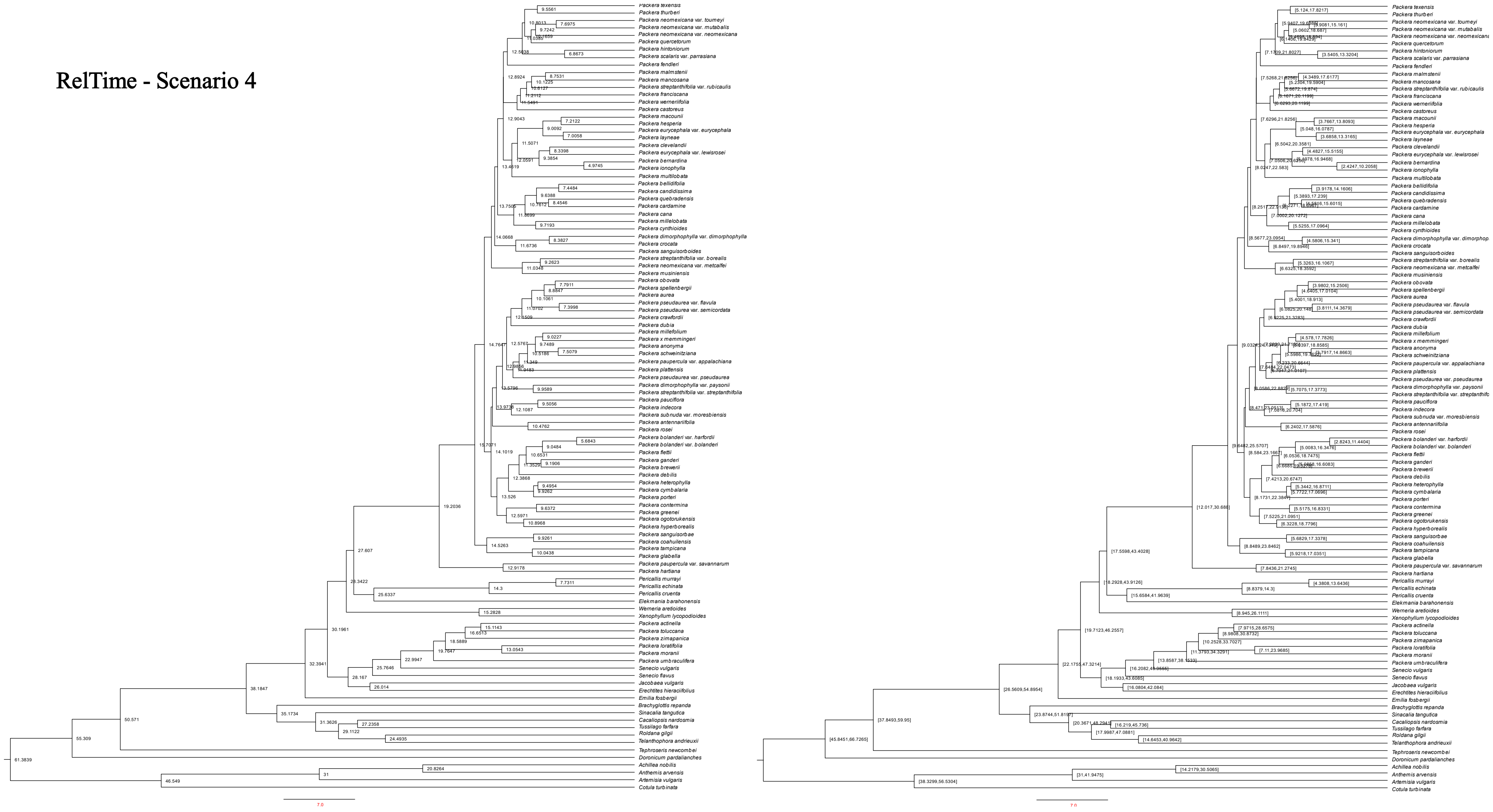

**Supplemental Fig. 17.** Divergence rate estimates of Scenario 4 using RelTime in millions of years. Left tree contains the mean age of that designated node, right tree shows the 95% age range for the designated node.

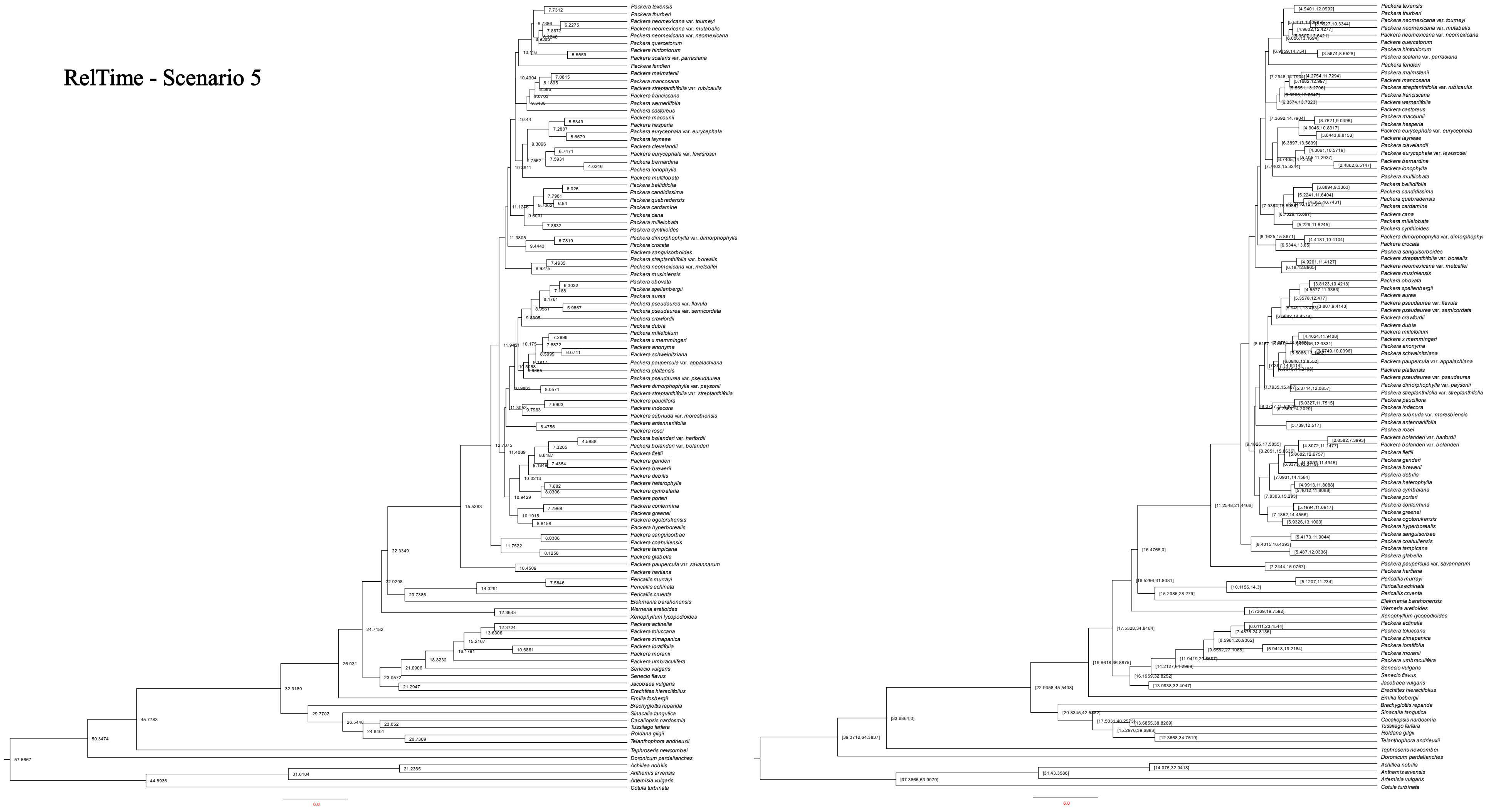


**Supplemental Fig. 18.** Divergence rate estimates of Scenario 5 using RelTime in millions of years. Left tree contains the mean age of that designated node, right tree shows the 95% age range for the designated node.

RelTime - Scenario 6

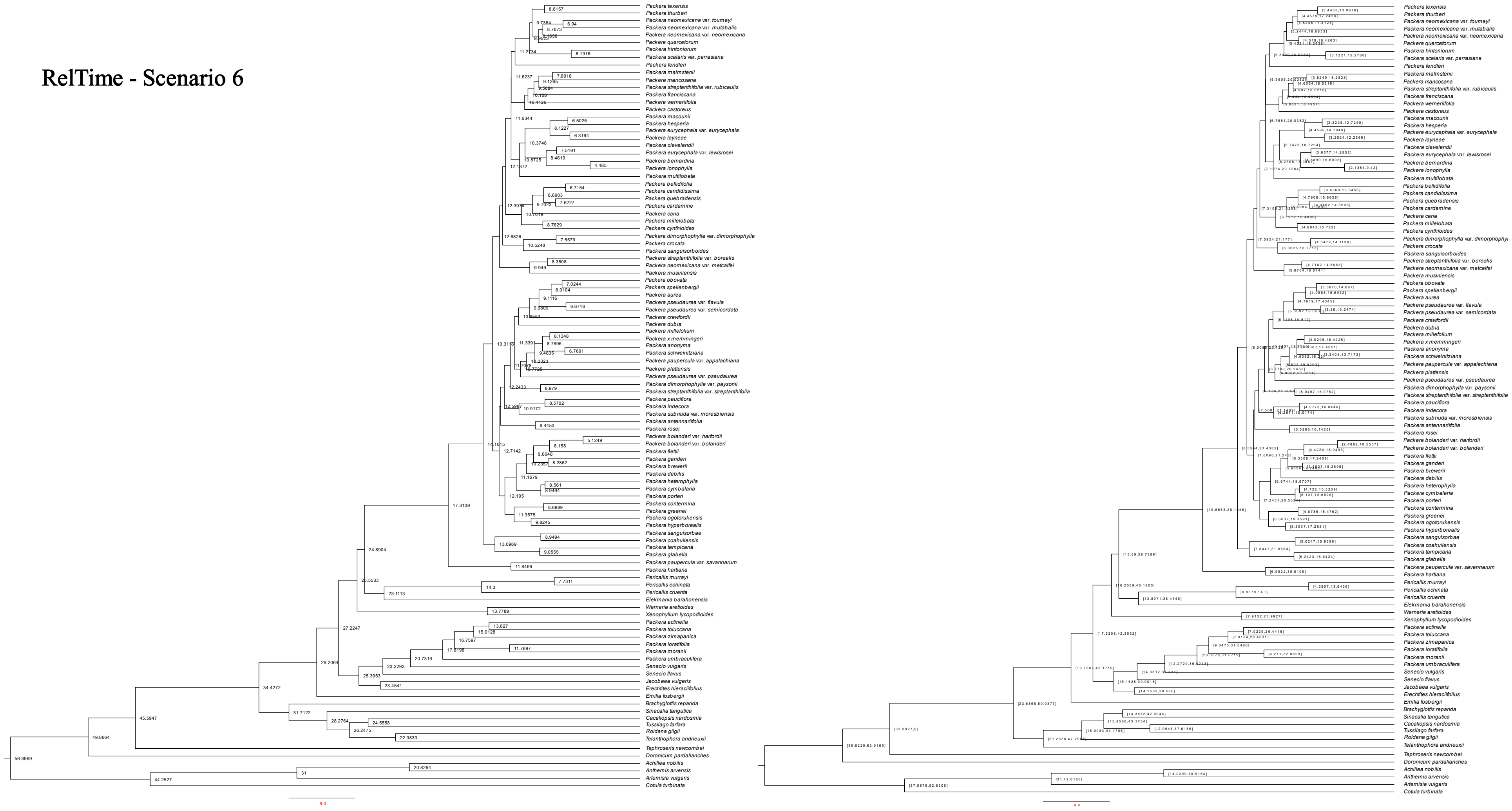

**Supplemental Fig. 19.** Divergence rate estimates of Scenario 6 using RelTime in millions of years. Left tree contains the mean age of that designated node, right tree shows the 95% age range for the designated node.

RelTime - Scenario 7

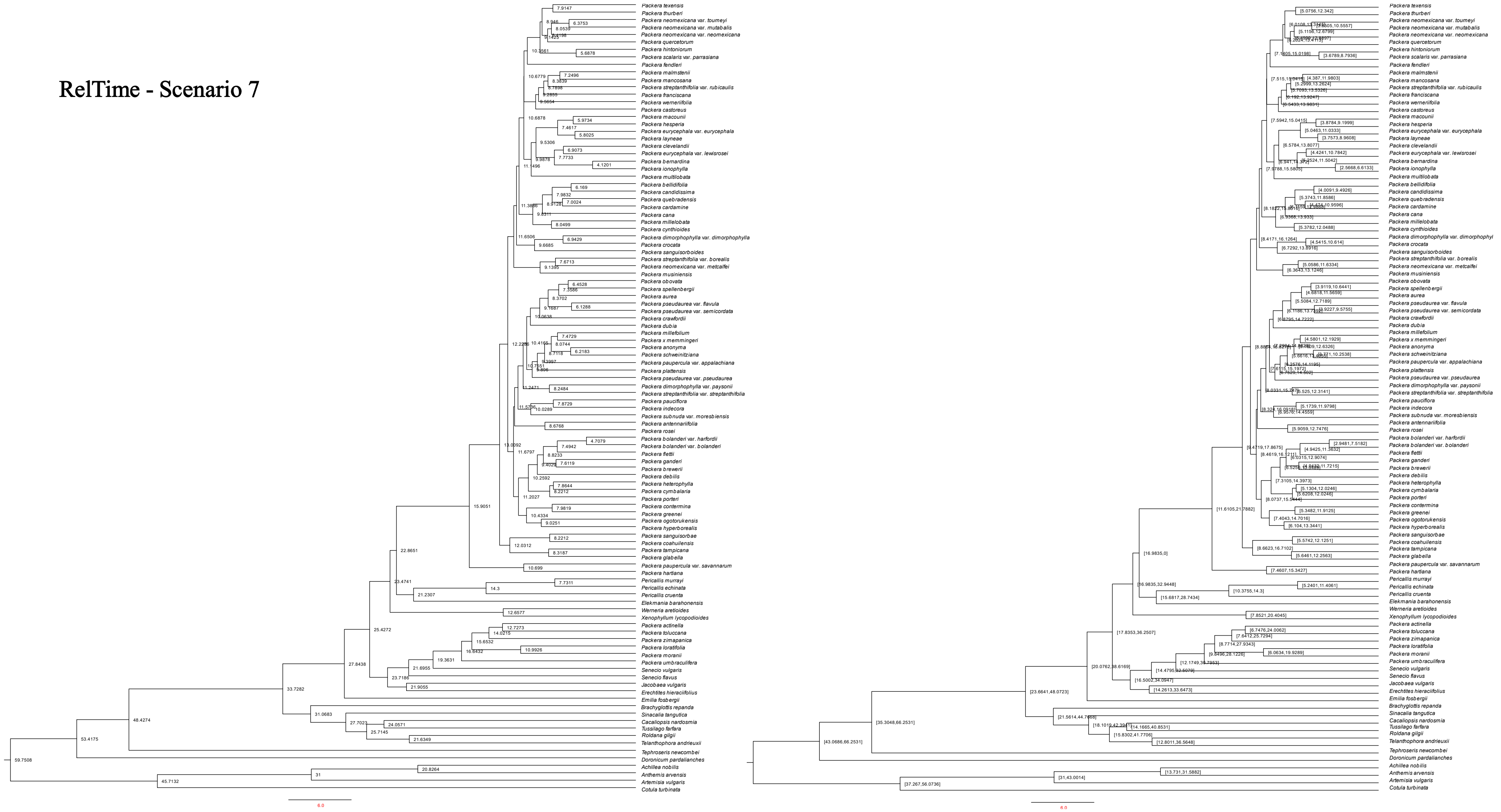

**Supplemental Fig. 20.** Divergence rate estimates of Scenario 7 using RelTime in millions of years. Left tree contains the mean age of that designated node, right tree shows the 95% age range for the designated node.

### RelTime - Scenario 8

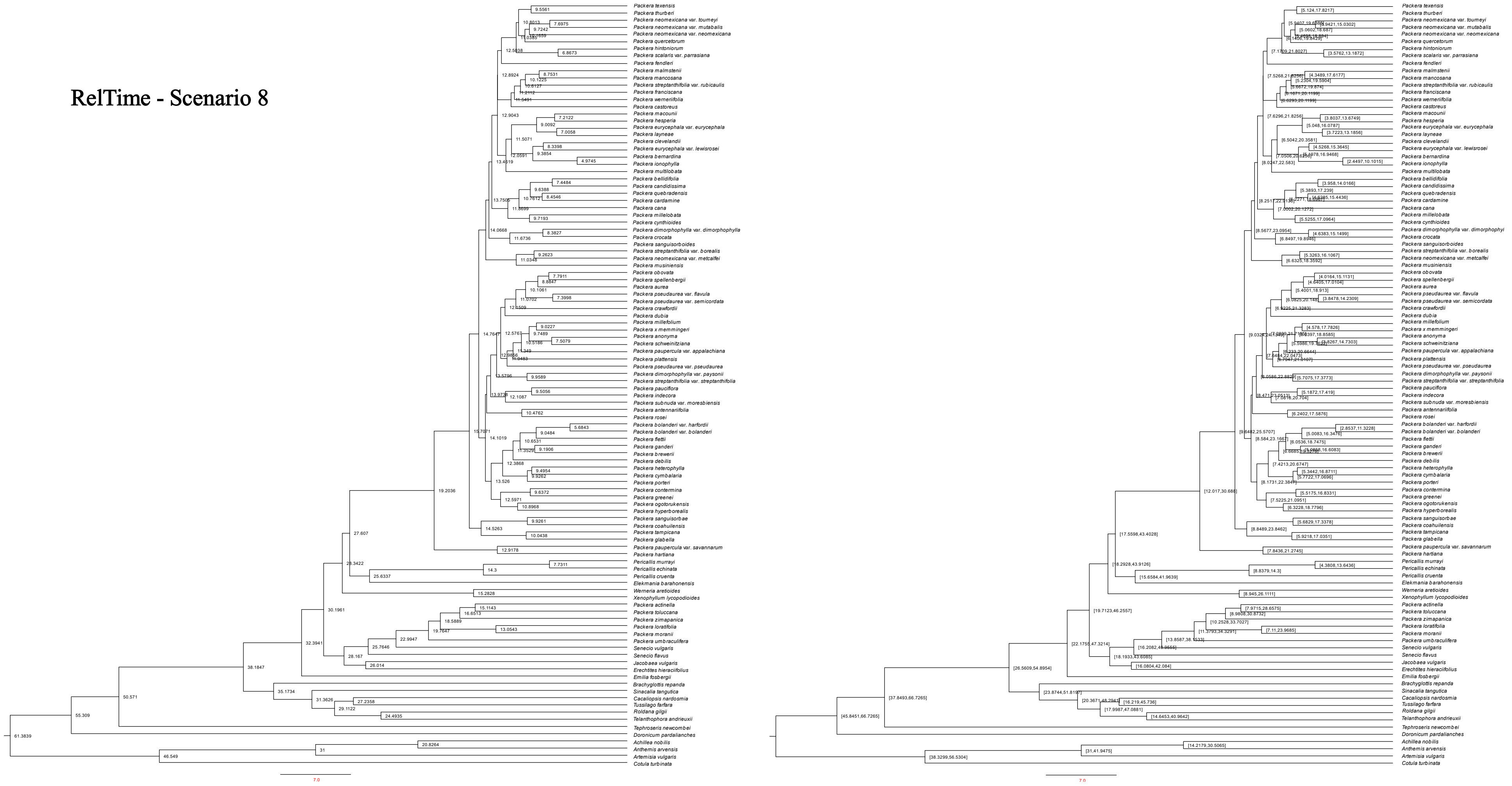

**Supplemental Fig. 21.** Divergence rate estimates of Scenario 8 using RelTime in millions of years. Left tree contains the mean age of that designated node, right tree shows the 95% age range for the designated node.

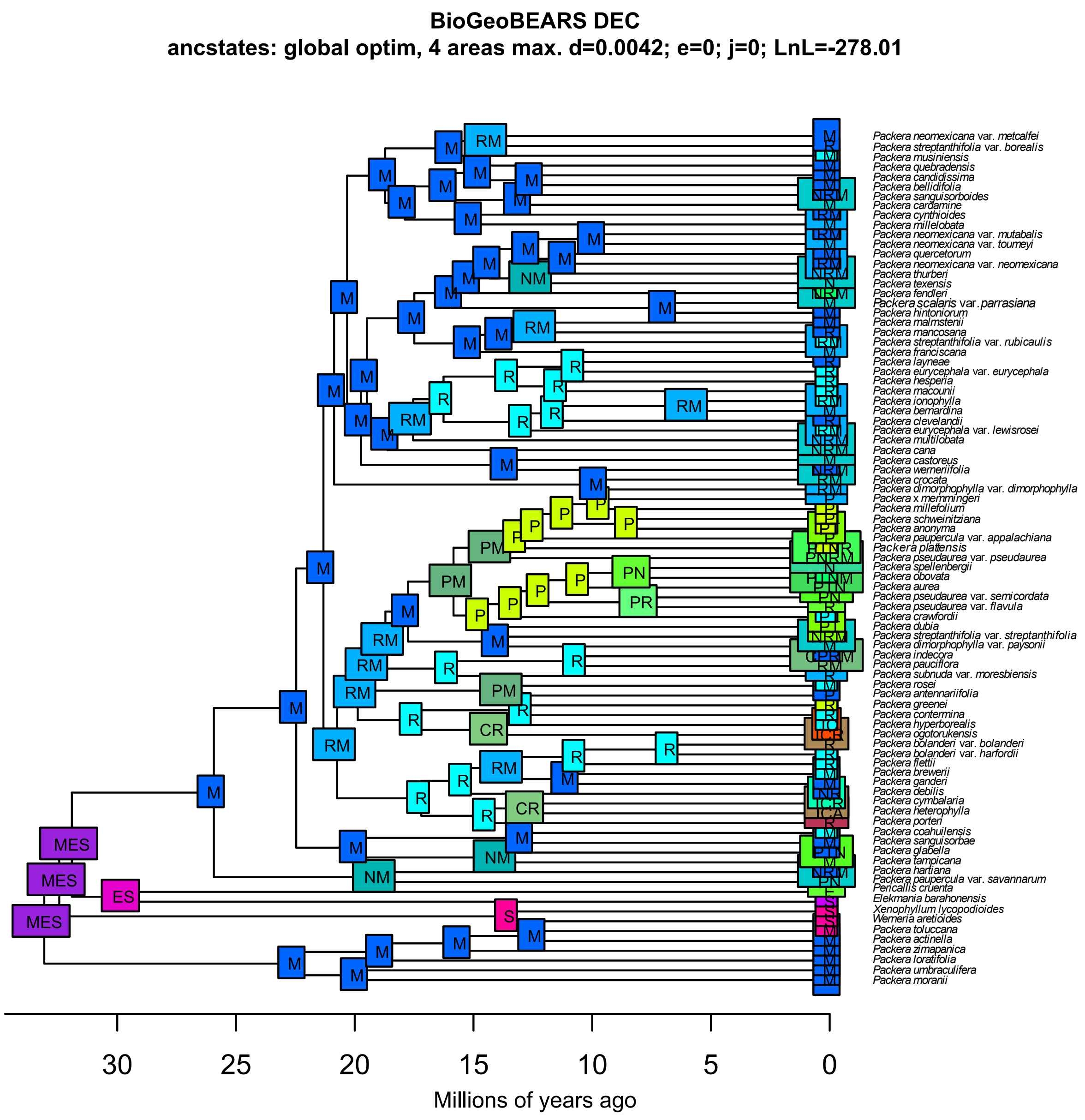


**Supplemental Fig. 22.** BioGeoBEARS results using DEC. Boxes at nodes represent the geographic regions provided in Figure 6.

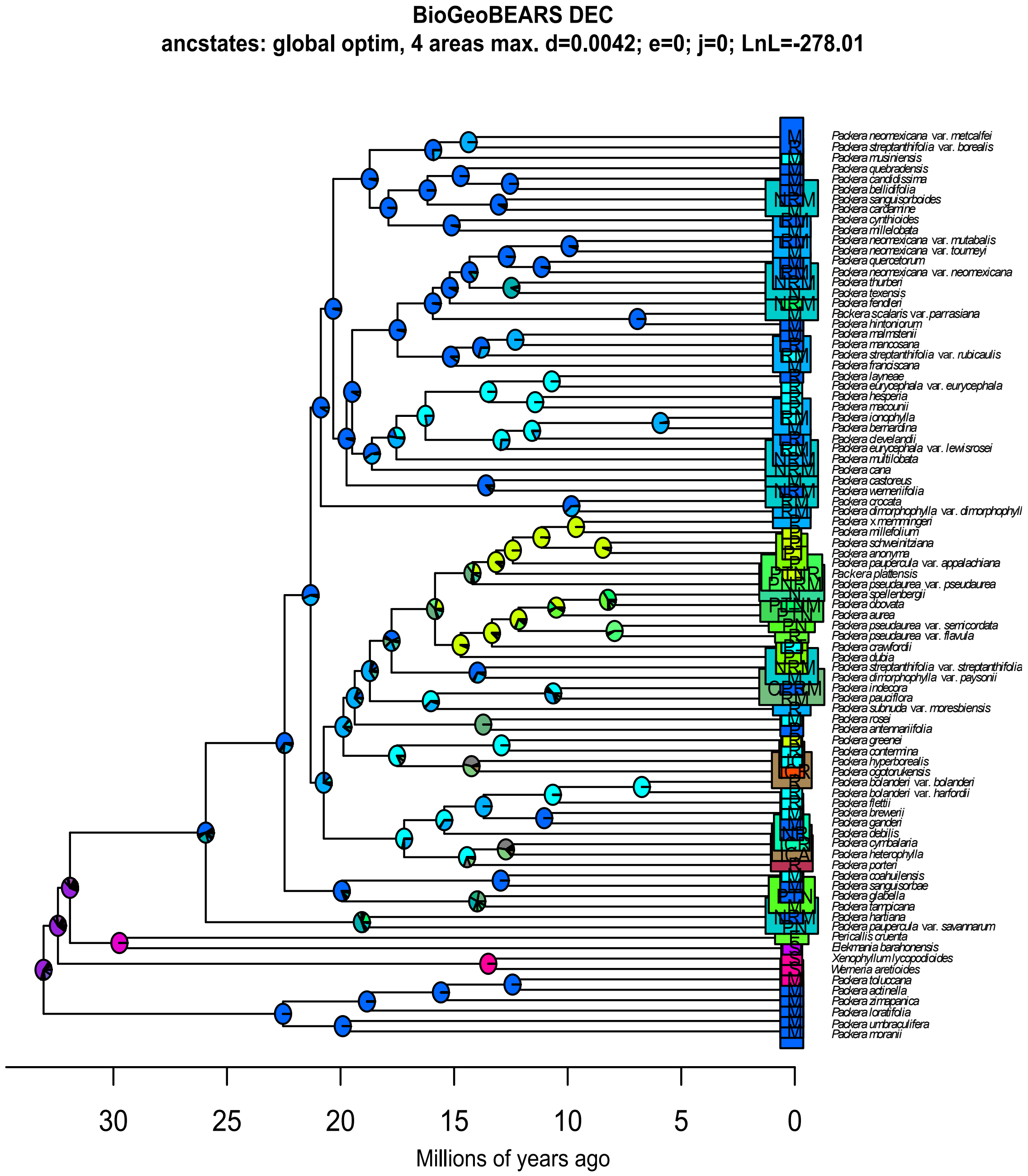


**Supplemental Fig. 23.** BioGeoBEARS results using DEC. Pie charts at node show the probability of geographic regions provided in Figure 6.

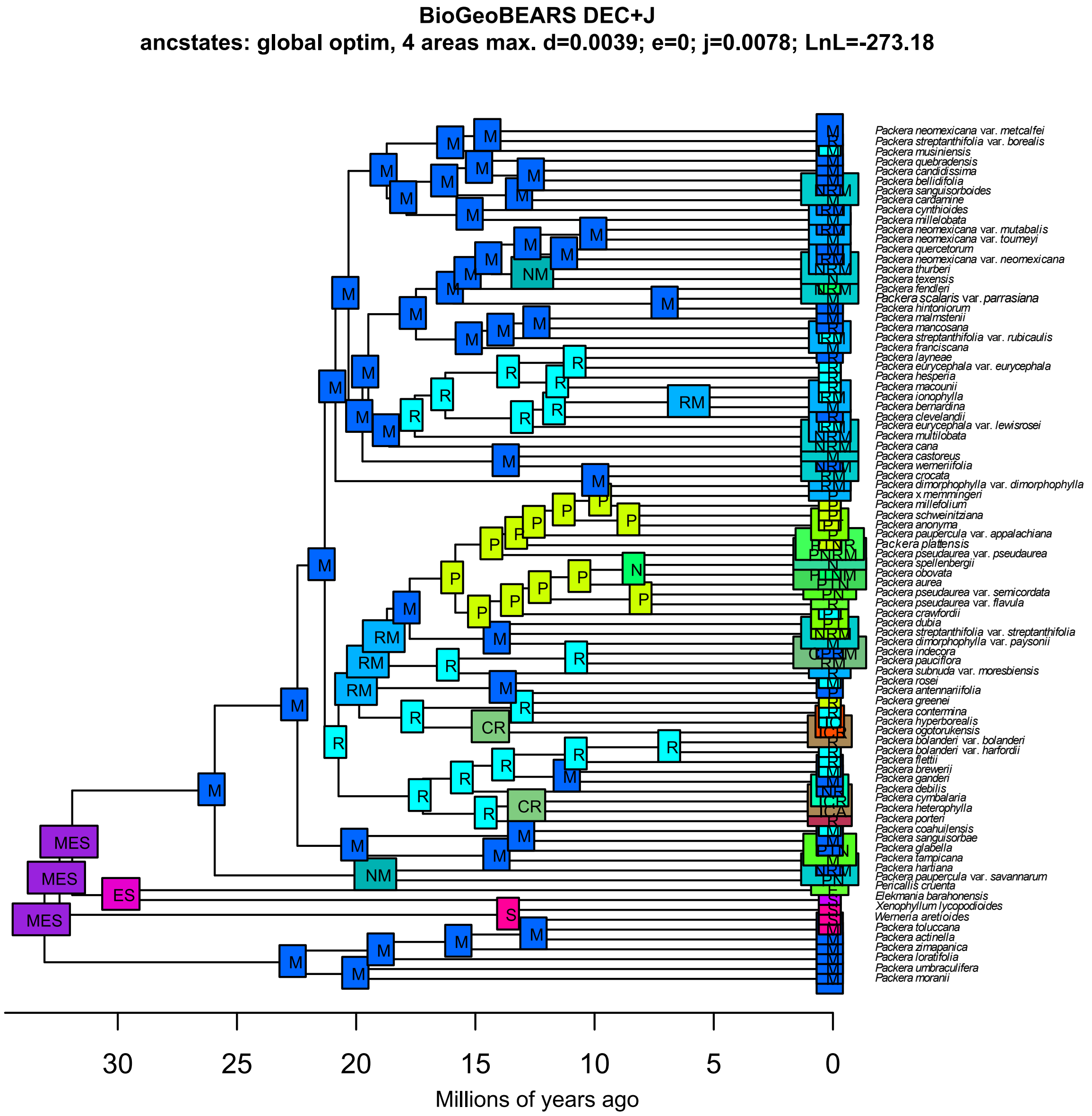


**Supplemental Fig. 24.** BioGeoBEARS results using DEC+*J*. Boxes at nodes represent the geographic regions provided in Figure 6.

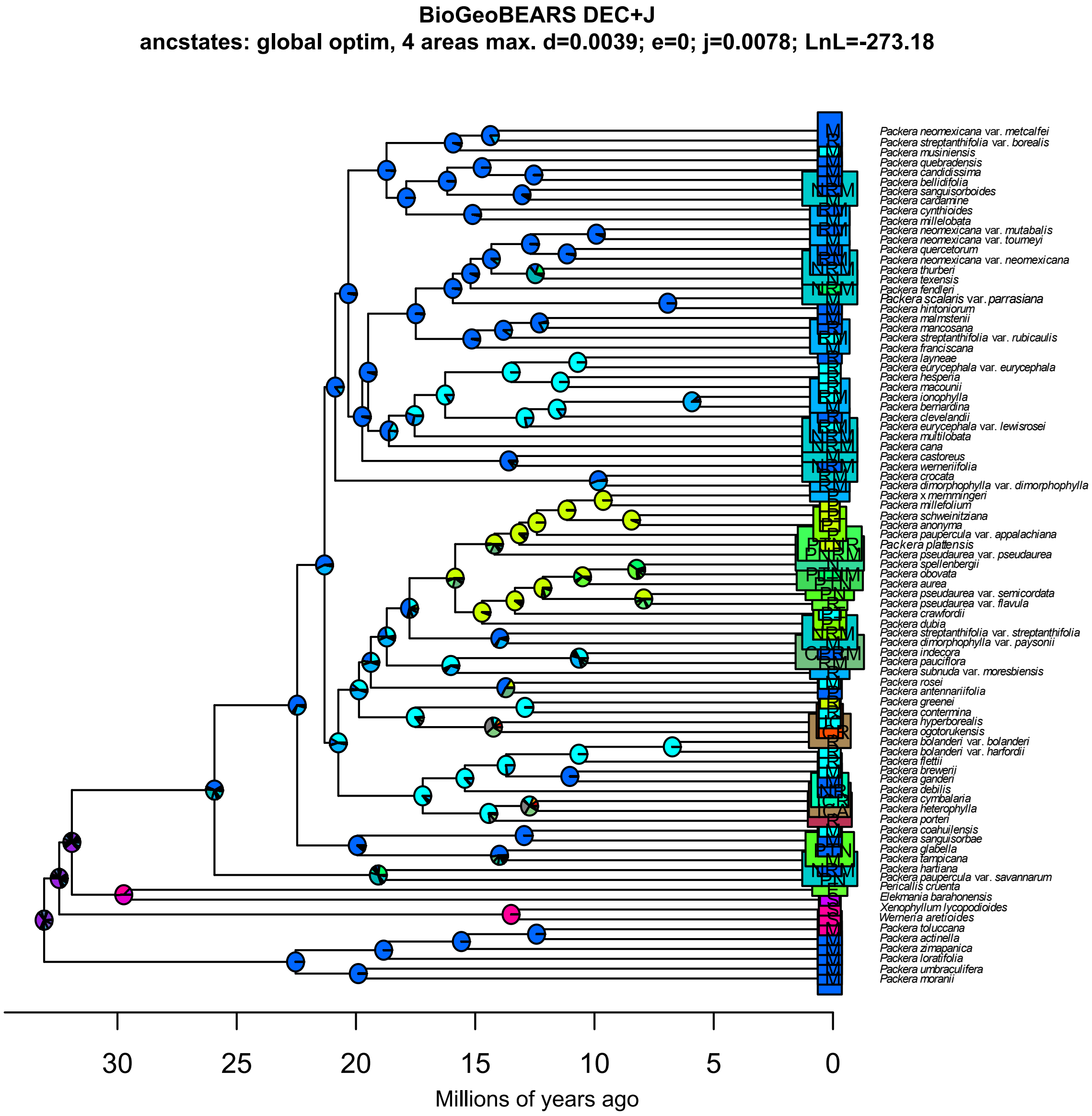


**Supplemental Fig. 25.** BioGeoBEARS results using DEC+*J*. Pie charts at node show the probability of geographic regions provided in Figure 6.

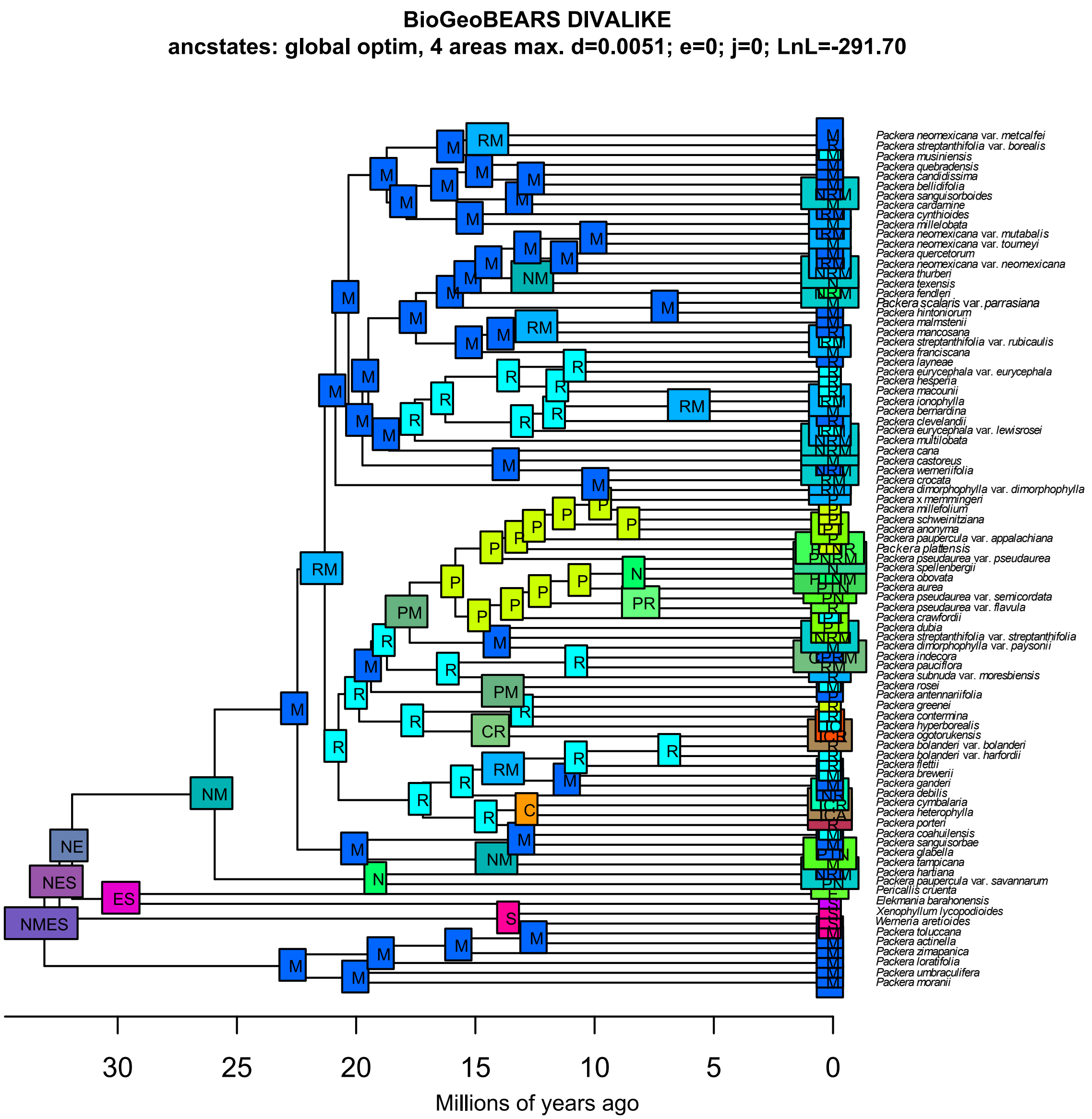


**Supplemental Fig. 26.** BioGeoBEARS results using DIVALIKE. Boxes at nodes represent the geographic regions provided in Figure 6.

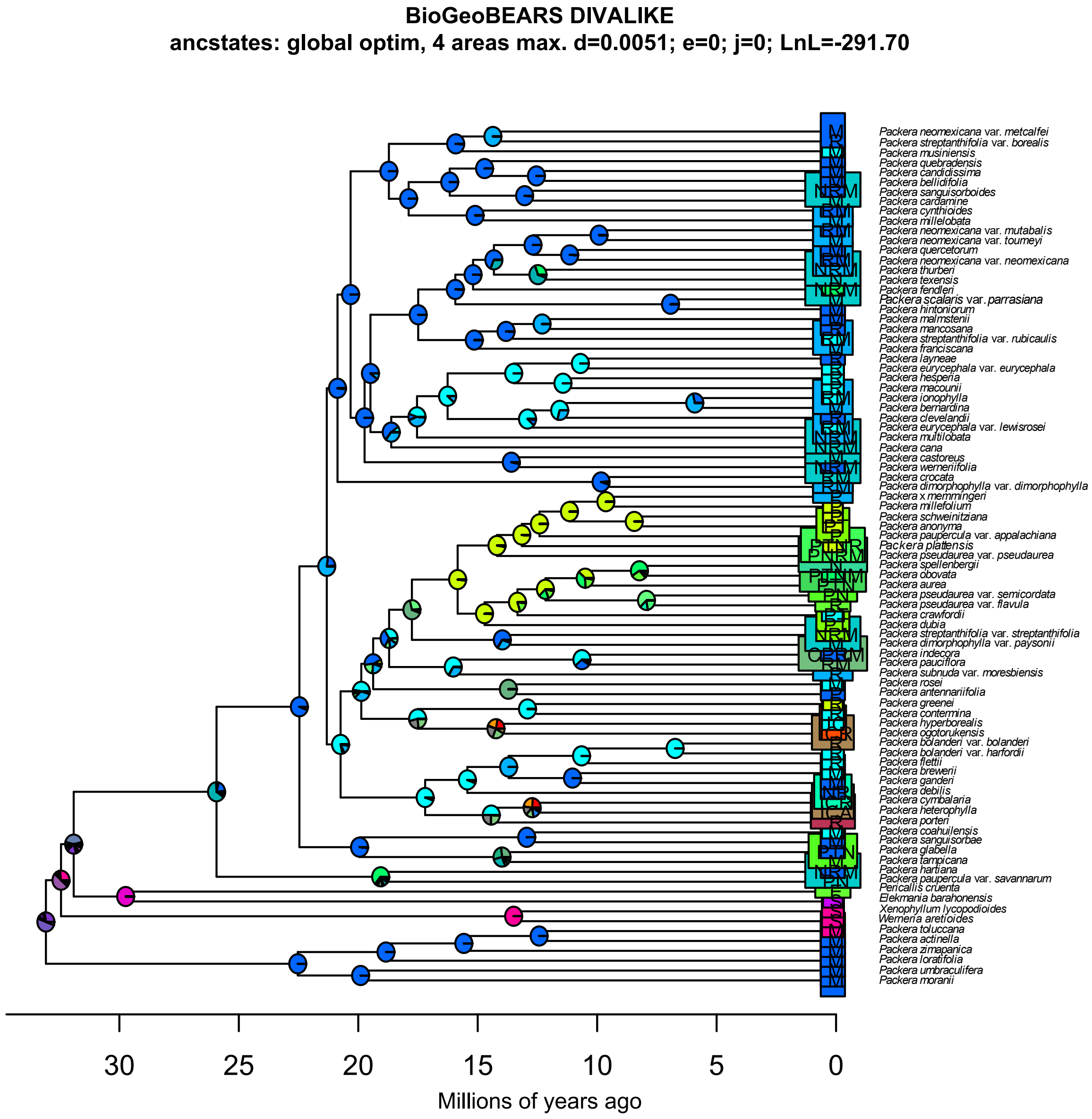


**Supplemental Fig. 27.** BioGeoBEARS results using DIVALIKE. Pie charts at node show the probability of geographic regions provided in Figure 6.

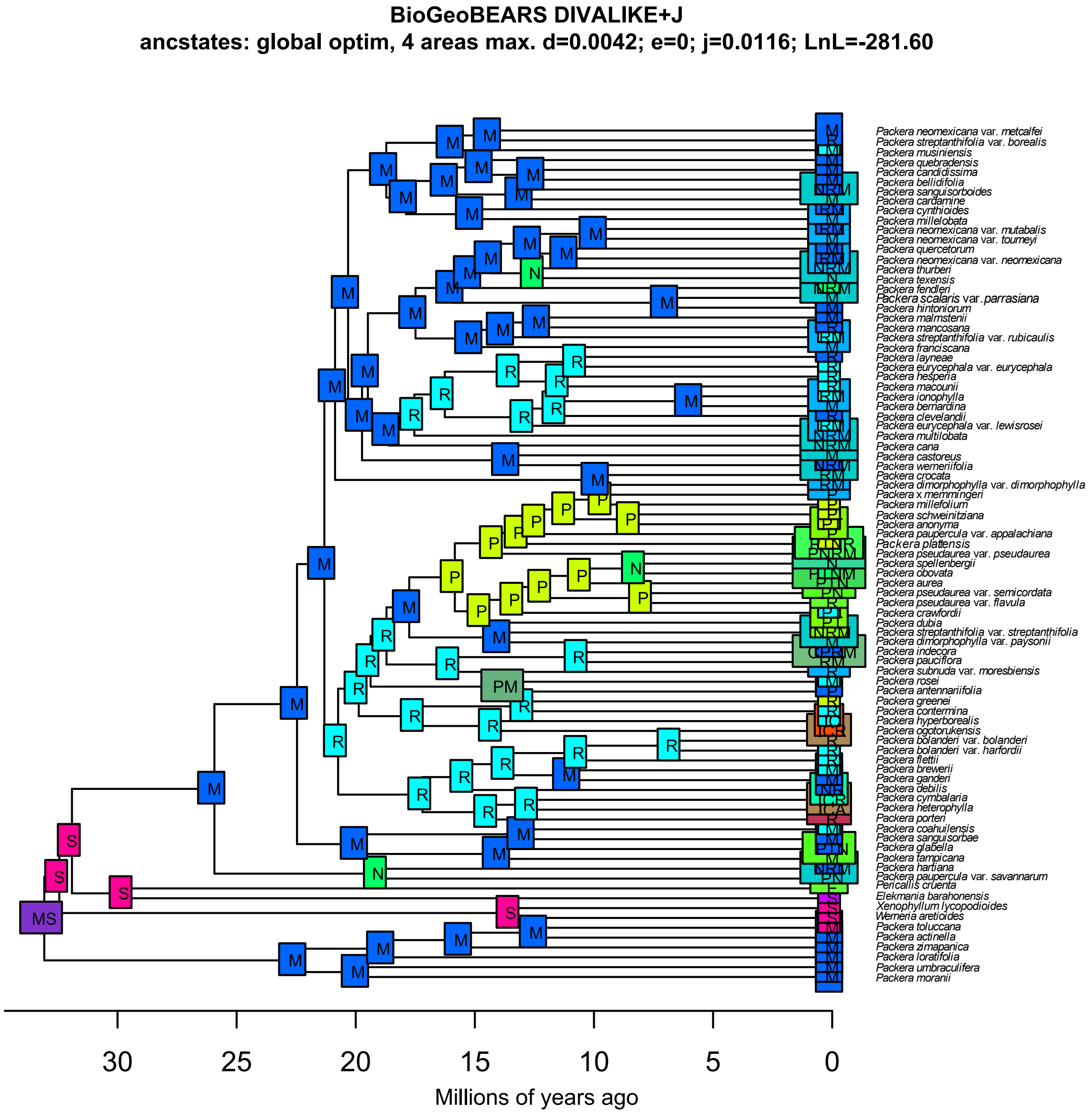


**Supplemental Fig. 28.** BioGeoBEARS results using DIVALIKE+*J*. Boxes at nodes represent the geographic regions provided in Figure 6.

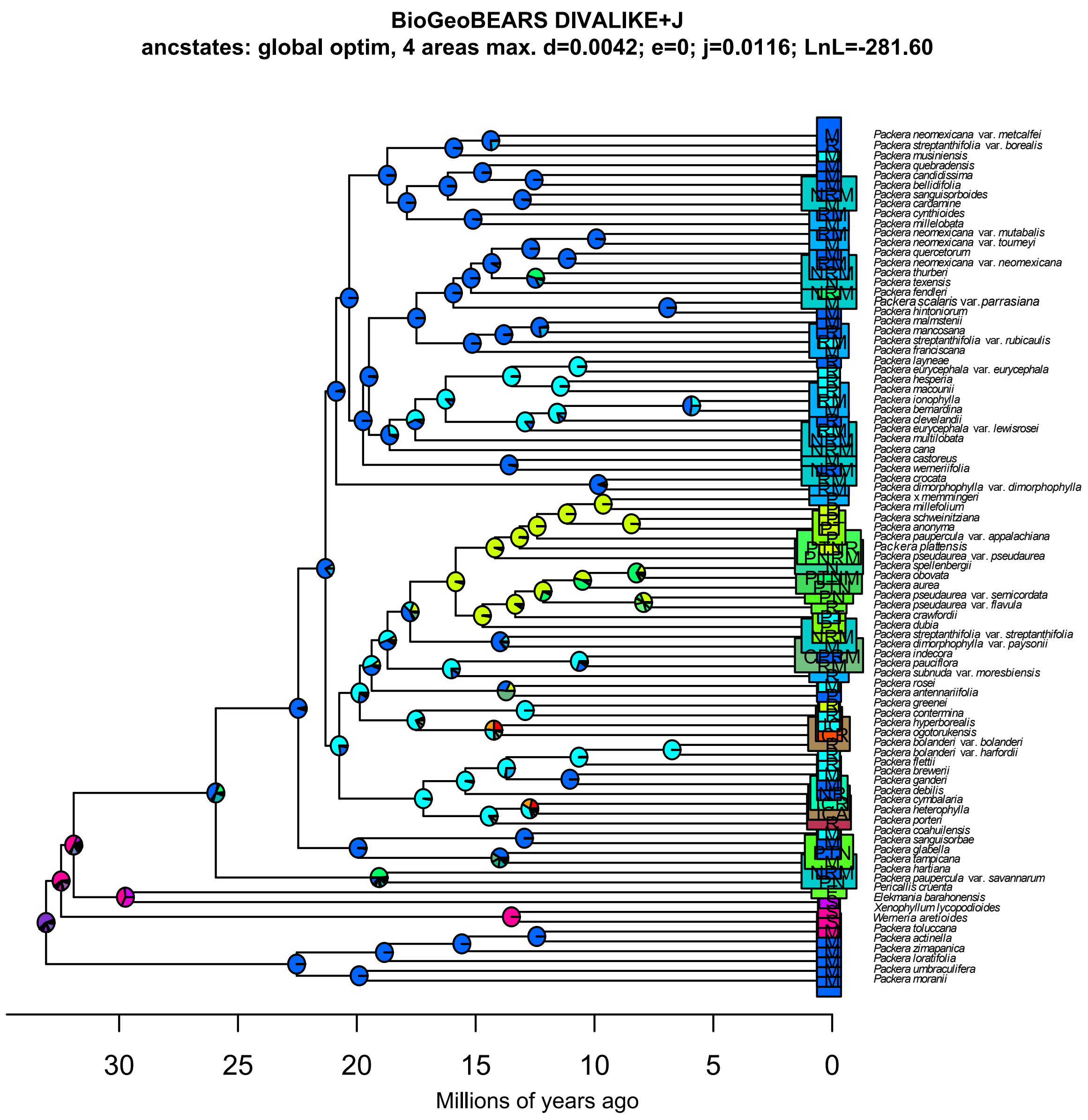


**Supplemental Fig. 29.** BioGeoBEARS results using DIVALIKE+*J*. Pie charts at node show probability of geographic regions provided in Figure 6.

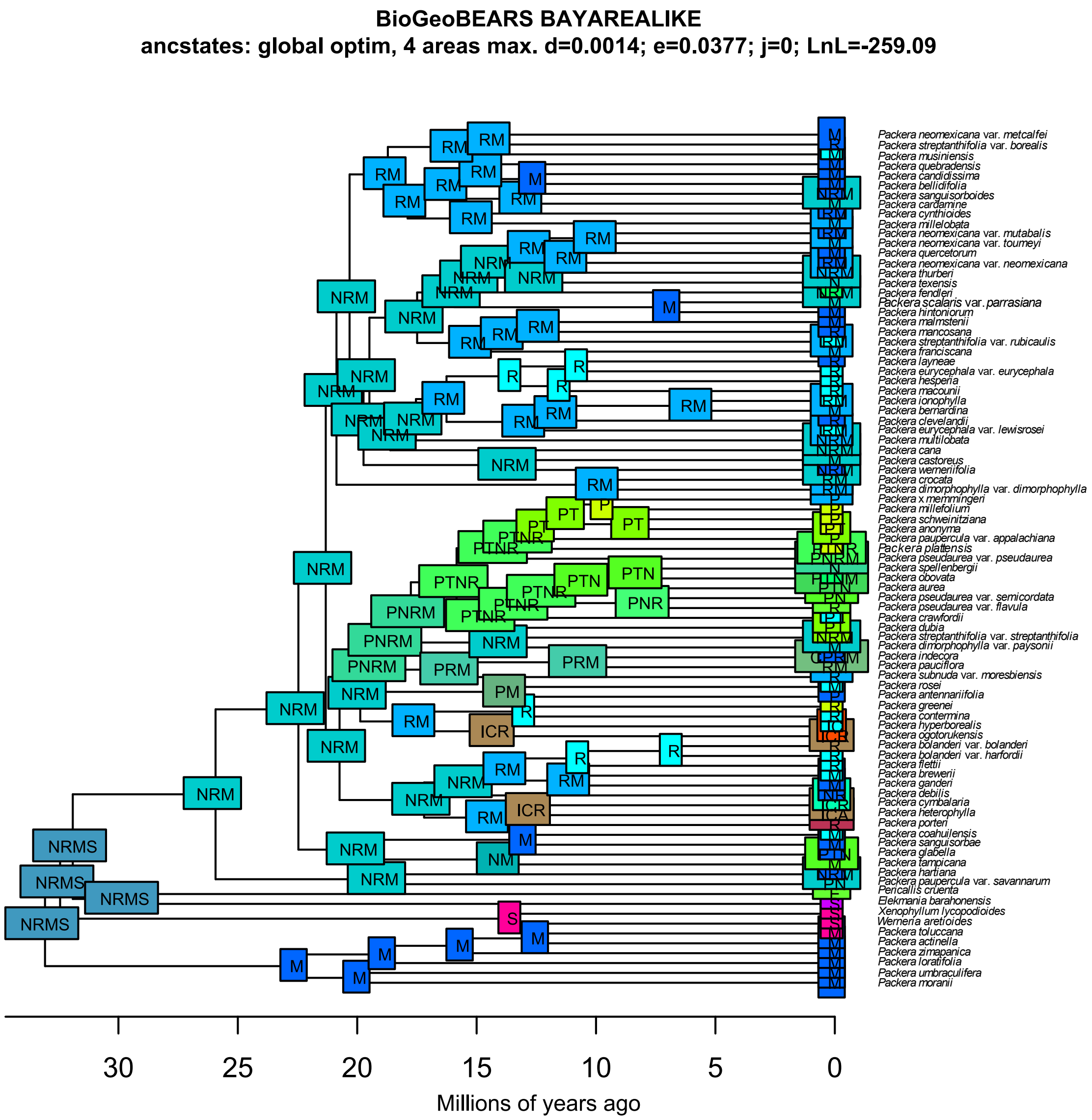


**Supplemental Fig. 30.** BioGeoBEARS results using BAYAREALIKE. Boxes at nodes represent the geographic regions provided in Figure 6.

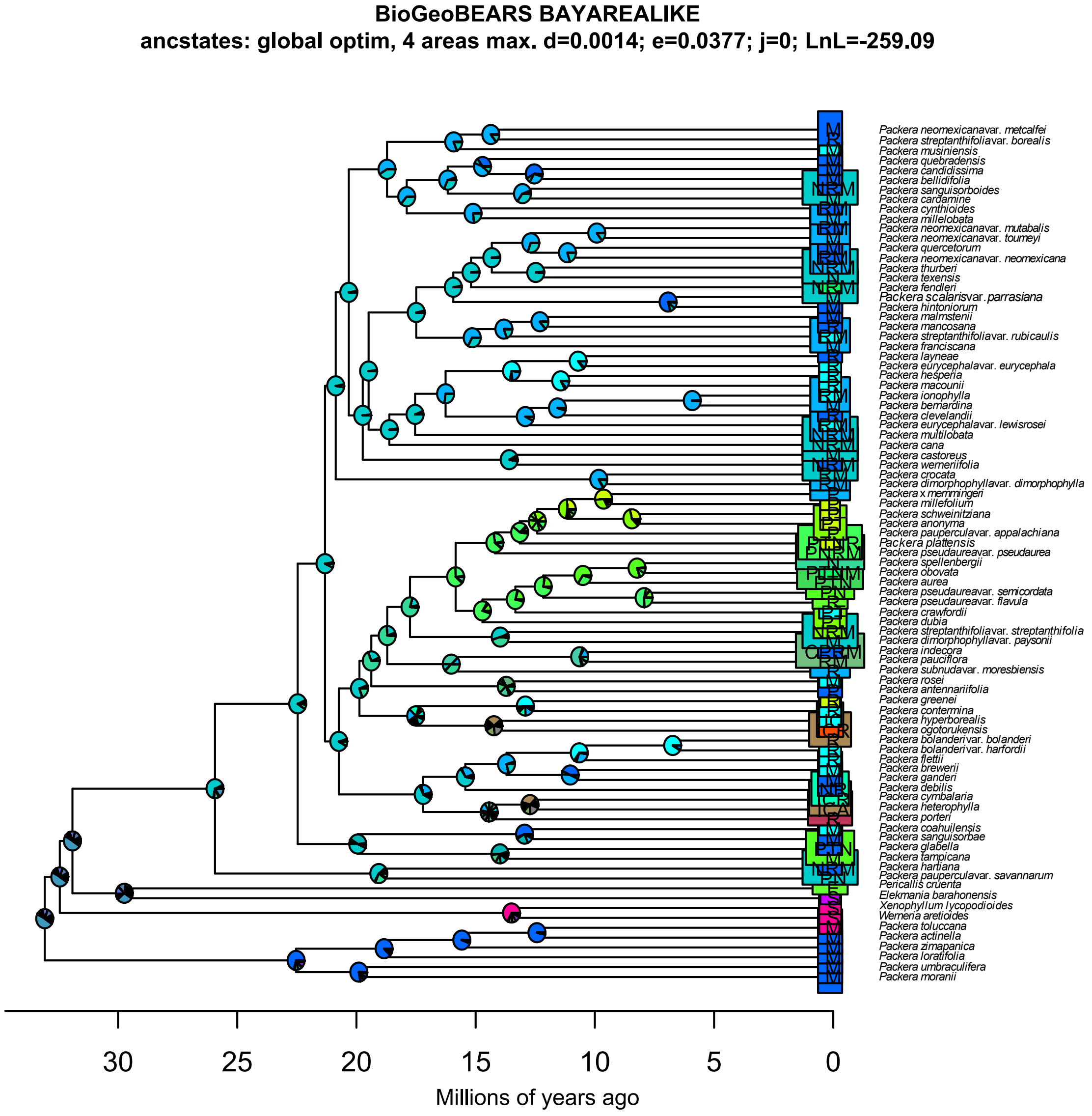


**Supplemental Fig. 31.** BioGeoBEARS results using BAYAREALIKE. Pie charts at node show the probability of geographic regions provided in Figure 6.

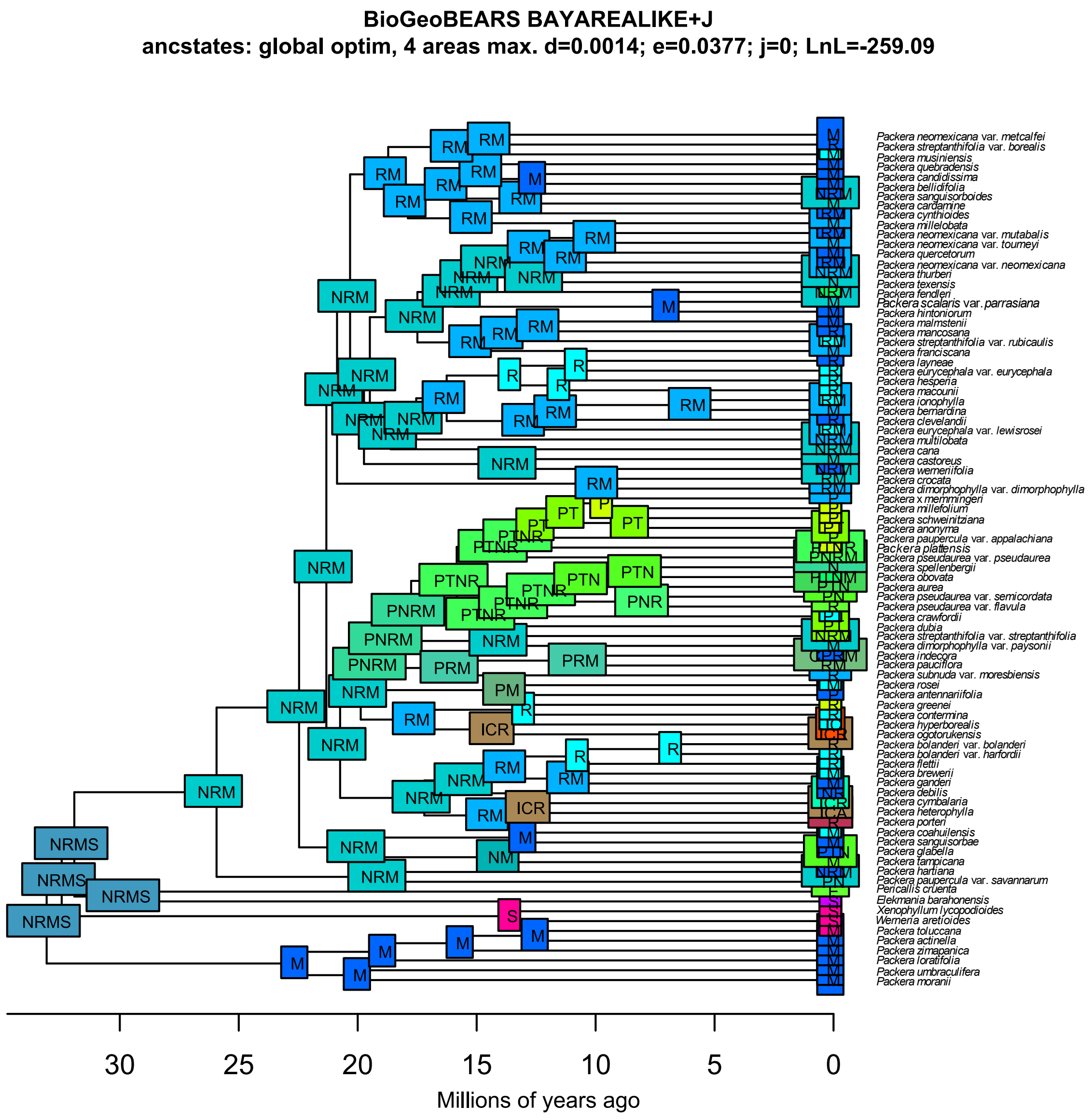


**Supplemental Fig. 32.** BioGeoBEARS results using BAYAREALIKE+*J*. Boxes at nodes represent the geographic regions provided in Figure 6.

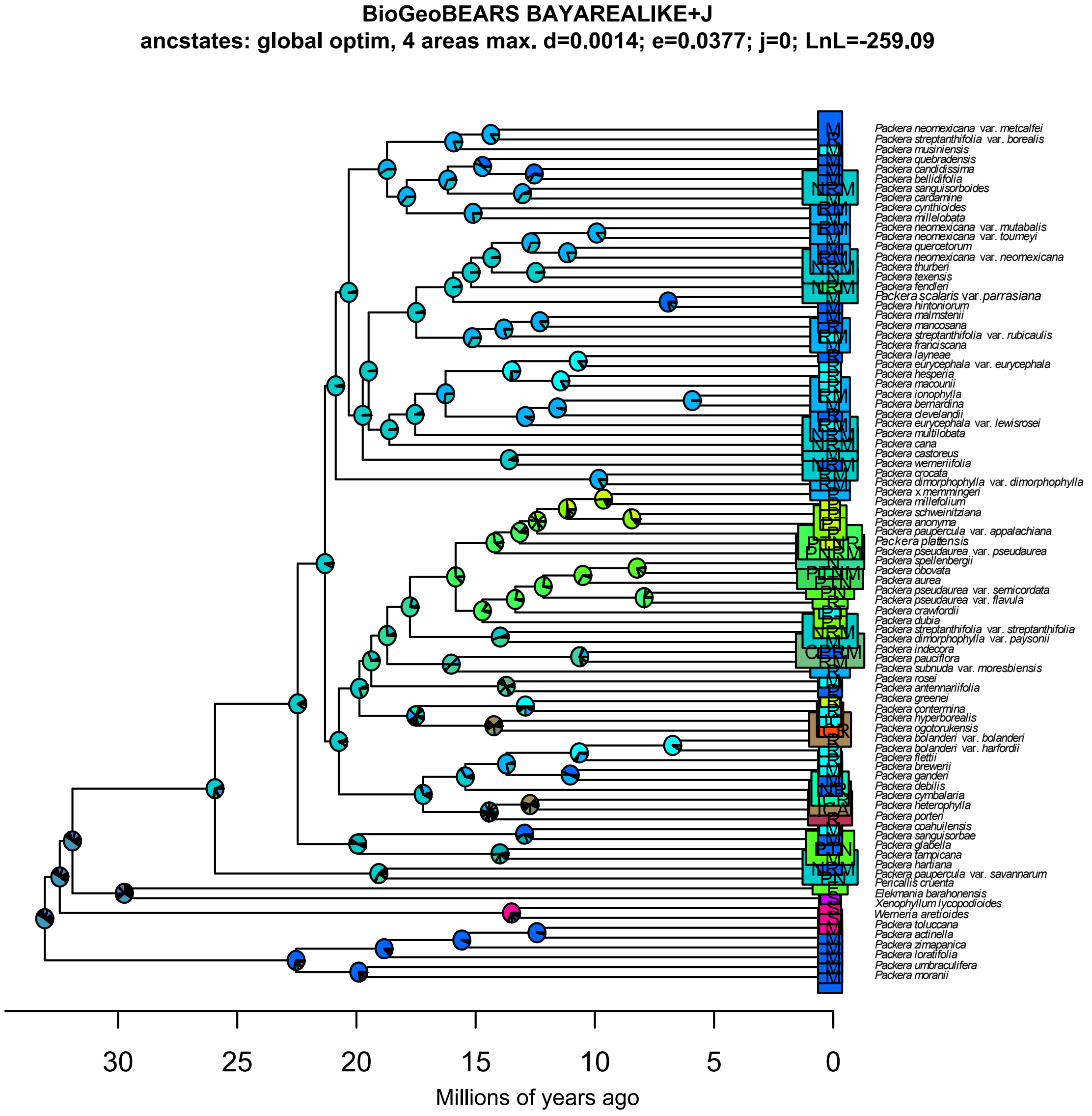


**Supplemental Fig. 33.** BioGeoBEARS results using BAYAREALIKE+*J*. Pie charts at node show the probability of geographic regions provided in Figure 6.
